# Untargeted Metabolomics Shows Alterations in Homocysteine, Lipids and Fatty Acids predicting Memory Decline in Healthy Middle-Aged Individuals

**DOI:** 10.1101/2020.02.23.949537

**Authors:** Ihab Hajjar, Qingpo Cai, Tianwei Yu, Dean P. Jones

**Author notes:** Corresponding Author: Ihab Hajjar, MD, MS.

## Abstract

**INTRODUCTION:** Some aspects of memory start declining in the fifth decade which may be related to systemic metabolic changes. These changes have not been fully identified. This is the first Metabolome-Wide Association Study of the human plasma for the longitudinal change in memory in healthy adults.

**METHODS:** Ultra-high resolution mass spectrometry with liquid chromatography was performed on 207 University employees’ plasma.

**RESULTS:** From 10,201 measured metabolic features, 558 differed between those experiencing change vs no change in memory (False Discovery Rate, FDR< 0.2). Differentially abundant metabolites were observed in lipid and fatty acid metabolism pathways: glycerophospholipid (p=0.0003), fatty acid (p=0.0003) and linoleate (p=0.0003) pathways. Within these pathways, higher homocysteine (OR for memory decline=1.09, FDR=0.19) and lower arachidonic acid (OR=0.97, FDR=0.19), sterol (OR=0.92, FDR=0.02), acetylcholine (OR=0.78, FDR=0.19), carnitine (OR=0.75, FDR=0.19) and linoleic acid (OR=0.74, FDR=0.19) were associated memory decline.

**DISCUSSION:** Altered systemic lipid and fatty acid are linked with early memory decline in middle-aged individuals.

## 1. Introduction

Although there is a wide heterogeneity within the population, there is evidence for progressive declines in memory processes with aging that may start in middle age, ^1,23^ In the some cases, these early declines progress to dementia and Alzheimer’s disease (AD) later in life. Prior evidence supports the existence of multiple metabolic changes or signature for AD.^4-7^ Whether similar or related metabolic changes can be identified in those with early age-related memory changes remain unknown. Identification of such metabolic alterations would offer insight into mechanisms and opportunities for drug development.

Recent advances in mass spectrometry and related computational methods offer the opportunity to assess a large number of metabolic features in small volume biospecimens. ^8^ This high resolution metabolomics (HRM) uses a combination of chromatography coupled to ultrahigh resolution mass spectrometry and advanced computational approaches for spectral feature alignment, peak integration, and feature extraction.^9-11^ With this workflow, HRM is capable of measuring greater than 10,000 unique spectral features with high reproducibility [defined by a characteristic mass-to-charge ratio (m/z), retention time (RT), and intensity]. This untargeted approach has been applied by our team and others in multiple populations and diseases and provides improved capabilities for biomarker discovery, mechanistic understanding of disease and biological phenomena by identifying related chemical pathways and metabolic networks.^12,13^

The human metabolome includes molecules from endogenous sources (e.g., lipids, carbohydrates, nucleotides, amino acids, metabolic intermediates, signaling molecules, small peptides) and exogenous dietary and environmental sources. The expansion of chemical databases such as Metlin (over 240,000 entries), the Kyoto Encyclopedia of Genes and Genomes (KEGG) (over 17,000 entries), or the Human Metabolome Database (HMDB) (over 40,000 entries) has facilitated efforts to identify many features obtained from HRM with very low abundance. To date, our coverage has exceeded 50% of identified metabolites in these databases and due to this wide coverage we are now able to perform metabolome-wide association studies (MWAS) that have detailed coverage of the human metabolome.^14^ Applying this workflow to early changes in memory in middle age has a significant impact on our understanding and potential prevention of age-related memory declines.

We aimed at conducting MWAS for memory changes over 4 years in a group of dementia-free middle aged adults using HRM to identify key metabolic features and pathways involved in early memory changes with aging.

## 2. Methods

### 2.1 Cohort Description

A cohort of healthy employees from Emory University and Georgia Institute of Technology were randomly recruited into the Center for Health Discovery and Well Being cohort as part of the Emory University/Georgia Tech Predictive Health Institute (http://predictivehealth.emory.edu). ^15,16^ Exclusion criteria were a history in the past year of non-accident related hospitalization, severe psychosocial disorder, or addition of new prescription medications to treat a chronic disease (except for changes in anti-hypertensive or anti-diabetic agents), active drug abuse or alcoholism, a current active malignant neoplasm, uncontrolled or poorly controlled autoimmune, cardiovascular, endocrine, gastrointestinal, hematologic, infectious, inflammatory, musculoskeletal, neurological, psychiatric, or respiratory disease, and any acute illness in the 2 weeks before baseline studies.

### 2.2 Cognitive assessment and derivation of memory scores

Commonly employed versions of neuropsychological measures were administered via computer at baseline, and then yearly for a total of 4 times, using software developed by Aharonson and colleagues.^17,18^ Cognitive tests included memory delayed recall, memory recognition, visual spatial learning, spatial short term memory, pattern recall, delayed pattern recall and recognition of pattern, executive function test, mental flexibility, digit symbol substitution test, forward and backward digit span, symbol spotting, and focused and sustained attention (computerized score:0-100% correct adjusted for skill levels). Cognitive scores for cognitive domains were derived using principal component analysis with Varimax (orthogonal) rotation and Kaiser Normalization to perform the exploratory factor analysis and then performed a confirmatory factor analysis by exploring the correlations and model fit of the derived factor-saved scores as reported previously. ^19^ The memory score (unit less) was differentially loaded by the delayed recall, pattern delayed recall and visual-spatial memory (results of the factor analysis are shown in the online supplement of reference^19^). Test scores were then divided into a binomial variables: low vs high performance during the follow-up relative to baseline performance if a participant’s score was below (or above) the lowest quartile of the corresponding score at baseline. We restricted our analysis to the memory domain since it may start to change by the 5^th^ decade of age and included those with at least 2 cognitive evaluations during the study period.

### 2.3 High Resolution metabolomics

Our analysis was performed using an ultra-high performance liquid chromatography (UPLC) system coupled with ultra-high-resolution mass spectrometry (MS) at Emory University Biomarker laboratory as described previously. ^20,21^ Samples were analyzed using the Reverse Phase (C18) chromatography with positive electrospray ionization (ESI). This method detects >10,000 ions with 49% KEGG matches and provides accurate mass matches to 55% of the human metabolome KEGG database.^14^ Plasma samples were extracted with acetonitrile, and then analyzed along with NIST 1950-calibrated reference pooled human plasma preceding and following each block according to standard operating procedures. Samples were analyzed in triplicate and a feature was included only if it was detected on at least two out of three technical replicates, and features with greater than median 50% CV for technical replicates were removed from analyses. The triplicate measurements from each subject/visit were merged by taking the mean of the non-zero values of each feature. Pooled reference plasma was run prior to and after each batch for quality control and quality assurance.

### 2.4 Bioinformatic analyses

An adaptive processing software package (apLCMS, http://web1.sph.emory.edu/apLCMS/) was used for peak extraction and quantification of ion intensities.^11^ This software provided feature tables containing m/z values, retention time, and integrated ion intensity for each m/z feature, obtained through five major processing steps: (1) noise filter, (2) peak identification, (3) retention time correction, (4) m/z peak alignment across multiple spectra, and (5) reanalysis to capture peaks originally missed because of weak signal relative to the signal to noise filter. We used xMSanalyzer (http://sourceforge.net/projects/xmsanalyzer/) to enhance the feature detection process by performing systematic data re-extraction, statistical filtering, and data merger to enhance quality of data extraction.^22^ xMSanalyzer also provided batch normalization using ComBat ^23^ to control for batch effects. Data then underwent log2 transformation to reduce heteroscedasticity and normalize results. Obtained dataset was then used for the statistical analyses.

### 2.5 Statistical analysis

We used the baseline metabolic features as our independent variables and the longitudinal memory measure as the dependent variable (discrete measure 0, 1 as defined above). Since we have longitudinal data with repeated measures, a Generalized Estimating Equation (GEE) was used to estimate the coefficient of associations. The analysis was done one metabolic feature at a time. Specially, let *y*_*ij*_ denote the value for memory score, *x*_*ij*_ denote the metabolic feature under study at baseline, and*z*_*ij*1_, …, *z*_*ij*p_ denote the value for *p* confounding variables (demographic and clinical factors) for subject *i* at visit *j.* In this analysis, *y*_*ij*_ is binary with *y*_*ij*_ = 1 indicating a decline relative to baseline performance. The model is:

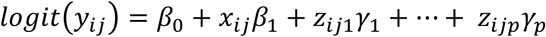

Where 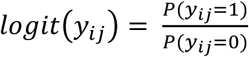 with p (*y*_*ij*_ = 1) being the probability for *y*_*ij*_ = 1. An exchangeable correlation structure is specified to estimate the coefficients in the GEE model.

Each metabolite is analyzed individually to find its marginal effect on the outcome, while adjusting for age, gender, race, years of education, hypertension/antihypertensive therapy, use of statin, mean systolic blood pressure, BMI, and diabetes. For each metabolite, we obtained the coefficient estimation 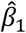 and its associated p-value from the GEE models. We then corrected the p-values from all metabolic features for multiple testing using False Discovery Rate (FDR) ^24^. We choose the significant metabolites by controlling the FDR at 0.2 (FDR<=0.2).

### 2.6 Pathway analysis

To study the metabolic pathways associated with the memory outcome, we used pathway analysis using Mummichog ^25^, which matches metabolic features to adduct ions of metabolites based on m/z, and selects pathways that are over-represented by the selected features. This approach predicts directly from mass spectrometry data without a priori identification of metabolites by unifying network analysis and metabolite prediction under the same framework. This process minimizes ambiguity, uses exiting knowledge from KEGG, UCSD Recon1, and Edinburgh human metabolic network, and has been validated by activation experiments, gene expression and metabolite identification methods reported previously (agreement ranges 80-97%). ^25^ We then calculated the odds ratio for a doubling level for each metabolite within the significant pathways. The interpretation for OR is that if the metabolite level doubles, OR gives us the 4-year cumulative odds of developing cognitive decline (compared to baseline) in the high vs low levels of the metabolite.

## 3. Results

### 3.1 Sample Characteristics

The main characteristics of the analytical sample (n=207, 28% had memory decline) are shown in **Table 1**. The mean derived memory score was 65 (SD=16, max 80) and the mean yearly decline in the sample was 0.05/year. There was no difference in yearly memory change by sex (p=0.22) but there were steeper declines in the non-white participants (0.001 in Whites vs 0.2 in non-Whites, p=0.02) and in those with higher BMI at baseline (β=0.02, p=0.01).

**Table 1:** Key clinical characteristics of the selected sample for the MWAS (N=207)

| **Characteristic** | **Mean or Count (standard deviation or %)** |
| --- | --- |
| Age in years, mean standard deviation (SD) | 50.5 (10.0) |
| Education in years, mean (SD) | 18.8 (4.4) |
| Body Mass Index, mean (SD) | 27.3 (5.7) |
| Systolic Blood Pressure in mm Hg, mean (SD) | 120.7 (17.3) |
| Number of women, n (%) | 136 (65.7%) |
| Number of Whites, n (%) | 157 (75.8%) |
| Number of African Americans, n (%) | 38(18.4%) |
| Number of those receiving anti-hypertensive medications, n (%) | 41 (19.8%) |
| Number of Diabetics, n (%) | 10 (4.8%) |
| Number of those receiving statin medications, n (%) | 41 (19.8%) |

### 3.2 Metabolomic-Wide-Association Study (MWAS), pathways, and features

Mass spectral data from the sample yielded 10,201 ions which were used in our binomial regression MWAS analyses. Of those, 1483 features were associated with memory change, after adjusting for covariate (age, sex, race, education, BMI, SBP, statins, DM and treatment for DM/HTN). After FDR correction (0.2), 558 features were significantly linked to memory change. These results are shown in the Manhattan Plot (–log P versus m/z), **Figure 1**. The full detailed results of the MWAS are also provided in a supplemental ***Table S1***.

**Figure 1:**
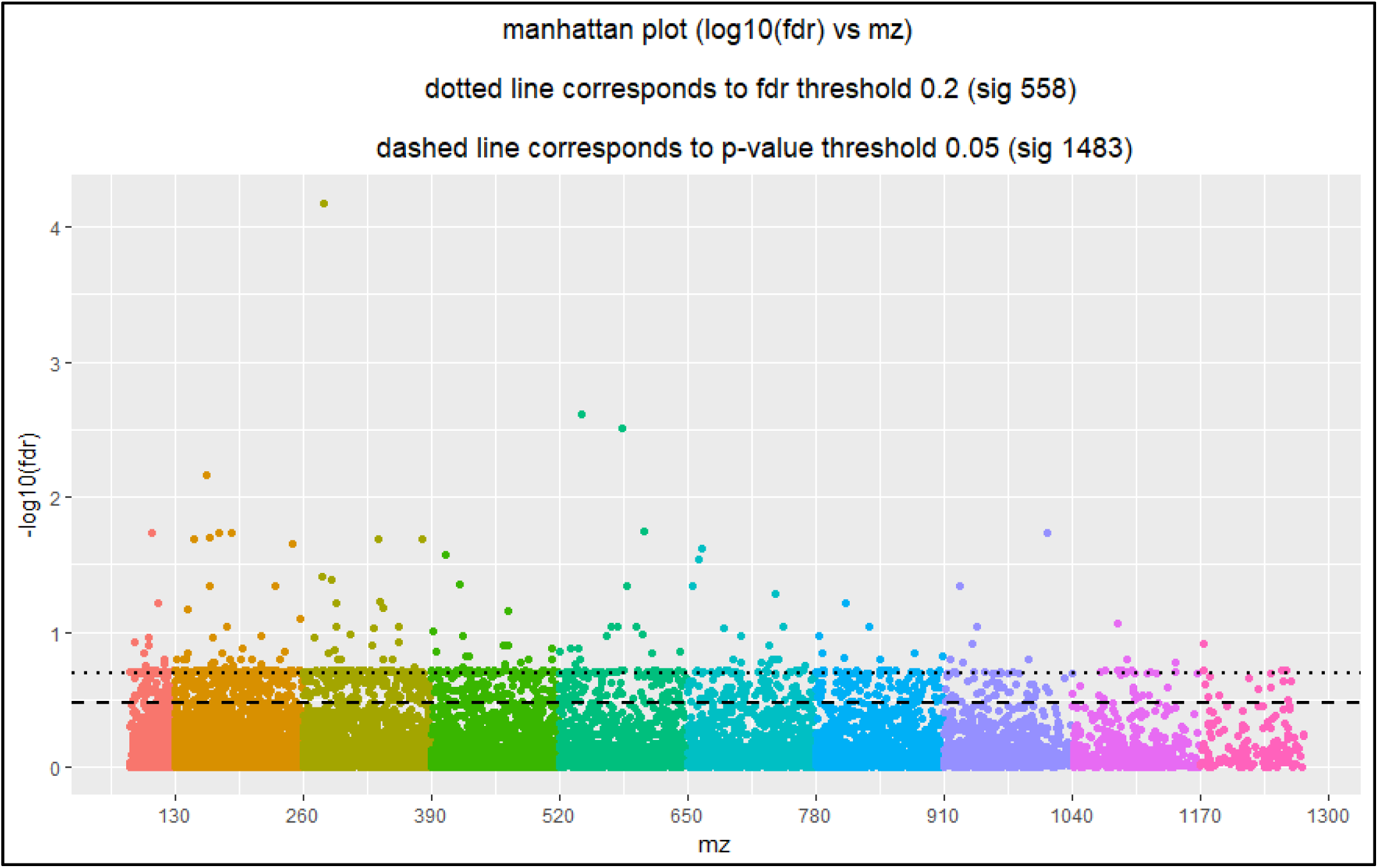
Visualization of the Metabolome-Wide-Association Study (MWAS) for the risk of having memory decline in middle-age adults.

Pathway analysis was subsequently conducted using Mummichog, which matches metabolic features to adduct ions of known metabolites based on m/z in metabolic databases and selects pathways that are over-represented by the selected features. This analysis resulted in identifying a set of pathways that included the significant metabolites showing differential abundance between those with or without memory change. The top pathways that include differentially abundant metabolites are shown in **Figure 2** and included Glycerophospholipid (GCP, P=0.0003), fatty acid (FA, p=0.0003) and linoleate (p=0.0003) metabolism. Within these 3 pathways, we identified 31 features that remained significant after FDR corrections and were matched to an identifiable metabolite. Results for the key significant features within the top 3 pathways along with their KEGG ID and related odds ratio are shown in **Table 2**.

**Table 2:** Results of MWAS and pathway analyses for the top 3 metabolic pathways significantly different between those with vs without memory change (decline) with associated M/Z ratio, retention time, odds ratio (<1 implies protective effect) with the 95% Confidence Interval (CI), Raw p-value and False Discovery Rate (FDR<=0.2 reflects metabolome wide significance with multitests adjustments)

| KEGG.id | Metabolite | M/Z | Retention time | OR | 95% CI, Lower limit | 95% CI, Upper limit | Raw p-value | FDR |
| --- | --- | --- | --- | --- | --- | --- | --- | --- |
|  | **Glycerophospholipid metabolism Pathway** |  |  |  |  |  |  | **0.000329** |
| C06459 | *N-Trimethyl-2-aminoethylphosphonate* | 169.09 | 464.27 | 0.706 | 0.003 | 0.88 | 0.001999 | 0.17 |
| C00137 | *myo-Inositol* | 203.05 | 74.30 | 0.708 | 0.004 | 0.89 | 0.002901 | 0.19 |
| CE0520 | *(R)-glycerol 1-acetate* | 203.05 | 74.30 | 0.708 | 0.004 | 0.89 | 0.002901 | 0.19 |
| C01582 | *Galactose* | 203.05 | 74.30 | 0.708 | 0.004 | 0.89 | 0.002901 | 0.19 |
| C06427 | *alpha-Linolenic acid* | 279.23 | 28.19 | 0.739 | 0.594 | 0.92 | 0.00663 | 0.20 |
| C01595 | *Linoleic acid* | 281.25 | 33.78 | 0.743 | 0.007 | 0.91 | 0.004619 | 0.19 |
| C06893 | *2-Deoxy-5-keto-D-gluconic acid 6-phosphate* | 327.01 | 550.4<br>4 | 0.75<br>8 | 0.01<br>0 | 0.93 | 0.006487 | 0.20 |
| C01996 | *Acetylcholine* | 146.12 | 38.69 | 0.77<br>9 | 0.01<br>7 | 0.94 | 0.010772 | 0.20 |
| C00307 | *Cytidine 5'-diphosphocholine* | 557.10 | 110.9<br>6 | 0.80<br>2 | 0.01<br>6 | 0.95 | 0.009606 | 0.20 |
| C00114 | *Choline* | 104.11 | 78.75 | 0.80<br>6 | 0.01<br>0 | 0.94 | 0.006413 | 0.20 |
| C00836 | *Sphinganine* | 370.29 | 94.02 | 0.85<br>4 | 0.01<br>4 | 0.96 | 0.008237 | 0.20 |
| C00370 | *Sterol* | 250.22 | 488.8<br>8 | 0.92<br>5 | 0.00<br>003 | 0.96 | 3.09E-05 | 0.02 |
| C00219 | *Arachidonic acid* | 307.25 | 518.1<br>0 | 0.97<br>3 | 0.00<br>8 | 0.99 | 0.004028 | 0.19 |
| C00093 | *Glycerol-3-phosphate* | 256.98 | 124.3<br>4 | 1.04<br>7 | 0.01<br>2 | 1.08 | 0.005887 | 0.20 |
| C00021 | *S-Adenosyl-L-homocysteine* | 407.11 | 86.78 | 1.09<br>0 | 0.00<br>8 | 1.15 | 0.003791 | 0.19 |
|  | **Fatty Acid Metabolism** |  |  |  |  |  |  | **0.00034** |
| C00318 | *L-Carnitine* | 163.12 | 88.58 | 0.73<br>3 | 0.00<br>8 | 0.91 | 0.005364 | 0.19 |
| C00249 | *Hexadecanoic acid* | 279.23 | 28.19 | 0.73<br>9 | 0.01<br>0 | 0.92 | 0.00663 | 0.20 |
| C01595 | *Linoleate* | 281.25 | 33.78 | 0.74<br>3 | 0.00<br>7 | 0.91 | 0.004619 | 0.19 |
| C01530 | *octadecenoate* | 281.25 | 33.78 | 0.74<br>3 | 0.00<br>7 | 0.91 | 0.004619 | 0.19 |
| C02571 | *O-Acetyl-L-carnitine* | 204.12 | 91.15 | 0.75<br>2 | 0.00<br>4 | 0.91 | 0.002908 | 0.19 |
| C06424 | *Tetradecanoic acid* | 251.20 | 128.3<br>3 | 0.95<br>8 | 0.01<br>4 | 0.99 | 0.007172 | 0.20 |
| C02679 | *Dodecanoic acid* | 269.17 | 17.40 | 0.96<br>3 | 0.01<br>8 | 0.99 | 0.009251 | 0.20 |
| C05274 | *Decanoyl-CoA* | 921.24 | 150.4<br>3 | 1.02<br>9 | 0.01<br>4 | 1.05 | 0.006995 | 0.20 |
| C01607 | *Phytanic acid* | 168.15 | 48.14 | 1.04<br>9 | 0.00<br>1 | 1.08 | 0.000603 | 0.11 |
|  | **Linoleate**<br>**metabolism** |  |  |  |  |  |  | 0.00035<br>3 |
| C14765 | *11-dienoic acid* | 295.23 | 29.04 | 0.70<br>9 | 0.00<br>4 | 0.89 | 0.003002 | 0.19 |
| C06426 | *gamma-Linolenic*<br>*acid; Gamolenic*<br>*acid* | 279.23 | 28.19 | 0.73<br>9 | 0.01<br>0 | 0.92 | 0.00663 | 0.20 |
| C01595 | *Linoleate* | 281.25 | 33.78 | 0.74<br>3 | 0.00<br>7 | 0.91 | 0.004619 | 0.19 |
| C08261 | *Azelaic acid* | 189.11 | 453.7<br>6 | 0.77<br>4 | 0.00<br>3 | 0.91 | 0.002267 | 0.18 |
| C14762 | *11-dienoic acid* | 381.20 | 570.8<br>8 | 1.05<br>3 | 0.02<br>0 | 1.09 | 0.009521 | 0.20 |
| C14826 | (12*R*,13*S*)-(9*Z*)-<br>12,13-<br>*Epoxyoctadecenoic*<br>*acid* | 381.20 | 570.8<br>8 | 1.05<br>3 | 0.02<br>0 | 1.09 | 0.009521 | 0.20 |
| C14825 | (9*R*,10*S*)-(12*Z*)-<br>9,10-<br>*Epoxyoctadecenoic*<br>*acid* | 381.20 | 570.8<br>8 | 1.05<br>3 | 0.02<br>0 | 1.09 | 0.009521 | 0.20 |

**Figure 2:**
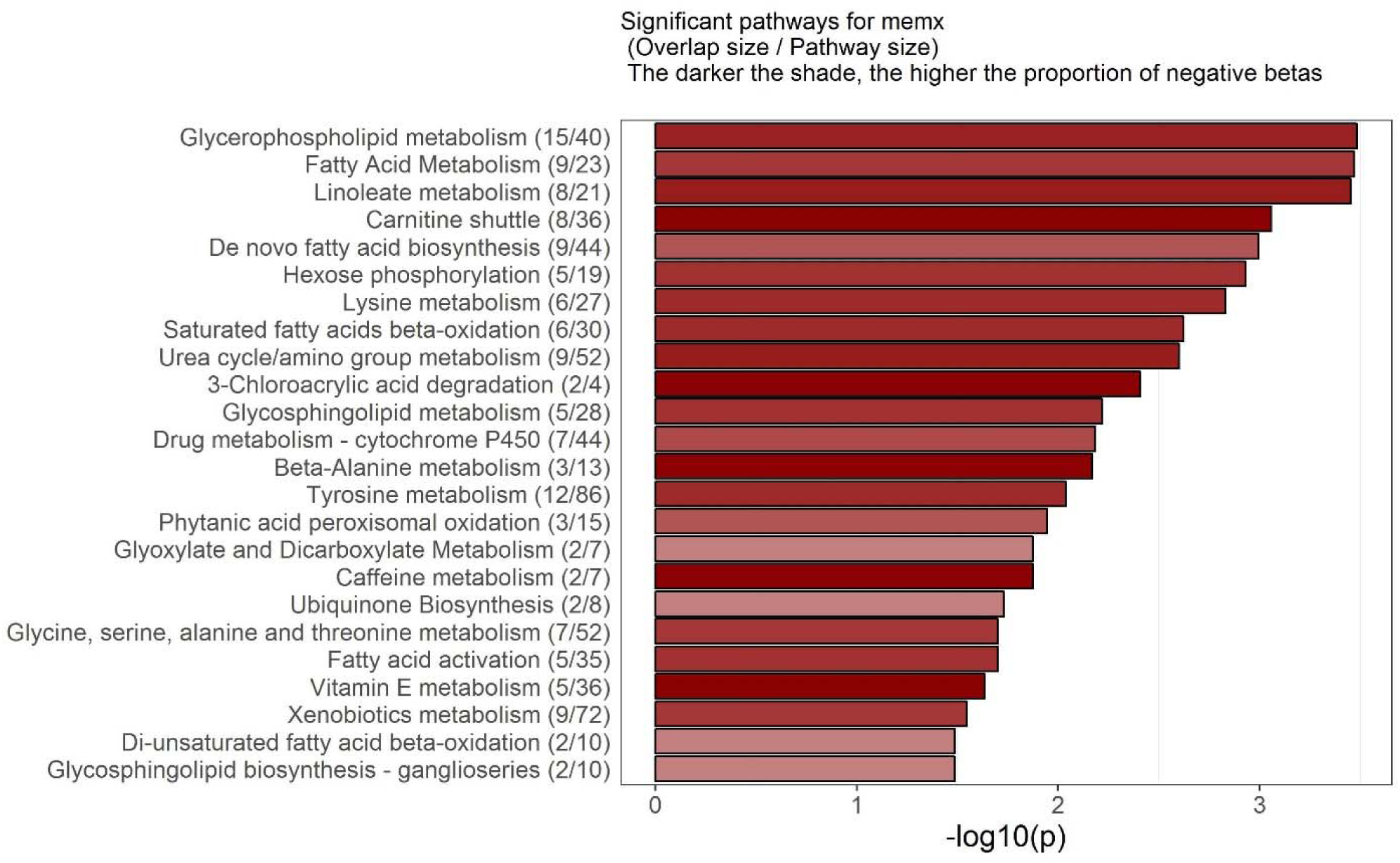
Bar-chart with p-value reflecting the pathways significantly altered in those with vs without memory decline. **Footnote:** X-axis is log p-value derived from the Mummichog package. Y axis outlines the name of the pathway, the number of significant features/total features identified in the MWAS. We consider a minimum of 3-5 significant feature within a pathway to mean a high level of certainty (significance) for its differential activity/feature abundance between the 2 groups (with and without memory changes). Numbers after the name of the pathway include (number of fetaures significant/number identified within the pathway). The darkness of the color of the bar reflects the magnitude or level of alteration in this pathway (darker= greater change in that pathway reflected by Beta from the GEE models)

Within the GCP pathway, lower levels linoleic acid, galactose, acetyl-choline, sterol, and arachidonic acid whereas higher homocysteine and glycerol3 phosphate at baseline were linked with a greater risk of having memory decline over the following 4 years. In the fatty acid pathway, lower L-carnitine, and octadeconate whereas higher phytanic acid and decanoyl coa were linked to increased risk of memory decline. In the linoleic acid metabolism, lower linoleic and azelaic acid and higher octadeconic acid were linked with greater risk of memory decline. We provide the full list of the significant pathways and the related features are provided in online supplement (**Figure S1** and **Table S2)**. Finally, in a confirmatory analysis we matched significant features identified in our analyses with prevalidated list of features and demonstrated that those in the linoleic acid pathways correctly corresponded to the validated metabolites within this pathway. In those validated, we provide the difference between those with vs without memory decline for Arachidonic and Linoleic acid in **Figure 3**.

**Figure 3:**
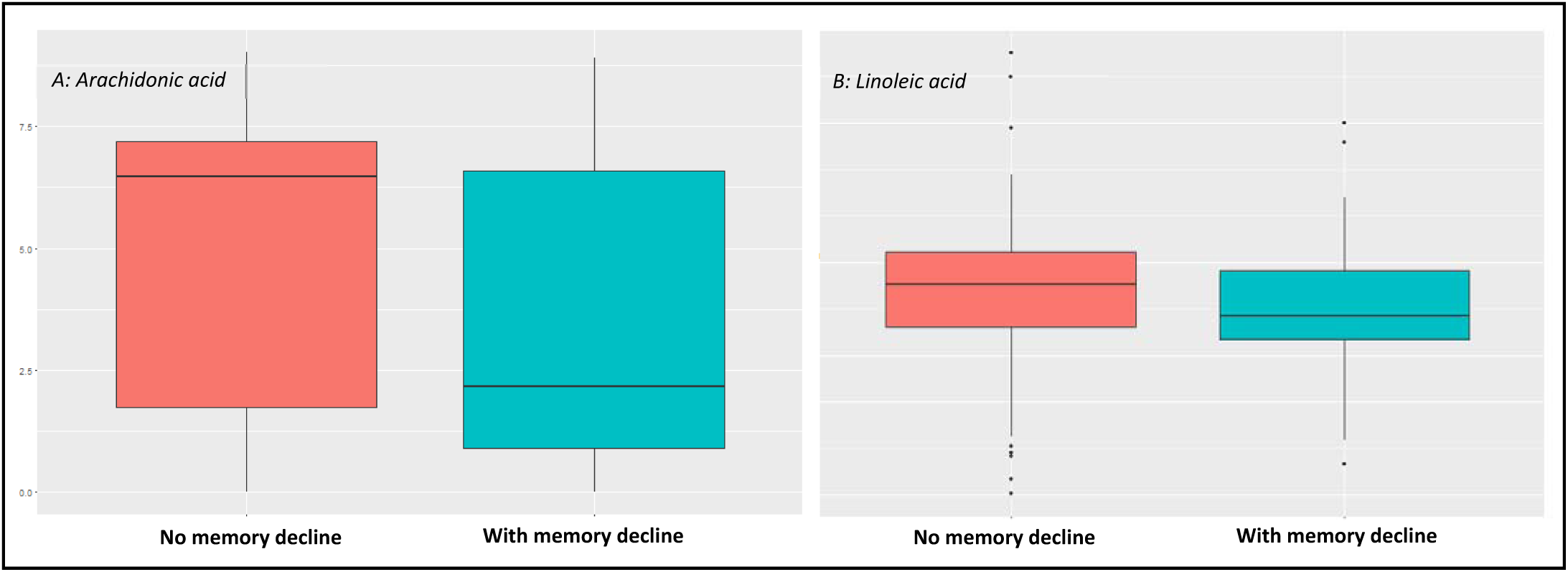
Difference between those with and without memory decline in 2 key validated metabolites Arachidonic acid (A) and Linoleic Acid (B).

## 4. Discussion

In this first Metabolome-Wide-Association-Study of early memory changes in a healthy dementia free middle aged sample, we identified multiple pathways/metabolites including phospholipids, fatty acid and linoleic metabolism that predicted memory decline. These lipid-based alterations were linked with very early memory changes and were detectable in plasma of healthy individuals.

The findings of multiple metabolites the phospholipid pathway are in agreement with prior research suggesting that glycerophospholipids alterations are highly involved in brain health and neurological disorders. ^26^ In particular, neural membrane glycerophospholipids are altered in multiple brain regions in AD patients.^27^ It was not known if these can correspond to a plsma alteration in these membrane lipids or in healthy individuals without AD. Our study suggests that these alterations are detectable in the peripheral plasma and are linked to memory changes in non-demented individuals.

Our study further describes additional key components within multiple fatty acid and lipid metabolic pathway that are linked to memory changes. The findings extends the previous reports of lipid alteration in AD to ealry memory changes. For example a higher level of multiple phospholipids (ALA^28^, Linoleic acid^29^, sterol^30^, and inositol^31^) is of particular interest since they have been previously reported to be altered in AD. More importantly, these are highly affected by dietary habits. ^32,33^ Hence, offering a new insight into the potential role of dietary interventions for memory preservations even before cognitive symptoms have appeared.

We found an association between homocysteine and memory decline even in a healthy population. These extend and support the accumulating evidence for the role of homocysteine in cognitive aging and disorders.^34-36^ Another key finding is the higher levels of acetylcholine levels in plasma are protective against future memory decline. Prior reports have suggested that acetylcholinesterase levels are altered in AD and are correlates with brain amyloid levels, and are potentially affected by many approved AD symptomatic therapies.^37^ To our knowledge this is the first report of the potential for peripheral acetylcholine levels in identifying those at risk of memory decline. Future more focused analysis may show its usefulness in identifying high risk individuals with respect to cognitive decline.

Our results in the fatty acid metabolism and beta oxidation also are in support to prior findings in memory loss. ^38^ For example, prior reports have supported that phytanic acid is neurotoxic. ^39,40^ In our MWAS we identified a negative effect of circulating levels of this molecule. Similarly carnitine has been shown to be lowered in those at risk of AD^41^ and in our study it was linked with a protective effect for memory decline.

Finally, multiple recent reports emphasizes the role of arachidonic and linoleic acid in AD. We again demonstrate that these are similarly altered in middle age prior to the evidence of memory decline. This enforces the role of neuro-inflammation in the aging brain prior to the clinical evidence neurodegeneration.^42^

There are many innovation aspects to our study including the use of high resolution Metabolomic platforms, the utilization of existing bioinformatics resources and databases to improve our detection of altered pathways, and the inclusion of healthy middle age individuals to detect early metabolic alterations that are linked to decline in memory. The limitations of our analyses is the relative small sample size with Metabolomic data conducted at one time point. The metabolome is likely to be dynamic and hence we might have missed the effect of metabolic changes on memory. Additionally, our approach relies on matching features to existing biochemical databases and not direct measurements.

## 5. Conclusion

Multiple key metabolic alterations in glycerophoshplipids, fatty acid and linoleic acid metabolism predict memory decline in middle aged healthy adults. These results support the role of these systemic fats and lipids in early changes in memory with ageing. This first MWAS of memory declines offer insight into the role of systemic metabolic changes in middle age and possibly offer targets for future research in cognitive protection against the aging process.

## Acknowledgement

Drs. Hajjar and Tianwei had full access to all the data in the study and take responsibility for the integrity of the data and the accuracy of the data analysis

