## Supplemental materials for "Untargeted Metabolomics Shows Alterations in Homocysteine, Lipids and Fatty Acids predicting Memory Decline in Healthy Middle-Aged Individuals"

**Online Figure A**: Significant pathways and fetaures mapped onto KEGG Global human metabolome (red dots are the signficnat fetaures linked to memory change in our analysis).

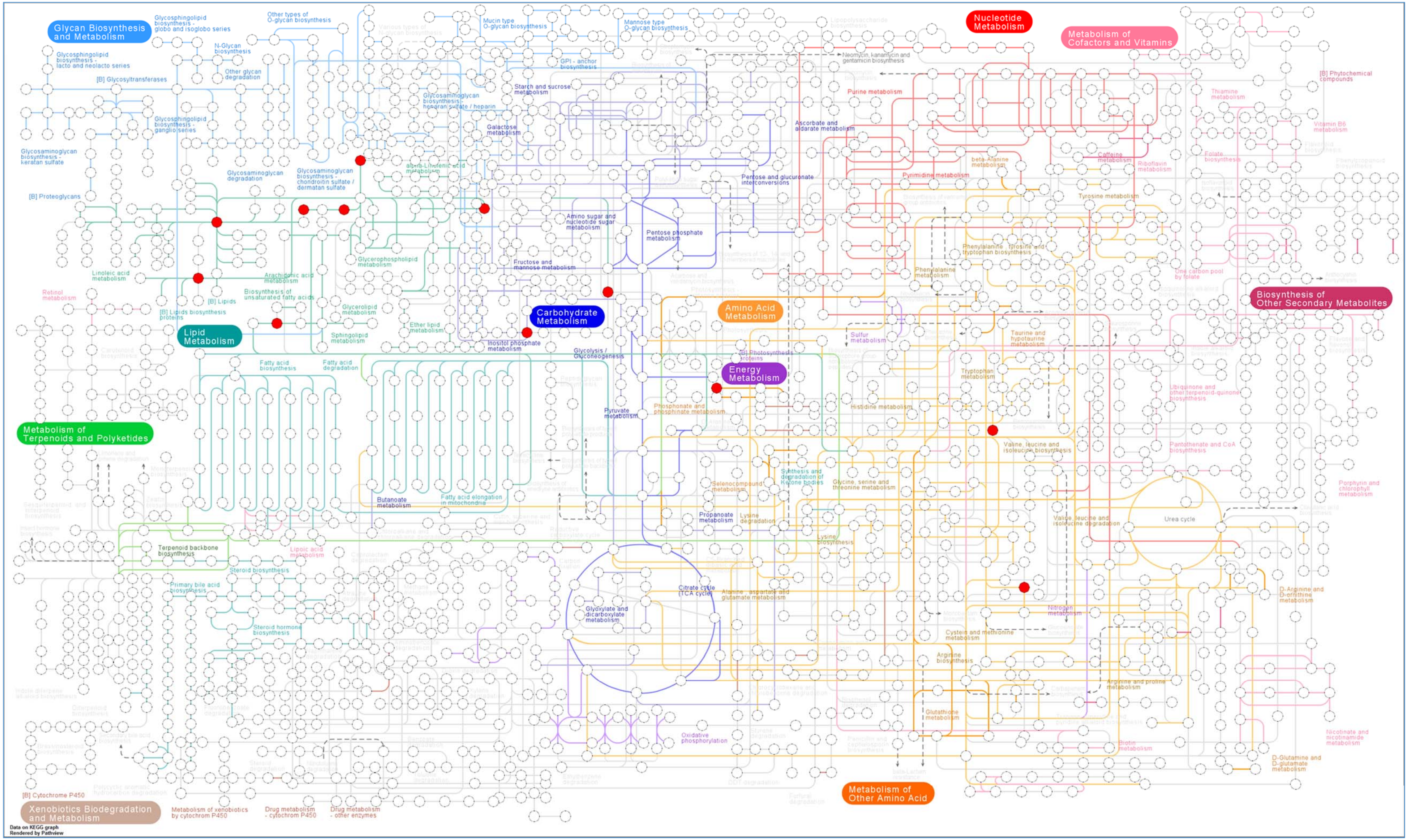

**TABLE-S1**: Mass to charge ratio, retention time and Beta with p-values and FDR for the 558 features identified as significant in the MWAS.

| M/Z values | retention time | beta | t-value | p-values | FDR |
|---|---|---|---|---|---|
| 85.04066694 | 400.5392983 | 0.180198 | 7.510854 | 0.006133 | 0.195586 |
| 86.09700659 | 93.29140287 | -0.80369 | 6.929527 | 0.008478 | 0.195586 |
| 89.07956462 | 42.08130895 | -0.9668 | 7.630159 | 0.00574 | 0.195586 |
| 90.00911912 | 202.454027 | -0.18539 | 11.5103 | 0.000692 | 0.117812 |
| 91.98031971 | 66.43651073 | -0.97457 | 6.708775 | 0.009594 | 0.196956 |
| 96.92220914 | 86.44857976 | -0.83146 | 7.023836 | 0.008043 | 0.195586 |
| 97.96881884 | 31.10987021 | 0.208896 | 7.001867 | 0.008142 | 0.195586 |
| 98.01419849 | 317.9806872 | -0.24975 | 7.160177 | 0.007454 | 0.195586 |
| 98.91922256 | 88.09806705 | -0.90657 | 7.329809 | 0.006782 | 0.195586 |
| 99.07512242 | 45.89514965 | 0.209807 | 10.54683 | 0.001164 | 0.142546 |
| 99.08098437 | 17.80027301 | -0.699 | 8.585318 | 0.003389 | 0.192905 |
| 100.0319686 | 463.6719735 | -0.10002 | 7.248022 | 0.007098 | 0.195586 |
| 100.1124794 | 34.73072382 | -0.72529 | 6.812107 | 0.009054 | 0.195998 |
| 100.1338192 | 33.94156819 | -1.15404 | 7.160261 | 0.007454 | 0.195586 |
| 100.139347 | 35.63147177 | -0.84707 | 7.133981 | 0.007564 | 0.195586 |
| 100.1392495 | 549.202534 | -0.1246 | 9.358697 | 0.002219 | 0.17585 |
| 100.9171256 | 77.49271043 | -1.13622 | 7.519999 | 0.006102 | 0.195586 |
| 101.1158321 | 37.46772179 | -0.84478 | 6.982159 | 0.008233 | 0.195586 |
| 103.0546666 | 114.0019938 | -0.91763 | 7.979919 | 0.00473 | 0.192905 |
| 104.079076 | 68.26414941 | -0.71587 | 11.18192 | 0.000826 | 0.125687 |
| 104.1074439 | 78.74522179 | -0.72076 | 7.430462 | 0.006413 | 0.195586 |
| 104.1359015 | 68.29826612 | -0.78067 | 11.77746 | 0.0006 | 0.108186 |
| 105.1106762 | 69.1993613 | -0.89355 | 8.449155 | 0.003652 | 0.192905 |
| 106.112492 | 68.78086745 | -0.77012 | 8.884887 | 0.002875 | 0.192156 |
| 106.9920833 | 128.7750758 | 0.140413 | 18.56813 | 1.64E-05 | 0.018294 |
| 107.0428852 | 84.85174501 | -1.04367 | 7.262111 | 0.007042 | 0.195586 |
| 109.0147508 | 30.22291024 | 0.153293 | 7.649916 | 0.005677 | 0.195586 |
| 110.0447117 | 161.3732951 | -0.20841 | 6.515675 | 0.010693 | 0.199227 |
| 112.0393418 | 376.4722357 | -0.09378 | 7.913579 | 0.004907 | 0.192905 |
| 112.8955796 | 85.73330894 | -0.79966 | 7.179502 | 0.007374 | 0.195586 |
| 112.9784458 | 126.9723738 | 0.528178 | 14.14747 | 0.000169 | 0.060625 |
| 114.066525 | 80.86940499 | -0.80582 | 7.775731 | 0.005295 | 0.194197 |
| 114.8928933 | 85.63922517 | -0.85302 | 7.159226 | 0.007458 | 0.195586 |
| 115.0697839 | 75.50640907 | -0.86819 | 6.636315 | 0.009992 | 0.199227 |
| 116.8910676 | 88.16746559 | -0.96676 | 7.308969 | 0.006861 | 0.195586 |
| 118.1228482 | 33.56656789 | -0.68185 | 6.449196 | 0.0111 | 0.199785 |
| 118.2260412 | 28.04716206 | -0.16417 | 7.179582 | 0.007374 | 0.195586 |
| 119.0493829 | 113.5225346 | -0.42041 | 9.975531 | 0.001586 | 0.156484 |

| | | | | | |
|---|---|---|---|---|---|
| 120.0444826 | 441.4022629 | -0.19221 | 9.974096 | 0.001588 | 0.156484 |
| 120.0809807 | 119.2121082 | -0.76676 | 7.456895 | 0.006319 | 0.195586 |
| 120.0914521 | 120.2025907 | -0.83129 | 6.953729 | 0.008364 | 0.195586 |
| 120.1041234 | 68.84299712 | 0.102463 | 9.437092 | 0.002126 | 0.172222 |
| 120.1230976 | 19.42142704 | -0.76269 | 6.511634 | 0.010717 | 0.199227 |
| 121.084333 | 112.8476705 | -0.8615 | 7.451644 | 0.006338 | 0.195586 |
| 121.1021027 | 549.5061942 | -1.08013 | 6.519679 | 0.010669 | 0.199227 |
| 121.9739308 | 83.50881687 | -0.87528 | 7.172002 | 0.007405 | 0.195586 |
| 122.9778195 | 73.80475157 | -1.18712 | 6.863047 | 0.0088 | 0.195586 |
| 123.9712282 | 81.32718004 | -0.92985 | 6.97151 | 0.008282 | 0.195586 |
| 129.0530982 | 331.0376475 | -0.9492 | 8.121853 | 0.004373 | 0.192905 |
| 130.0598344 | 19.98804994 | -0.22935 | 7.20311 | 0.007278 | 0.195586 |
| 130.159113 | 64.39003577 | -0.98033 | 8.118879 | 0.004381 | 0.192905 |
| 132.0352746 | 219.0164097 | -0.25725 | 10.25935 | 0.00136 | 0.156484 |
| 132.1019745 | 115.7493493 | -0.70868 | 7.176316 | 0.007387 | 0.195586 |
| 132.5313192 | 213.5426584 | -0.14298 | 9.825647 | 0.001721 | 0.156484 |
| 133.0066441 | 64.77408193 | -0.97685 | 7.537137 | 0.006044 | 0.195586 |
| 133.0329398 | 481.7674867 | -0.27851 | 8.992354 | 0.002711 | 0.192156 |
| 133.105292 | 110.0345562 | -0.77177 | 6.892528 | 0.008656 | 0.195586 |
| 136.0701418 | 559.167163 | -0.38927 | 7.080744 | 0.007792 | 0.195586 |
| 137.9482679 | 79.07664883 | -0.89526 | 6.886394 | 0.008685 | 0.195586 |
| 137.9870283 | 323.8822378 | -0.28672 | 9.896539 | 0.001656 | 0.156484 |
| 137.9986259 | 295.1410101 | 0.279566 | 7.3398 | 0.006744 | 0.195586 |
| 138.0914433 | 32.60634949 | -0.57104 | 8.151229 | 0.004303 | 0.192905 |
| 138.9065561 | 81.30094793 | -1.14263 | 8.167064 | 0.004266 | 0.192905 |
| 139.945193 | 76.77779463 | -1.03965 | 7.579157 | 0.005905 | 0.195586 |
| 139.9822489 | 305.0112022 | -0.81701 | 7.504731 | 0.006154 | 0.195586 |
| 140.0106896 | 23.6056295 | -1.03064 | 7.990268 | 0.004703 | 0.192905 |
| 140.0688697 | 119.1617142 | -0.64269 | 6.937238 | 0.008442 | 0.195586 |
| 140.0934863 | 424.1545567 | 0.219177 | 9.851357 | 0.001697 | 0.156484 |
| 140.9032516 | 77.19547728 | -1.26337 | 8.124627 | 0.004367 | 0.192905 |
| 140.982056 | 310.601687 | -0.91491 | 7.524232 | 0.006087 | 0.195586 |
| 140.9959208 | 31.66153769 | 0.284628 | 9.850907 | 0.001697 | 0.156484 |
| 141.0141121 | 546.4195964 | -1.19083 | 6.69938 | 0.009645 | 0.196956 |
| 141.0912382 | 18.05920667 | -0.9911 | 7.487818 | 0.006212 | 0.195586 |
| 141.9791038 | 314.8857112 | -0.98232 | 8.510187 | 0.003532 | 0.192905 |
| 143.1266657 | 425.5861123 | -0.68443 | 10.65578 | 0.001097 | 0.141281 |
| 143.9809891 | 348.6192831 | -1.04476 | 13.67793 | 0.000217 | 0.068101 |
| 146.1175766 | 38.69227607 | -0.83355 | 6.502598 | 0.010772 | 0.199227 |
| 147.0437218 | 118.4691122 | -0.73348 | 6.677148 | 0.009766 | 0.197337 |
| 148.934188 | 68.55591265 | -0.96909 | 6.700856 | 0.009637 | 0.196956 |

| | | | | | |
|---|---|---|---|---|---|
| 149.0598274 | 587.0655521 | -1.03388 | 7.165351 | 0.007433 | 0.195586 |
| 149.1328327 | 553.0016241 | -0.58399 | 17.64634 | 2.66E-05 | 0.020552 |
| 150.9313122 | 68.41116854 | -0.92193 | 6.723606 | 0.009515 | 0.196956 |
| 152.0215565 | 72.4853062 | -0.90084 | 9.035838 | 0.002647 | 0.191572 |
| 154.0202236 | 71.02044566 | -1.02806 | 8.358973 | 0.003838 | 0.192905 |
| 154.8797761 | 79.77303363 | -0.89702 | 7.184808 | 0.007352 | 0.195586 |
| 155.5048905 | 127.4049566 | 0.098594 | 6.57216 | 0.010359 | 0.199227 |
| 156.0417746 | 114.6904196 | -0.64401 | 6.708631 | 0.009595 | 0.196956 |
| 156.1384516 | 572.8703509 | -0.6667 | 6.617088 | 0.0101 | 0.199227 |
| 156.8773318 | 79.87156583 | -1.07734 | 7.891905 | 0.004966 | 0.192905 |
| 157.013341 | 137.2151832 | 0.337457 | 7.774719 | 0.005298 | 0.194197 |
| 157.0132073 | 568.6917011 | 0.300815 | 6.549456 | 0.010492 | 0.199227 |
| 157.9940457 | 480.1863886 | -0.37848 | 7.342602 | 0.006734 | 0.195586 |
| 161.0185024 | 80.24859603 | -1.03981 | 8.544306 | 0.003466 | 0.192905 |
| 162.1125459 | 93.30364711 | -0.91151 | 8.130122 | 0.004354 | 0.192905 |
| 163.0868185 | 416.9397585 | 0.152876 | 21.99998 | 2.73E-06 | 0.006846 |
| 163.1158014 | 88.57765923 | -1.03676 | 7.752535 | 0.005364 | 0.194469 |
| 164.9085174 | 80.60347979 | -0.86024 | 6.769526 | 0.009273 | 0.196469 |
| 165.0546954 | 116.3672439 | -0.80434 | 6.99009 | 0.008196 | 0.195586 |
| 166.0862292 | 119.8723206 | -0.75409 | 8.024548 | 0.004615 | 0.192905 |
| 166.0982938 | 49.14901517 | 0.166584 | 15.22197 | 9.56E-05 | 0.045476 |
| 166.1340133 | 47.32670634 | 0.190335 | 18.21952 | 1.97E-05 | 0.01977 |
| 167.0126521 | 9.898183893 | -0.16083 | 6.812304 | 0.009053 | 0.195998 |
| 167.0926732 | 49.69771729 | 0.269294 | 9.511902 | 0.002041 | 0.170851 |
| 167.514567 | 127.3866504 | 0.119466 | 8.868568 | 0.002901 | 0.192156 |
| 168.1494235 | 48.14287546 | 0.159669 | 11.76593 | 0.000603 | 0.108186 |
| 169.0860056 | 464.2730752 | -1.16214 | 9.55084 | 0.001999 | 0.16867 |
| 169.133534 | 567.0159721 | -0.33671 | 6.966587 | 0.008305 | 0.195586 |
| 170.0964043 | 44.24422796 | -0.83613 | 6.479948 | 0.01091 | 0.199576 |
| 170.8540386 | 88.51415116 | -0.99348 | 7.87852 | 0.005003 | 0.192905 |
| 171.0138459 | 55.23433448 | -0.31953 | 9.610953 | 0.001934 | 0.166027 |
| 172.0401051 | 89.69090601 | -0.87407 | 7.006651 | 0.008121 | 0.195586 |
| 172.8512748 | 88.63879396 | -1.0192 | 8.071593 | 0.004496 | 0.192905 |
| 172.9771695 | 26.41699942 | -0.15182 | 6.551421 | 0.01048 | 0.199227 |
| 174.1502071 | 377.2224822 | -0.09138 | 7.044006 | 0.007953 | 0.195586 |
| 174.5133412 | 148.6765235 | 0.122208 | 8.370239 | 0.003814 | 0.192905 |
| 174.9572157 | 349.0349592 | -0.53186 | 18.61957 | 1.60E-05 | 0.018294 |
| 176.9718748 | 328.3602593 | -0.925 | 9.070948 | 0.002597 | 0.191572 |
| 177.0744798 | 74.16461778 | -0.8988 | 6.736986 | 0.009443 | 0.196956 |
| 178.0587525 | 128.8388946 | -0.97551 | 10.60285 | 0.001129 | 0.141605 |
| 179.1430622 | 550.0073101 | -1.02063 | 6.722918 | 0.009518 | 0.196956 |

| | | | | | |
|---|---|---|---|---|---|
| 179.5492302 | 140.9644804 | 0.097383 | 7.887249 | 0.004978 | 0.192905 |
| 179.9322563 | 74.02153557 | -1.09318 | 7.023456 | 0.008045 | 0.195586 |
| 180.8827357 | 86.18165458 | -0.89086 | 6.629992 | 0.010028 | 0.199227 |
| 181.9293801 | 73.1106726 | -1.19704 | 7.650115 | 0.005677 | 0.195586 |
| 182.0811497 | 115.684673 | -0.72354 | 6.646278 | 0.009936 | 0.199069 |
| 182.1284819 | 44.25241981 | 0.19897 | 9.288016 | 0.002307 | 0.179573 |
| 182.511771 | 127.7259209 | 0.12502 | 6.563173 | 0.010411 | 0.199227 |
| 183.0350225 | 127.88159 | 0.257393 | 12.89686 | 0.000329 | 0.0895 |
| 185.114641 | 410.2522135 | -1.07723 | 8.270266 | 0.00403 | 0.192905 |
| 185.1671677 | 572.1771111 | 0.121371 | 8.645436 | 0.003279 | 0.192905 |
| 185.5405164 | 137.0855367 | 0.107137 | 10.05925 | 0.001516 | 0.156484 |
| 186.2216091 | 473.2974324 | -0.88663 | 7.393587 | 0.006546 | 0.195586 |
| 186.8283258 | 95.91070818 | -0.95586 | 7.630506 | 0.005739 | 0.195586 |
| 187.5401596 | 128.5412001 | 0.135753 | 18.94215 | 1.35E-05 | 0.018294 |
| 189.0733952 | 109.2357067 | -0.76806 | 6.849366 | 0.008867 | 0.195586 |
| 189.1123907 | 40.75737488 | -0.17457 | 7.622679 | 0.005764 | 0.195586 |
| 189.1120392 | 453.7583875 | -0.85308 | 9.319517 | 0.002267 | 0.177891 |
| 189.9617957 | 68.51083823 | -1.279 | 7.025452 | 0.008036 | 0.195586 |
| 191.0403807 | 116.1727381 | -0.93408 | 8.654173 | 0.003263 | 0.192905 |
| 191.5169107 | 126.4251045 | 0.110107 | 6.795056 | 0.009141 | 0.196469 |
| 191.8866077 | 139.548091 | -0.12235 | 6.773204 | 0.009254 | 0.196469 |
| 192.0415978 | 104.3776768 | -0.90817 | 6.966728 | 0.008304 | 0.195586 |
| 193.1588404 | 551.1012196 | -0.96997 | 6.482244 | 0.010896 | 0.199576 |
| 195.1862619 | 61.82140408 | 0.140448 | 6.863327 | 0.008798 | 0.195586 |
| 197.5220036 | 169.3038342 | 0.166351 | 9.959938 | 0.0016 | 0.156484 |
| 198.1851546 | 30.3141648 | -1.29068 | 11.0768 | 0.000874 | 0.131031 |
| 200.1837864 | 30.16447567 | -0.91019 | 7.716127 | 0.005473 | 0.195586 |
| 200.1838229 | 499.6552468 | -0.95286 | 8.076729 | 0.004484 | 0.192905 |
| 200.2009189 | 40.89246122 | -0.96378 | 8.033406 | 0.004592 | 0.192905 |
| 200.2010821 | 586.3039495 | -1.03921 | 7.487323 | 0.006213 | 0.195586 |
| 203.0537326 | 74.29784637 | -1.15318 | 8.868542 | 0.002901 | 0.192156 |
| 203.1502688 | 122.6859895 | -1.07162 | 6.544399 | 0.010521 | 0.199227 |
| 204.1222975 | 91.15248909 | -0.952 | 8.864144 | 0.002908 | 0.192156 |
| 204.1386058 | 22.79133422 | -0.86315 | 6.827383 | 0.008977 | 0.195586 |
| 207.1742989 | 27.06228352 | -0.85353 | 6.833685 | 0.008945 | 0.195586 |
| 208.889623 | 75.27451454 | -1.21463 | 9.821017 | 0.001725 | 0.156484 |
| 210.8866703 | 70.9240815 | -0.64684 | 8.427046 | 0.003697 | 0.192905 |
| 212.8375748 | 76.24602756 | -1.1632 | 8.436577 | 0.003677 | 0.192905 |
| 213.1461093 | 493.9448294 | -0.75566 | 6.989782 | 0.008198 | 0.195586 |
| 213.1596165 | 23.94984948 | -0.98879 | 7.117687 | 0.007633 | 0.195586 |
| 213.4984344 | 137.4531149 | 0.566164 | 6.704176 | 0.009619 | 0.196956 |

| | | | | | |
|---|---|---|---|---|---|
| 214.8349824 | 75.99430291 | -1.16908 | 8.506831 | 0.003538 | 0.192905 |
| 215.1254266 | 42.99477684 | -0.52579 | 7.272284 | 0.007003 | 0.195586 |
| 215.9799595 | 454.2295993 | -0.23982 | 8.098497 | 0.00443 | 0.192905 |
| 216.0607437 | 82.9319316 | -0.87849 | 6.617078 | 0.010101 | 0.199227 |
| 217.0640556 | 92.08318418 | -0.28216 | 9.44022 | 0.002123 | 0.172222 |
| 217.1045601 | 70.15922513 | -0.6769 | 6.662822 | 0.009844 | 0.198133 |
| 217.5367352 | 129.3977633 | 0.111416 | 8.485016 | 0.003581 | 0.192905 |
| 217.9542492 | 195.7119292 | -0.11955 | 12.0062 | 0.00053 | 0.105681 |
| 219.1746039 | 545.3051205 | -0.68273 | 7.59409 | 0.005856 | 0.195586 |
| 221.1895885 | 465.2725212 | -0.89471 | 6.874523 | 0.008743 | 0.195586 |
| 222.8659329 | 73.27967518 | -0.99706 | 7.425342 | 0.006431 | 0.195586 |
| 223.0635815 | 21.74096859 | -1.20301 | 9.045331 | 0.002634 | 0.191572 |
| 223.0634033 | 530.2806734 | -0.93151 | 7.30153 | 0.00689 | 0.195586 |
| 223.1328981 | 522.1245205 | -0.9665 | 7.273007 | 0.007 | 0.195586 |
| 223.9627249 | 139.6202806 | 0.080572 | 6.56585 | 0.010395 | 0.199227 |
| 224.8649905 | 70.44137224 | -1.00791 | 7.18757 | 0.007341 | 0.195586 |
| 225.0346469 | 75.44071254 | -0.99793 | 8.764061 | 0.003072 | 0.192905 |
| 226.5260258 | 136.8016349 | 0.107657 | 8.670897 | 0.003233 | 0.192905 |
| 228.2321176 | 47.47595626 | -1.00778 | 8.905204 | 0.002844 | 0.192156 |
| 228.8117158 | 88.17879217 | -1.08382 | 7.922768 | 0.004882 | 0.192905 |
| 229.1050885 | 410.410172 | -0.18443 | 7.276924 | 0.006985 | 0.195586 |
| 229.23443 | 44.89114305 | -1.13301 | 9.098513 | 0.002558 | 0.191572 |
| 230.8088187 | 88.49521587 | -1.10462 | 8.324918 | 0.00391 | 0.192905 |
| 230.9151689 | 143.9478047 | -0.8933 | 7.264252 | 0.007034 | 0.195586 |
| 232.190568 | 424.5930955 | 0.179843 | 15.09844 | 0.000102 | 0.045476 |
| 233.1897161 | 588.5725376 | -0.17058 | 8.095495 | 0.004438 | 0.192905 |
| 234.8938081 | 70.7222922 | -0.86405 | 6.449562 | 0.011098 | 0.199785 |
| 235.180427 | 572.5929462 | -0.92996 | 6.909396 | 0.008574 | 0.195586 |
| 235.9580522 | 58.41778185 | 0.097389 | 7.668222 | 0.00562 | 0.195586 |
| 236.0951595 | 145.7497085 | 0.132088 | 10.13702 | 0.001453 | 0.156484 |
| 238.8399939 | 85.73449118 | -0.93079 | 6.832829 | 0.00895 | 0.195586 |
| 239.2129459 | 18.977446 | -0.15955 | 7.249966 | 0.00709 | 0.195586 |
| 240.8389607 | 85.33394934 | -0.9615 | 6.960912 | 0.008331 | 0.195586 |
| 241.1786909 | 45.82580629 | -0.60705 | 10.79532 | 0.001018 | 0.139993 |
| 243.1356523 | 54.13640305 | 0.093043 | 6.579041 | 0.010319 | 0.199227 |
| 244.7865997 | 94.09665121 | -0.98403 | 7.282823 | 0.006962 | 0.195586 |
| 246.5390144 | 137.7795957 | 0.114107 | 6.731639 | 0.009472 | 0.196956 |
| 246.7837341 | 94.72750631 | -1.0707 | 8.042431 | 0.004569 | 0.192905 |
| 246.9543522 | 151.6833079 | 0.09466 | 7.988027 | 0.004709 | 0.192905 |
| 249.0606721 | 99.35982412 | -0.8845 | 8.017933 | 0.004632 | 0.192905 |
| 249.2212521 | 17.38127164 | -1.0283 | 6.684251 | 0.009727 | 0.197313 |

| | | | | | |
|---|---|---|---|---|---|
| 249.8454157 | 135.929084 | -0.13763 | 7.57322 | 0.005924 | 0.195586 |
| 250.1192798 | 35.28905265 | -0.80024 | 6.498531 | 0.010796 | 0.199316 |
| 250.2242621 | 488.87855 | -0.25863 | 17.35942 | 3.09E-05 | 0.022192 |
| 251.0001041 | 549.9986375 | -1.04158 | 7.183496 | 0.007358 | 0.195586 |
| 251.1971601 | 128.3250616 | -0.14377 | 7.229503 | 0.007172 | 0.195586 |
| 252.9972693 | 17.65429948 | -1.00134 | 7.225165 | 0.007189 | 0.195586 |
| 252.9970382 | 550.2136249 | -1.09833 | 7.625345 | 0.005755 | 0.195586 |
| 254.81423 | 91.80132167 | -0.9099 | 7.120588 | 0.00762 | 0.195586 |
| 254.9867635 | 134.0977431 | 0.199145 | 6.976773 | 0.008257 | 0.195586 |
| 256.2633285 | 77.00368731 | -0.99125 | 8.798384 | 0.003015 | 0.192905 |
| 256.8116184 | 92.44289499 | -0.93202 | 7.155309 | 0.007474 | 0.195586 |
| 256.98379 | 124.3396881 | 0.153295 | 7.584476 | 0.005887 | 0.195586 |
| 257.1467964 | 121.2918009 | -0.55799 | 8.274324 | 0.004021 | 0.192905 |
| 257.2667451 | 568.3585272 | -0.56659 | 8.028026 | 0.004606 | 0.192905 |
| 258.1508872 | 98.6969406 | -0.16757 | 13.26592 | 0.00027 | 0.079834 |
| 258.8863086 | 335.5018179 | -0.12426 | 7.107588 | 0.007676 | 0.195586 |
| 260.9222161 | 83.27587592 | -0.21566 | 7.853377 | 0.005073 | 0.193703 |
| 263.2367507 | 32.31618916 | -0.8996 | 7.020007 | 0.00806 | 0.195586 |
| 263.8215642 | 138.2901893 | -0.14465 | 7.101153 | 0.007703 | 0.195586 |
| 264.8432095 | 91.28267129 | -0.78181 | 6.592334 | 0.010242 | 0.199227 |
| 264.9725535 | 114.6804577 | -1.0135 | 7.763939 | 0.00533 | 0.194197 |
| 266.6081517 | 576.0526799 | -0.88903 | 6.613482 | 0.010121 | 0.199227 |
| 269.1745588 | 17.39933981 | -0.12732 | 6.773654 | 0.009251 | 0.196469 |
| 270.7954461 | 80.22669827 | -1.24225 | 7.704784 | 0.005507 | 0.195586 |
| 271.0383688 | 88.4284262 | -1.20278 | 11.70332 | 0.000624 | 0.109924 |
| 272.7924776 | 80.72634346 | -1.22721 | 8.170015 | 0.004259 | 0.192905 |
| 277.0606625 | 76.67790354 | -0.79939 | 6.699699 | 0.009643 | 0.196956 |
| 277.2157743 | 49.81840758 | -0.12494 | 6.566941 | 0.010389 | 0.199227 |
| 278.9099165 | 127.8812289 | 0.154656 | 8.714738 | 0.003156 | 0.192905 |
| 279.0935836 | 23.32580687 | -1.00317 | 7.342754 | 0.006733 | 0.195586 |
| 279.0932266 | 508.3480615 | -1.03804 | 6.88203 | 0.008707 | 0.195586 |
| 279.2319117 | 28.18917246 | -1.00717 | 7.370558 | 0.00663 | 0.195586 |
| 280.2343463 | 545.5037103 | -0.42171 | 15.84131 | 6.89E-05 | 0.038432 |
| 280.8240286 | 78.64605407 | -1.13961 | 7.522073 | 0.006095 | 0.195586 |
| 281.2475225 | 33.7840677 | -0.99152 | 8.022817 | 0.004619 | 0.192905 |
| 281.2475416 | 556.6516687 | -0.85894 | 6.538965 | 0.010554 | 0.199227 |
| 281.9132977 | 127.4812729 | 0.815731 | 33.6451 | 6.61E-09 | 6.64E-05 |
| 282.8210672 | 75.9433787 | -1.15426 | 7.642637 | 0.0057 | 0.195586 |
| 284.8189757 | 73.0189317 | -1.16488 | 6.509294 | 0.010731 | 0.199227 |
| 285.2968964 | 40.50116357 | -0.81175 | 7.802401 | 0.005218 | 0.194197 |
| 285.2976587 | 586.9952701 | -1.27911 | 10.58175 | 0.001142 | 0.141605 |

| | | | | | |
|---|---|---|---|---|---|
| 286.7698358 | 84.23193573 | -1.18264 | 7.781226 | 0.005279 | 0.194197 |
| 288.2884755 | 91.34264353 | -0.42192 | 8.09147 | 0.004447 | 0.192905 |
| 288.7667655 | 86.11074202 | -1.12621 | 7.787056 | 0.005262 | 0.194197 |
| 289.0419796 | 66.09733118 | -0.11032 | 6.770188 | 0.009269 | 0.196469 |
| 289.8328777 | 136.9004242 | -0.1764 | 15.61113 | 7.78E-05 | 0.041121 |
| 290.8533004 | 73.8355392 | -0.94153 | 6.591982 | 0.010244 | 0.199227 |
| 292.6940947 | 148.4209565 | 0.146904 | 8.314766 | 0.003932 | 0.192905 |
| 292.8504367 | 74.05591347 | -0.96276 | 6.658872 | 0.009866 | 0.198175 |
| 292.885056 | 131.7146653 | 0.372566 | 10.91675 | 0.000953 | 0.134801 |
| 293.8274583 | 137.6773864 | -0.27734 | 12.69301 | 0.000367 | 0.0895 |
| 293.9667764 | 141.6953739 | 0.133293 | 14.01327 | 0.000182 | 0.060768 |
| 295.1508532 | 25.57254938 | -0.09768 | 9.948917 | 0.001609 | 0.156484 |
| 295.2264063 | 29.03783535 | -1.14831 | 8.806241 | 0.003002 | 0.192905 |
| 296.7983132 | 84.98735821 | -1.06302 | 7.061531 | 0.007876 | 0.195586 |
| 297.2422652 | 35.20754479 | -0.98818 | 6.643739 | 0.00995 | 0.199069 |
| 298.5547499 | 134.7870781 | 0.142771 | 10.00567 | 0.001561 | 0.156484 |
| 298.7953466 | 84.91408591 | -1.0413 | 6.829586 | 0.008966 | 0.195586 |
| 299.1710771 | 41.28086902 | 0.126038 | 7.040478 | 0.007969 | 0.195586 |
| 299.2580106 | 31.395483 | -1.02102 | 7.863894 | 0.005043 | 0.193315 |
| 299.9116788 | 22.84229992 | 0.126176 | 8.986266 | 0.00272 | 0.192156 |
| 300.1395854 | 120.7901237 | -0.11174 | 8.246263 | 0.004084 | 0.192905 |
| 300.2605836 | 557.5198895 | -0.85693 | 6.462636 | 0.011017 | 0.199785 |
| 303.9598498 | 125.8794446 | 0.135331 | 8.138398 | 0.004334 | 0.192905 |
| 306.2426492 | 559.1041495 | -0.9511 | 8.489473 | 0.003572 | 0.192905 |
| 307.2465652 | 518.1047968 | -0.09288 | 8.271081 | 0.004028 | 0.192905 |
| 307.8038964 | 136.5156428 | -0.16218 | 12.12026 | 0.000499 | 0.103905 |
| 308.122476 | 71.70888497 | -0.13578 | 9.052604 | 0.002623 | 0.191572 |
| 308.82399 | 82.80307528 | -0.99885 | 7.262949 | 0.007039 | 0.195586 |
| 309.1306991 | 511.1722873 | -0.89543 | 7.450382 | 0.006342 | 0.195586 |
| 309.226965 | 20.8249898 | -0.42514 | 8.880431 | 0.002882 | 0.192156 |
| 309.6572735 | 559.312205 | -0.18752 | 8.525143 | 0.003503 | 0.192905 |
| 311.0076804 | 81.77328007 | -0.93275 | 6.563273 | 0.01041 | 0.199227 |
| 312.3253075 | 582.9536501 | -1.10983 | 7.335676 | 0.00676 | 0.195586 |
| 316.2470692 | 92.08272271 | -0.87232 | 8.478161 | 0.003594 | 0.192905 |
| 317.2684696 | 30.77335741 | -1.13115 | 7.81945 | 0.005169 | 0.194197 |
| 317.9378193 | 116.0933797 | 0.170319 | 8.242213 | 0.004093 | 0.192905 |
| 323.0561587 | 133.541323 | 0.094795 | 7.886246 | 0.004981 | 0.192905 |
| 323.0820706 | 119.5757366 | 0.299973 | 8.513378 | 0.003525 | 0.192905 |
| 326.2689548 | 22.82420012 | -0.1135 | 6.94707 | 0.008396 | 0.195586 |
| 327.0078309 | 550.4371931 | -0.92332 | 7.40968 | 0.006487 | 0.195586 |
| 327.9685737 | 127.4004058 | 0.101718 | 7.601357 | 0.005832 | 0.195586 |

| | | | | | |
|---|---|---|---|---|---|
| 328.2404807 | 539.1097652 | 0.106405 | 8.767061 | 0.003067 | 0.192905 |
| 330.9332233 | 69.24516101 | 0.116291 | 11.20454 | 0.000816 | 0.125687 |
| 331.9159955 | 300.1901002 | -0.12761 | 12.41438 | 0.000426 | 0.093016 |
| 332.007824 | 18.42721041 | -1.06538 | 7.05838 | 0.00789 | 0.195586 |
| 332.9987134 | 549.4728398 | -1.01052 | 6.839162 | 0.008918 | 0.195586 |
| 336.272348 | 42.07895789 | -0.56766 | 17.69634 | 2.59E-05 | 0.020552 |
| 338.0377462 | 171.9015874 | -0.32757 | 14.25233 | 0.00016 | 0.059463 |
| 338.7823321 | 74.00764197 | -1.22588 | 7.346939 | 0.006718 | 0.195586 |
| 338.8313214 | 67.79221065 | -0.13253 | 7.558791 | 0.005972 | 0.195586 |
| 339.2511683 | 554.4216748 | -1.00252 | 7.963248 | 0.004774 | 0.192905 |
| 340.7792039 | 74.41124499 | -1.25345 | 7.761593 | 0.005337 | 0.194197 |
| 340.9022871 | 116.0098258 | -0.30289 | 9.033031 | 0.002651 | 0.191572 |
| 340.9640574 | 80.41465297 | -0.2793 | 9.837413 | 0.00171 | 0.156484 |
| 341.2675421 | 84.8252707 | -0.13614 | 13.78754 | 0.000205 | 0.066313 |
| 342.5982027 | 127.2887164 | 0.322694 | 6.833917 | 0.008944 | 0.195586 |
| 343.9408736 | 127.7312651 | 0.159386 | 7.08922 | 0.007755 | 0.195586 |
| 346.7252165 | 83.55890611 | -1.27157 | 8.029562 | 0.004602 | 0.192905 |
| 348.8111652 | 71.23696174 | -1.07775 | 7.371501 | 0.006627 | 0.195586 |
| 348.9898827 | 315.8374043 | -1.04849 | 7.882423 | 0.004992 | 0.192905 |
| 350.8087599 | 71.52280584 | -1.10617 | 7.979508 | 0.004731 | 0.192905 |
| 351.6030104 | 125.7334073 | 0.323543 | 10.06209 | 0.001514 | 0.156484 |
| 352.8056983 | 71.59684835 | -1.16317 | 7.67938 | 0.005586 | 0.195586 |
| 354.7563833 | 81.01031636 | -1.1614 | 7.340745 | 0.006741 | 0.195586 |
| 356.6176488 | 125.7359692 | 0.366494 | 11.54134 | 0.000681 | 0.117812 |
| 356.7533462 | 81.84290422 | -1.13973 | 7.480983 | 0.006235 | 0.195586 |
| 357.2608093 | 60.03956174 | -0.78279 | 12.72277 | 0.000361 | 0.0895 |
| 359.8458876 | 71.55486461 | -0.17962 | 8.519554 | 0.003514 | 0.192905 |
| 364.7859181 | 81.51307727 | -1.0083 | 6.850698 | 0.008861 | 0.195586 |
| 370.2931186 | 94.02020521 | -0.52735 | 6.981196 | 0.008237 | 0.195586 |
| 370.5312595 | 133.9918794 | 0.126721 | 7.169455 | 0.007416 | 0.195586 |
| 371.2988144 | 101.8815654 | -0.1229 | 7.467946 | 0.006281 | 0.195586 |
| 373.0714194 | 137.3284463 | 0.103094 | 6.521857 | 0.010656 | 0.199227 |
| 373.2565648 | 38.09158829 | -0.90694 | 9.24945 | 0.002356 | 0.180595 |
| 378.0530918 | 143.1527373 | 0.109571 | 7.3046 | 0.006878 | 0.195586 |
| 380.2990391 | 44.26024731 | -0.94622 | 6.793562 | 0.009149 | 0.196469 |
| 381.0296735 | 161.4936337 | -0.26908 | 17.92009 | 2.30E-05 | 0.020552 |
| 381.2036278 | 570.8842068 | 0.171519 | 6.722352 | 0.009521 | 0.196956 |
| 382.7564278 | 95.62857376 | -0.7483 | 6.473788 | 0.010948 | 0.1996 |
| 389.9741378 | 162.4970743 | -0.82296 | 6.695404 | 0.009666 | 0.196956 |
| 390.2822909 | 31.83341357 | -0.96362 | 7.428506 | 0.00642 | 0.195586 |
| 390.8885411 | 86.54642273 | 0.137755 | 8.500035 | 0.003551 | 0.192905 |

| | | | | | |
|---|---|---|---|---|---|
| 392.9155012 | 114.1106668 | -0.14719 | 12.26897 | 0.000461 | 0.098411 |
| 394.9800878 | 108.9782225 | -0.20994 | 10.80123 | 0.001014 | 0.139993 |
| 395.2767516 | 116.9384376 | -0.16135 | 6.456386 | 0.011055 | 0.199785 |
| 396.7409049 | 72.30867547 | -1.40146 | 7.986491 | 0.004713 | 0.192905 |
| 398.3250667 | 94.42134708 | -0.32034 | 6.633018 | 0.010011 | 0.199227 |
| 398.7379876 | 72.9237705 | -1.25202 | 8.878524 | 0.002885 | 0.192156 |
| 404.8190314 | 67.82285432 | -0.51357 | 16.77463 | 4.21E-05 | 0.026421 |
| 407.1072447 | 86.78484033 | 0.285799 | 8.381107 | 0.003791 | 0.192905 |
| 408.7671166 | 67.79845761 | -1.18343 | 7.824044 | 0.005156 | 0.194197 |
| 416.0715099 | 551.7610947 | -0.95273 | 8.452509 | 0.003645 | 0.192905 |
| 416.7977375 | 70.29652548 | -0.96285 | 6.591312 | 0.010248 | 0.199227 |
| 418.7084206 | 80.15757685 | -0.23588 | 15.366 | 8.86E-05 | 0.044474 |
| 418.798016 | 67.76896334 | -0.99917 | 7.089689 | 0.007753 | 0.195586 |
| 420.7939857 | 63.64287804 | -1.21996 | 9.001769 | 0.002697 | 0.192156 |
| 422.32374 | 98.54327069 | -0.14715 | 11.97948 | 0.000538 | 0.105681 |
| 422.9264124 | 87.1407763 | 0.123579 | 9.259529 | 0.002343 | 0.180595 |
| 424.7408178 | 78.78632649 | -1.01664 | 6.473274 | 0.010951 | 0.1996 |
| 426.1120457 | 551.6054329 | -0.19112 | 10.40186 | 0.001259 | 0.149944 |
| 426.1904122 | 81.15217561 | 0.177757 | 7.241979 | 0.007122 | 0.195586 |
| 426.7378746 | 77.63257861 | -1.11095 | 6.835108 | 0.008938 | 0.195586 |
| 427.8306865 | 591.7721025 | -0.1818 | 8.174702 | 0.004248 | 0.192905 |
| 429.0204221 | 127.9186668 | 0.167614 | 10.38698 | 0.001269 | 0.149944 |
| 430.1072218 | 552.2998048 | -0.23402 | 8.029494 | 0.004602 | 0.192905 |
| 431.1211128 | 340.1238828 | 0.080288 | 6.538288 | 0.010558 | 0.199227 |
| 437.7786809 | 594.9362016 | -0.1208 | 7.917696 | 0.004895 | 0.192905 |
| 438.3433232 | 41.19973529 | -0.14824 | 8.722541 | 0.003143 | 0.192905 |
| 445.2550861 | 60.10405199 | -0.7147 | 7.801455 | 0.00522 | 0.194197 |
| 446.2627744 | 54.50234297 | -0.94299 | 7.717497 | 0.005469 | 0.195586 |
| 448.3471594 | 53.92532172 | -0.97005 | 7.617172 | 0.005782 | 0.195586 |
| 448.3492635 | 590.0525543 | -1.08934 | 8.000937 | 0.004675 | 0.192905 |
| 449.3262478 | 595.1905 | -0.34247 | 7.372151 | 0.006624 | 0.195586 |
| 454.2945842 | 44.55559685 | -0.84252 | 8.071439 | 0.004497 | 0.192905 |
| 456.3687223 | 32.34447012 | -0.13737 | 6.488575 | 0.010857 | 0.199576 |
| 456.6959023 | 78.0500582 | -1.35753 | 7.471254 | 0.006269 | 0.195586 |
| 458.8774378 | 78.62284431 | 0.129365 | 6.784225 | 0.009197 | 0.196469 |
| 459.7903389 | 68.75011847 | -0.09186 | 6.859643 | 0.008816 | 0.195586 |
| 461.3595365 | 41.13194072 | -1.00434 | 7.126571 | 0.007595 | 0.195586 |
| 463.2774269 | 113.262385 | -0.18409 | 8.944192 | 0.002784 | 0.192156 |
| 463.2996635 | 69.78674841 | -0.66472 | 11.20908 | 0.000814 | 0.125687 |
| 464.7282385 | 74.9608594 | -1.35743 | 9.466264 | 0.002093 | 0.172222 |
| 466.7250088 | 73.97964883 | -1.30269 | 8.165163 | 0.00427 | 0.192905 |

| | | | | | |
|---|---|---|---|---|---|
| 468.2804997 | 58.90236576 | -0.51228 | 11.31642 | 0.000768 | 0.124444 |
| 468.3517877 | 45.86922256 | -0.42308 | 13.58291 | 0.000228 | 0.069465 |
| 474.755907 | 72.86563359 | -1.1663 | 7.940436 | 0.004834 | 0.192905 |
| 476.7537024 | 71.52365572 | -1.14256 | 7.515058 | 0.006119 | 0.195586 |
| 479.2925402 | 577.055641 | -0.71655 | 6.463778 | 0.01101 | 0.199785 |
| 481.2244215 | 74.24474393 | 0.174207 | 9.628872 | 0.001915 | 0.166027 |
| 482.698954 | 78.71722873 | -1.18446 | 6.848269 | 0.008873 | 0.195586 |
| 485.300476 | 88.83925436 | -0.82993 | 7.926906 | 0.004871 | 0.192905 |
| 486.1132201 | 222.651888 | 0.097426 | 6.70289 | 0.009626 | 0.196956 |
| 488.781739 | 69.25192201 | -1.07583 | 7.162784 | 0.007443 | 0.195586 |
| 489.191351 | 117.3073487 | 0.144818 | 9.877916 | 0.001673 | 0.156484 |
| 490.2874821 | 54.44269141 | -0.96867 | 7.018987 | 0.008065 | 0.195586 |
| 490.3893771 | 338.7669783 | 0.094302 | 6.729014 | 0.009486 | 0.196956 |
| 498.3440045 | 23.81070263 | -0.79268 | 6.523642 | 0.010645 | 0.199227 |
| 501.8060627 | 55.70304115 | 0.242255 | 6.508966 | 0.010733 | 0.199227 |
| 502.2926247 | 574.9548878 | -0.81685 | 6.836463 | 0.008932 | 0.195586 |
| 502.337472 | 123.7777773 | 0.200632 | 6.608943 | 0.010147 | 0.199227 |
| 503.2911034 | 572.812096 | -0.84175 | 8.088496 | 0.004455 | 0.192905 |
| 504.2973727 | 575.0346666 | -0.65266 | 7.373498 | 0.006619 | 0.195586 |
| 504.3434483 | 572.6349251 | 0.09567 | 8.091866 | 0.004446 | 0.192905 |
| 507.8521639 | 39.10437109 | -0.18847 | 7.771715 | 0.005307 | 0.194197 |
| 507.9970409 | 146.4312164 | 0.091621 | 7.106362 | 0.007681 | 0.195586 |
| 509.1750864 | 108.7330993 | 0.13 | 7.714778 | 0.005477 | 0.195586 |
| 510.7885313 | 75.26859788 | -0.85463 | 7.615616 | 0.005786 | 0.195586 |
| 512.3354271 | 160.9839197 | -1.02456 | 10.9908 | 0.000916 | 0.13137 |
| 512.3367503 | 471.8457715 | -0.97578 | 9.824191 | 0.001722 | 0.156484 |
| 512.3351118 | 579.8249896 | -0.81467 | 6.933393 | 0.00846 | 0.195586 |
| 514.8736078 | 73.22940033 | -0.96014 | 8.705286 | 0.003173 | 0.192905 |
| 515.2615254 | 115.8493383 | 0.241502 | 7.790248 | 0.005253 | 0.194197 |
| 518.3237478 | 52.92004125 | -0.85158 | 7.318762 | 0.006824 | 0.195586 |
| 520.436837 | 35.22151121 | -0.26396 | 10.76604 | 0.001034 | 0.140129 |
| 523.3558417 | 580.9090811 | -0.6067 | 6.554597 | 0.010461 | 0.199227 |
| 526.3753584 | 21.43073499 | -0.88852 | 6.959851 | 0.008336 | 0.195586 |
| 528.3064111 | 583.7067925 | -0.82102 | 7.23347 | 0.007156 | 0.195586 |
| 532.1094849 | 306.2955926 | 0.1134 | 11.03709 | 0.000893 | 0.13137 |
| 532.7154886 | 74.38014231 | -1.12281 | 7.12132 | 0.007617 | 0.195586 |
| 534.3199215 | 73.42406128 | -0.59913 | 6.602366 | 0.010184 | 0.199227 |
| 534.7117435 | 74.67324329 | -1.15531 | 6.87713 | 0.008731 | 0.195586 |
| 535.8379397 | 201.920608 | -0.09385 | 6.584571 | 0.010287 | 0.199227 |
| 536.30532 | 555.0254281 | 0.10224 | 9.474081 | 0.002084 | 0.172222 |
| 536.8396866 | 24.70235032 | -0.11904 | 6.758099 | 0.009332 | 0.196956 |

| | | | | | |
|---|---|---|---|---|---|
| 539.8025905 | 66.33019132 | 0.186149 | 11.01382 | 0.000904 | 0.13137 |
| 540.3632104 | 523.4517868 | -0.91559 | 8.637853 | 0.003292 | 0.192905 |
| 540.9052475 | 71.52445983 | 0.143358 | 9.997881 | 0.001567 | 0.156484 |
| 541.3671026 | 524.7542288 | -0.6698 | 6.492335 | 0.010834 | 0.199576 |
| 542.3206338 | 56.37072681 | -0.24619 | 25.31023 | 4.88E-07 | 0.002451 |
| 542.319745 | 569.8841912 | -0.54869 | 7.245498 | 0.007108 | 0.195586 |
| 545.3384736 | 589.1572207 | -0.41379 | 6.616415 | 0.010104 | 0.199227 |
| 546.3510458 | 68.18654827 | -0.95549 | 8.554801 | 0.003446 | 0.192905 |
| 554.7707887 | 70.22823482 | -0.95797 | 7.096879 | 0.007722 | 0.195586 |
| 557.097268 | 110.9601551 | -0.73469 | 6.706624 | 0.009606 | 0.196956 |
| 558.7187718 | 75.80337807 | -1.02378 | 7.12806 | 0.007589 | 0.195586 |
| 562.9096546 | 147.4223493 | -0.09896 | 6.87453 | 0.008743 | 0.195586 |
| 566.8853849 | 62.44197786 | -0.81049 | 6.486365 | 0.010871 | 0.199576 |
| 568.7477403 | 76.26669466 | -0.81698 | 11.94757 | 0.000547 | 0.105681 |
| 570.3658676 | 111.5643221 | -0.51468 | 7.049289 | 0.00793 | 0.195586 |
| 573.271284 | 134.2022333 | 0.186605 | 12.71503 | 0.000363 | 0.0895 |
| 574.7962358 | 73.12710961 | 0.102353 | 7.303302 | 0.006883 | 0.195586 |
| 578.7777991 | 77.97414699 | -0.80588 | 6.989025 | 0.008201 | 0.195586 |
| 579.0754841 | 143.6831327 | 0.1409 | 12.84499 | 0.000338 | 0.0895 |
| 583.1060977 | 348.5503737 | -0.13112 | 7.903741 | 0.004933 | 0.192905 |
| 583.3425604 | 88.00795362 | 0.360004 | 24.09796 | 9.16E-07 | 0.003065 |
| 586.1924468 | 146.7020934 | 0.132571 | 8.26608 | 0.004039 | 0.192905 |
| 588.8075899 | 86.66134479 | -0.37244 | 14.90265 | 0.000113 | 0.045476 |
| 598.8005087 | 72.82620638 | 0.191144 | 12.63772 | 0.000378 | 0.0895 |
| 599.4311083 | 79.98119913 | 0.120552 | 8.117344 | 0.004384 | 0.192905 |
| 604.6984813 | 69.69789891 | -0.61032 | 12.08992 | 0.000507 | 0.103905 |
| 605.8983878 | 59.13748362 | -0.9101 | 19.73091 | 8.91E-06 | 0.017906 |
| 608.6462058 | 74.99812442 | -1.19744 | 6.892729 | 0.008655 | 0.195586 |
| 613.377823 | 393.0807875 | 0.135124 | 10.65917 | 0.001095 | 0.141281 |
| 615.8823263 | 100.3060927 | -0.49409 | 8.405519 | 0.003741 | 0.192905 |
| 618.6745576 | 75.13445193 | -1.21065 | 8.570671 | 0.003416 | 0.192905 |
| 620.7624044 | 69.89481878 | -0.86007 | 6.680954 | 0.009745 | 0.197313 |
| 621.182736 | 353.6653727 | -0.14523 | 6.503913 | 0.010764 | 0.199227 |
| 625.2098164 | 562.1997992 | -0.09972 | 7.293343 | 0.006921 | 0.195586 |
| 628.8135498 | 66.25371864 | -0.11537 | 6.722285 | 0.009522 | 0.196956 |
| 632.6548174 | 80.63983943 | -0.58188 | 7.45249 | 0.006335 | 0.195586 |
| 641.1083734 | 162.0402769 | 0.148464 | 8.555003 | 0.003446 | 0.192905 |
| 642.8424759 | 66.41287276 | -0.16027 | 10.74349 | 0.001046 | 0.140129 |
| 646.1284449 | 149.8049532 | 0.136612 | 7.291114 | 0.00693 | 0.195586 |
| 655.3953058 | 531.1244669 | 0.231681 | 15.05329 | 0.000105 | 0.045476 |
| 658.7567555 | 72.0169231 | 0.148862 | 6.699776 | 0.009643 | 0.196956 |

| | | | | | |
|---|---|---|---|---|---|
| 660.4877713 | 41.95159006 | -0.44935 | 7.967157 | 0.004763 | 0.192905 |
| 660.8445875 | 71.36401447 | 0.145683 | 16.50406 | 4.85E-05 | 0.028679 |
| 662.0900634 | 143.8873385 | 0.100572 | 6.844772 | 0.00889 | 0.195586 |
| 663.4370047 | 51.2302181 | -0.71863 | 7.618416 | 0.005778 | 0.195586 |
| 665.4438388 | 596.9650051 | -0.15742 | 17.06973 | 3.60E-05 | 0.024125 |
| 666.4167606 | 68.07325663 | -0.12085 | 6.808027 | 0.009075 | 0.195998 |
| 675.0812578 | 160.584026 | 0.171262 | 7.048571 | 0.007933 | 0.195586 |
| 682.4237292 | 593.1798414 | -0.40544 | 8.459531 | 0.003631 | 0.192905 |
| 685.4342448 | 55.02744042 | -0.88117 | 6.827219 | 0.008978 | 0.195586 |
| 687.1117539 | 390.8988534 | 0.143318 | 12.46005 | 0.000416 | 0.092786 |
| 688.4818963 | 44.58083739 | -0.95805 | 8.267768 | 0.004036 | 0.192905 |
| 692.0168251 | 154.8198366 | 0.105994 | 7.949347 | 0.00481 | 0.192905 |
| 692.6106518 | 78.91029122 | -0.66285 | 6.811164 | 0.009059 | 0.195998 |
| 692.8323627 | 63.16597004 | -1.00316 | 6.692001 | 0.009685 | 0.196956 |
| 696.0589115 | 161.0914148 | 0.142254 | 9.870999 | 0.001679 | 0.156484 |
| 698.6903813 | 76.29577917 | -0.98525 | 7.388602 | 0.006564 | 0.195586 |
| 699.5987277 | 336.9100379 | -0.15592 | 7.308213 | 0.006864 | 0.195586 |
| 704.5764774 | 24.7604849 | -0.47216 | 9.61761 | 0.001927 | 0.166027 |
| 704.5779184 | 591.0342987 | -0.61855 | 11.85747 | 0.000574 | 0.10681 |
| 721.4686191 | 441.9595082 | -0.1343 | 7.023979 | 0.008043 | 0.195586 |
| 724.7942782 | 79.22665494 | -0.16353 | 9.639077 | 0.001905 | 0.166027 |
| 730.5354266 | 27.81005161 | -0.29992 | 8.181919 | 0.004231 | 0.192905 |
| 731.5395754 | 28.13838815 | -0.14841 | 7.083717 | 0.007779 | 0.195586 |
| 732.5523872 | 590.9130684 | -1.05744 | 9.407754 | 0.002161 | 0.173599 |
| 733.2709132 | 138.777077 | 0.281991 | 11.28669 | 0.000781 | 0.124445 |
| 736.0741049 | 143.8340782 | 0.165042 | 9.829818 | 0.001717 | 0.156484 |
| 738.8892048 | 66.32837244 | 0.142302 | 14.59063 | 0.000134 | 0.051597 |
| 739.470792 | 58.0090511 | -0.86328 | 6.458019 | 0.011045 | 0.199785 |
| 740.5284373 | 44.54099601 | -1.08071 | 10.10708 | 0.001477 | 0.156484 |
| 741.8742682 | 58.96055752 | -0.79328 | 9.816497 | 0.00173 | 0.156484 |
| 747.4287653 | 139.3591938 | 0.114606 | 12.61236 | 0.000383 | 0.0895 |
| 753.4397397 | 47.00659339 | -0.93102 | 7.333293 | 0.006769 | 0.195586 |
| 754.6501605 | 76.1668752 | -0.89757 | 7.186105 | 0.007347 | 0.195586 |
| 760.5820225 | 136.195641 | -0.61963 | 7.957541 | 0.004789 | 0.192905 |
| 762.5925136 | 42.57556649 | -0.95233 | 6.671725 | 0.009795 | 0.197541 |
| 763.1810525 | 410.4101873 | -0.15928 | 6.784458 | 0.009195 | 0.196469 |
| 763.5934842 | 584.1271651 | 0.144601 | 7.874651 | 0.005013 | 0.192905 |
| 768.5284771 | 68.82258072 | -0.37178 | 7.904444 | 0.004931 | 0.192905 |
| 769.4910647 | 86.22392949 | -0.60486 | 9.355041 | 0.002224 | 0.17585 |
| 780.5487454 | 75.80191572 | -0.79798 | 7.024493 | 0.00804 | 0.195586 |
| 780.5489869 | 592.9126886 | -0.80548 | 6.976638 | 0.008258 | 0.195586 |

| | | | | | |
|---|---|---|---|---|---|
| 781.5536814 | 72.77506392 | -0.76995 | 6.553223 | 0.010469 | 0.199227 |
| 781.5535438 | 592.4575367 | -0.73141 | 6.541229 | 0.01054 | 0.199227 |
| 783.5724017 | 136.4767158 | -0.70527 | 11.87855 | 0.000568 | 0.10681 |
| 786.5173302 | 70.31584726 | -0.15377 | 8.135171 | 0.004341 | 0.192905 |
| 786.6001327 | 136.8967551 | -0.60402 | 10.69274 | 0.001076 | 0.141281 |
| 788.6057547 | 592.2175612 | -0.79873 | 8.512568 | 0.003527 | 0.192905 |
| 800.6287971 | 69.49312988 | -0.12591 | 8.060523 | 0.004524 | 0.192905 |
| 801.5059375 | 457.9192802 | -0.10307 | 6.691389 | 0.009688 | 0.196956 |
| 802.5750752 | 65.76904823 | -0.61212 | 7.12488 | 0.007602 | 0.195586 |
| 804.5502705 | 77.15289606 | -0.84235 | 7.814305 | 0.005183 | 0.194197 |
| 804.933119 | 437.2668508 | 0.115735 | 7.709733 | 0.005492 | 0.195586 |
| 807.5720539 | 135.6156674 | -0.0824 | 6.449958 | 0.011095 | 0.199785 |
| 808.5829558 | 135.9442924 | -0.71129 | 10.33082 | 0.001308 | 0.152778 |
| 810.5976614 | 44.54289512 | -0.86837 | 7.295326 | 0.006913 | 0.195586 |
| 810.7384425 | 66.60543086 | -0.37285 | 14.0467 | 0.000178 | 0.060768 |
| 813.6186409 | 28.03332954 | -0.60181 | 6.941857 | 0.00842 | 0.195586 |
| 821.5653246 | 43.04572916 | -1.0577 | 8.714202 | 0.003157 | 0.192905 |
| 833.8235651 | 59.42773112 | -0.71082 | 12.76126 | 0.000354 | 0.0895 |
| 844.7426552 | 109.1571963 | 0.124634 | 9.966648 | 0.001594 | 0.156484 |
| 845.7330622 | 446.4562027 | 0.120423 | 8.490551 | 0.00357 | 0.192905 |
| 846.5595957 | 71.36545283 | -0.70075 | 8.200352 | 0.004188 | 0.192905 |
| 851.7438571 | 460.691371 | 0.096195 | 8.630794 | 0.003305 | 0.192905 |
| 854.5935427 | 69.29333304 | -0.93127 | 7.163553 | 0.00744 | 0.195586 |
| 854.8090244 | 64.45936822 | 0.124389 | 9.148056 | 0.00249 | 0.189434 |
| 861.6920106 | 456.7347803 | 0.107189 | 8.087719 | 0.004457 | 0.192905 |
| 877.2754305 | 145.4507299 | 0.083242 | 6.928902 | 0.008481 | 0.195586 |
| 880.1169814 | 450.6372534 | 0.104321 | 10.58483 | 0.00114 | 0.141605 |
| 881.6945533 | 443.6975001 | 0.112127 | 7.265987 | 0.007027 | 0.195586 |
| 883.2601661 | 441.7467182 | 0.083549 | 6.877013 | 0.008731 | 0.195586 |
| 896.7946951 | 59.74769056 | -0.97995 | 7.481772 | 0.006233 | 0.195586 |
| 905.738705 | 459.207443 | 0.097537 | 8.8655 | 0.002906 | 0.192156 |
| 906.5003296 | 470.1044339 | 0.137322 | 8.0879 | 0.004456 | 0.192905 |
| 908.2075394 | 427.3041531 | -0.10531 | 10.38799 | 0.001268 | 0.149944 |
| 913.5696271 | 53.65111137 | -0.88009 | 6.552684 | 0.010473 | 0.199227 |
| 916.5173703 | 57.72705694 | 0.125744 | 6.514502 | 0.0107 | 0.199227 |
| 916.5266211 | 59.47250409 | -0.71097 | 9.79747 | 0.001748 | 0.1567 |
| 919.3593254 | 437.1885782 | 0.074664 | 6.778204 | 0.009228 | 0.196469 |
| 920.2136834 | 442.1848742 | 0.092162 | 6.874064 | 0.008746 | 0.195586 |
| 921.2423711 | 150.4304652 | 0.093699 | 7.274207 | 0.006995 | 0.195586 |
| 926.667673 | 69.62313461 | -0.25015 | 14.9212 | 0.000112 | 0.045476 |
| 929.7582587 | 439.0295944 | 0.094192 | 9.581039 | 0.001966 | 0.167324 |

| | | | | | |
|---|---|---|---|---|---|
| 938.6825713 | 461.2115171 | 0.122097 | 11.36708 | 0.000748 | 0.12308 |
| 940.8175295 | 60.69109091 | -0.74178 | 7.605497 | 0.005819 | 0.195586 |
| 943.2216084 | 157.3469565 | 0.132588 | 12.52831 | 0.000401 | 0.09149 |
| 951.8338497 | 422.5948998 | 0.091245 | 7.54126 | 0.00603 | 0.195586 |
| 967.1777114 | 452.2938121 | 0.111406 | 7.271795 | 0.007005 | 0.195586 |
| 979.5754792 | 572.2565896 | -0.12065 | 6.971969 | 0.00828 | 0.195586 |
| 995.8883416 | 447.9868708 | 0.13753 | 9.985845 | 0.001577 | 0.156484 |
| 1015.845101 | 424.4227761 | 0.146661 | 18.58305 | 1.63E-05 | 0.018294 |
| 1068.517335 | 180.7074107 | -0.14092 | 6.953328 | 0.008366 | 0.195586 |
| 1074.304254 | 454.046443 | 0.094115 | 7.945474 | 0.004821 | 0.192905 |
| 1086.0495 | 148.5673178 | 0.13592 | 13.08862 | 0.000297 | 0.085249 |
| 1086.286683 | 464.7570785 | 0.125012 | 8.159263 | 0.004284 | 0.192905 |
| 1086.554068 | 459.9223156 | 0.118656 | 8.957808 | 0.002763 | 0.192156 |
| 1087.488702 | 438.8357602 | 0.105003 | 6.89037 | 0.008666 | 0.195586 |
| 1091.552695 | 435.2957383 | -0.10055 | 8.847809 | 0.002934 | 0.192616 |
| 1093.567769 | 439.2836091 | 0.092633 | 6.507221 | 0.010744 | 0.199227 |
| 1096.283319 | 437.8806466 | 0.109682 | 10.08995 | 0.001491 | 0.156484 |
| 1096.668819 | 441.7698068 | 0.09362 | 7.125376 | 0.0076 | 0.195586 |
| 1100.75934 | 58.95524154 | -0.99071 | 8.174258 | 0.004249 | 0.192905 |
| 1120.303606 | 443.596767 | 0.119858 | 7.079902 | 0.007795 | 0.195586 |
| 1125.56474 | 433.3198026 | 0.092316 | 6.626461 | 0.010047 | 0.199227 |
| 1145.284603 | 60.79783807 | -0.6728 | 9.621466 | 0.001923 | 0.166027 |
| 1145.502131 | 418.8533682 | 0.095228 | 6.885114 | 0.008692 | 0.195586 |
| 1173.566774 | 445.3123494 | 0.08994 | 8.192473 | 0.004206 | 0.192905 |
| 1173.760997 | 59.31980044 | -0.39782 | 11.43257 | 0.000722 | 0.120797 |
| 1241.63629 | 151.1895873 | 0.126254 | 6.521853 | 0.010656 | 0.199227 |
| 1253.604972 | 440.0430819 | -0.19545 | 9.033763 | 0.00265 | 0.191572 |
| 1257.779343 | 440.8086827 | 0.090686 | 7.948123 | 0.004814 | 0.192905 |

**TABLE S2**: Full list of pathways and related features (M/Z, retention time, ODDS ratio, P-value and FDR correction) and related KEGG ID obtained from the Pathway enrichment analyses.

| pathway | p.value..raw. | p.value | Feature Name | kegg.id | m/z | ret.time | odds ratio | lower | upper | p.val | adj.pval |
|---|---|---|---|---|---|---|---|---|---|---|---|
| **3-Chloroacrylic acid degradation** | 0.0837433021019 | 0.00390993122125 | 3-Chloroallyl aldehyde; trans-3-Chloroallyl aldehyde | C06613 | 174.95721567721 | 349.034959223277 | 0.852521 | 1.5E-05 | 0.916602 | 1.5957411414691e-05 | 0.0182937654350227 |
| **3-Chloroacrylic acid degradation** | 0.0837433021019 | 0.00390993122125 | cis-3-Chloroallyl aldehyde | C16348 | 174.95721567721 | 349.034959223277 | 0.852521 | 1.5E-05 | 0.916602 | 1.5957411414691e-05 | 0.0182937654350227 |
| **Xenobiotics metabolism** | 0.600684633088 | 0.0283528809697 | Bromobenzene-2,3-oxide; Bromobenzene-2,3-epoxide | C14840 | 174.95721567721 | 349.034959223277 | 0.852521 | 1.5E-05 | 0.916602 | 1.5957411414691e-05 | 0.0182937654350227 |
| **Xenobiotics metabolism** | 0.600684633088 | 0.0283528809697 | 4-Bromophenol | C14453 | 174.95721567721 | 349.034959223277 | 0.852521 | 1.5E-05 | 0.916602 | 1.5957411414691e-05 | 0.0182937654350227 |
| **Xenobiotics metabolism** | 0.600684633088 | 0.0283528809697 | | C14841 | 174.95721567721 | 349.034959223277 | 0.852521 | 1.5E-05 | 0.916602 | 1.5957411414691e-05 | 0.0182937654350227 |
| **Xenobiotics metabolism** | 0.600684633088 | 0.0283528809697 | Bromobenzene-3,4-oxide; Bromobenzene-3,4-epoxide | C14839 | 174.95721567721 | 349.034959223277 | 0.852521 | 1.5E-05 | 0.916602 | 1.5957411414691e-05 | 0.0182937654350227 |
| **Drug metabolism - other enzymes** | 0.777441728451 | 0.179119678411 | 6-Thioxanthine 5'-monophosphate | C16618 | 381.029673535904 | 161.493633672044 | 0.92245 | 2.45E-05 | 0.957578 | 2.30376346672667e-05 | 0.0205515689138621 |
| **Fatty Acid Metabolism** | 0.00136264372743 | 0.000339697971178 | Phytanate; Phytanic acid | C01607 | 168.149423477562 | 48.1428754571379 | 1.049066 | 0.001038 | 1.078177 | 0.00060324655087729 | 0.10818580554394 |
| **C21-steroid hormone biosynthesis and metabolism** | 0.997864412275 | 0.655533608793 | Sterol | C00370 | 250.224262076074 | 488.878549975896 | 0.925344 | 3.41E-05 | 0.959743 | 3.09361595268731e-05 | 0.0221922750091705 |

| | | | | | | | | | | | |
|---|---|---|---|---|---|---|---|---|---|---|---|
| **Vitamin B5 - CoA biosynthesis from pantothenate** | 0.750902553927 | 0.999986263096 | D-4'-Phosphopantothenate; (R)-4'-Phosphopantothenate | C03492 | 338.037746150498 | 171.901587356365 | 0.906402 | 0.000208 | 0.953847 | 0.000159863055168241 | 0.0594631356686906 |
| **Vitamin K metabolism** | 0.340311783887 | 0.999986263096 | Vitamin K1 epoxide; (2,3-Epoxyphytyl)menaquinone; 2,3-Epoxy-2,3-dihydro-2-methyl-3-phytyl-1,4-naphthoquinone; 2,3-Epoxyphylloquinone; 1a,7a-Dihydro-1a-methyl-7a-(3,7,11,15-tetramethyl-2-hexadecenyl)-naphth[2,3-b]oxirene-2,7-dione; Phylloquinone oxide; Phylloquinone epoxide; Phylloquinone-2,3-epoxide; Vitamin K1 2,3-epoxide; Vitamin K1 oxide | C05849 | 468.351787724214 | 45.8692225611401 | 0.880802 | 0.000299 | 0.942308 | 0.000228253552651214 | 0.0694651645235195 |
| **Carnitine shuttle** | 0.0831103651174 | 0.000874817459984 | gamma-linolenyl carnitine | lnlncgcrn | 422.323740028004 | 98.5432706882734 | 0.956816 | 0.000835 | 0.981036 | 0.000537896974246888 | 0.105680785606795 |
| **Carnitine shuttle** | 0.0831103651174 | 0.000874817459984 | alpha-linolenyl carnitine | lnlncacrn | 422.323740028004 | 98.5432706882734 | 0.956816 | 0.000835 | 0.981036 | 0.000537896974246888 | 0.105680785606795 |
| **Carnitine shuttle** | 0.0831103651174 | 0.000874817459984 | L-Palmitoylcarnitine | C02990 | 422.323740028004 | 98.5432706882734 | 0.956816 | 0.000835 | 0.981036 | 0.000537896974246888 | 0.105680785606795 |
| **Saturated fatty acids beta-oxidation** | 0.182491247164 | 0.00237820952794 | L-Palmitoylcarnitine | C02990 | 422.323740028004 | 98.5432706882734 | 0.956816 | 0.000835 | 0.981036 | 0.000537896974246888 | 0.105680785606795 |
| **Fatty Acid Metabolism** | 0.00136264372743 | 0.000339697971178 | O-Acetylcarnitine; O-Acetyl-L-carnitine | C02571 | 204.122297531216 | 91.1524890948529 | 0.751563 | 0.004096 | 0.907027 | 0.00290826579096826 | 0.192156008807199 |
| **De novo fatty acid biosynthesis** | 0.104709729668 | 0.00101148958738 | Icosanoic acid; Eicosanoic acid; Arachidic acid | C06425 | 168.149423477562 | 48.1428754571379 | 1.049066 | 0.001038 | 1.078177 | 0.00060324655087729 | 0.10818580554394 |
| **Phytanic acid peroxisomal oxidation** | 0.305493893555 | 0.0113522941772 | Phytanate; Phytanic acid | C01607 | 168.149423477562 | 48.1428754571379 | 1.049066 | 0.001038 | 1.078177 | 0.00060324655087729 | 0.10818580554394 |
| **Fatty acid activation** | 0.481162063704 | 0.0199323827058 | Icosanoic acid; Eicosanoic acid; Arachidic acid | C06425 | 168.149423477562 | 48.1428754571379 | 1.049066 | 0.001038 | 1.078177 | 0.00060324655087729 | 0.10818580554394 |

| | | | | | | | | | | | |
|---|---|---|---|---|---|---|---|---|---|---|---|
| **Fatty acid activation** | 0.481162063704 | 0.0199323827058 | Phytanate; Phytanic acid | C01607 | 168.149423477562 | 48.1428754571379 | 1.049066 | 0.001038 | 1.078177 | 0.00060324655087729 | 0.10818580554394 |
| **Urea cycle/amino group metabolism** | 0.221709062926 | 0.00252163988827 | N-Acetyl-L-glutamate 5-phosphate; N-Acetyl-L-glutamyl 5-phosphate | C04133 | 271.038368796265 | 88.4284262049476 | 0.697094 | 0.000716 | 0.857185 | 0.000623886237959614 | 0.109924376979446 |
| **Vitamin E metabolism** | 0.50735458165 | 0.023187298019 | alpha-Tocotrienol | C14153 | 463.299663471208 | 69.7867484051683 | 0.81921 | 0.001125 | 0.920653 | 0.000813982784083511 | 0.125687336353801 |
| **Ascorbate (Vitamin C) and Aldarate Metabolism** | 0.918621397261 | 0.999986263096 | alpha-Tocotrienol | C14153 | 463.299663471208 | 69.7867484051683 | 0.81921 | 0.001125 | 0.920653 | 0.000813982784083511 | 0.125687336353801 |
| **Urea cycle/amino group metabolism** | 0.221709062926 | 0.00252163988827 | Spermine; N,N'-Bis(3-aminopropyl)-1,4-butanediamine | C00750 | 241.178690927244 | 45.8258062949099 | 0.833506 | 0.001459 | 0.929158 | 0.001017571014582 | 0.139992680814342 |
| **Beta-Alanine metabolism** | 0.231059797676 | 0.00680777802656 | Spermine; N,N'-Bis(3-aminopropyl)-1,4-butanediamine | C00750 | 241.178690927244 | 45.8258062949099 | 0.833506 | 0.001459 | 0.929158 | 0.001017571014582 | 0.139992680814342 |
| **Aspartate and asparagine metabolism** | 0.822873368732 | 0.123478035657 | Spermine; N,N'-Bis(3-aminopropyl)-1,4-butanediamine | C00750 | 241.178690927244 | 45.8258062949099 | 0.833506 | 0.001459 | 0.929158 | 0.001017571014582 | 0.139992680814342 |
| **Beta-Alanine metabolism** | 0.231059797676 | 0.00680777802656 | L-Histidine; (S)-alpha-Amino-1H-imidazole-4-propionic acid | C00135 | 178.058752484365 | 128.838894614929 | 0.746282 | 0.001463 | 0.890033 | 0.00112913618094368 | 0.141605435508952 |
| **Phytanic acid peroxisomal oxidation** | 0.305493893555 | 0.0113522941772 | 2(R)-pristanal | CE5127 | 285.297658676797 | 586.99527009136 | 0.681313 | 0.001352 | 0.858539 | 0.00114209302760382 | 0.141605435508952 |
| **Phytanic acid peroxisomal oxidation** | 0.305493893555 | 0.0113522941772 | 2(S)-pristanal | CE5124 | 285.297658676797 | 586.99527009136 | 0.681313 | 0.001352 | 0.858539 | 0.00114209302760382 | 0.141605435508952 |
| **Histidine metabolism** | 0.69820367706 | 0.130985025415 | L-Histidine; (S)-alpha-Amino-1H-imidazole-4-propionic acid | C00135 | 178.058752484365 | 128.838894614929 | 0.746282 | 0.001463 | 0.890033 | 0.00112913618094368 | 0.141605435508952 |
| **Hexose phosphorylation** | 0.0877751 | 0.00117065 | IDP; Inosine 5'-diphosphate; Inosine diphosphate | C00104 | 429.020422096477 | 127.918666816868 | 1.05157 | 0.002341 | 1.084224 | 0.0012690 | 0.149944102146144 |

| | | | | | | | | | | | |
|---|---|---|---|---|---|---|---|---|---|---|---|
| | 590204 | 111852 | | | | | | | | 6787637382 | |
| **Saturated fatty acids beta-oxidation** | 0.182491247164 | 0.002378209527 94 | 3-Oxooctanoyl-CoA | C05267 | 908.207539412075 | 427.30415312774 | 0.968901 | 0.002156 | 0.987696 | 0.00126837557352133 | 0.149944102146144 |
| **Aminosugars metabolism** | 0.745836911495 | 0.115588027425 | IDP; Inosine 5'-diphosphate; Inosine diphosphate | C00104 | 429.020422096477 | 127.918666816868 | 1.05157 | 0.002341 | 1.084224 | 0.00126906787637382 | 0.149944102146144 |
| **Galactose metabolism** | 0.821437186772 | 0.166638916907 | IDP; Inosine 5'-diphosphate; Inosine diphosphate | C00104 | 429.020422096477 | 127.918666816868 | 1.05157 | 0.002341 | 1.084224 | 0.00126906787637382 | 0.149944102146144 |
| **Pyrimidine metabolism** | 0.92475584473 | 0.292941294268 | IDP; Inosine 5'-diphosphate; Inosine diphosphate | C00104 | 429.020422096477 | 127.918666816868 | 1.05157 | 0.002341 | 1.084224 | 0.00126906787637382 | 0.149944102146144 |
| **Purine metabolism** | 0.987538751028 | 0.549295313914 | IDP; Inosine 5'-diphosphate; Inosine diphosphate | C00104 | 429.020422096477 | 127.918666816868 | 1.05157 | 0.002341 | 1.084224 | 0.00126906787637382 | 0.149944102146144 |
| **TCA cycle** | 0.93853450406 | 0.999986263096 | IDP; Inosine 5'-diphosphate; Inosine diphosphate | C00104 | 429.020422096477 | 127.918666816868 | 1.05157 | 0.002341 | 1.084224 | 0.00126906787637382 | 0.149944102146144 |
| **Hexose phosphorylation** | 0.0877751590204 | 0.00117065111852 | Sorbitol 6-phosphate; D-Sorbitol 6-phosphate; D-Glucitol 6-phosphate | C01096 | 132.531319235311 | 213.542658388674 | 0.958012 | 0.002966 | 0.984055 | 0.00172095165383301 | 0.156483926095199 |
| **Drug metabolism - cytochrome P450** | 0.339766036825 | 0.00657261942317 | Carbamazepine | C06868 | 236.095159512481 | 145.749708458655 | 1.040422 | 0.002683 | 1.066114 | 0.00145321222058836 | 0.156483926095199 |
| **Glycine, serine, alanine and threonine metabolism** | 0.518583298632 | 0.0199044796148 | L-2-Amino-3-oxobutanoic acid; L-2-Amino-3-oxobutanoate; L-2-Amino-acetoacetate; (S)-2-Amino-3-oxobutanoic acid | C03508 | 120.044482578797 | 441.402262862449 | 0.943967 | 0.002679 | 0.97836 | 0.00158757923416941 | 0.156483926095199 |
| **Glycolysis and Gluconeogenesis** | 0.764250008433 | 0.106764598315 | Acetyl phosphate | C00227 | 140.995920790709 | 31.6615376938635 | 1.08914 | 0.003323 | 1.148793 | 0.00169748084582089 | 0.156483926095199 |

| | | | | | | | | | | | |
|---|---|---|---|---|---|---|---|---|---|---|---|
| **Valine, leucine and isoleucine degradation** | 0.86642965905 8 | 0.185929680637 | 3-Methyl-2-oxobutanoic acid; 3-Methyl-2-oxobutyric acid; 3-Methyl-2-oxobutanoate; 2-Oxo-3-methylbutanoate; 2-Oxoisovalerate; 2-Oxoisopentanoate; alpha-Ketovaline; 2-Ketovaline; 2-Keto-3-methylbutyric acid | C00141 | 119.049382903908 | 113.522534613914 | 0.881505 | 0.002499 | 0.953271 | 0.00158634205722552 | 0.156483926095199 |
| **Sialic acid metabolism** | 0.88320246975 8 | 0.279395532525 | D-Galactosyl-3-(N-acetyl-beta-D-galactosaminyl)-L-serine | C04776 | 236.095159512481 | 145.749708458655 | 1.040422 | 0.002683 | 1.066114 | 0.00145321222058836 | 0.156483926095199 |
| **Methionine and cysteine metabolism** | 0.87740544892 2 | 0.198255230658 | L-Cysteate; L-Cysteic acid; 3-Sulfoalanine; 2-Amino-3-sulfopropionic acid | C00506 | 171.013845858458 | 55.234334477129 | 0.908593 | 0.003192 | 0.96536 | 0.0019342022863863 | 0.166027295403227 |
| **Androgen and estrogen biosynthesis and metabolism** | 0.98779638849 3 | 0.510935384898 | 2-Methoxy-estradiol-17beta 3-glucuronide | C11131 | 481.224421529167 | 74.2447439315526 | 1.053652 | 0.003663 | 1.089014 | 0.00191542220248631 | 0.166027295403227 |
| **Fatty Acid Metabolism** | 0.00136264372743 | 0.0003396979711 78 | Linoleate; Linoleic acid; (9Z,12Z)-Octadecadienoic acid; 9-cis,12-cis-Octadecadienoate; 9-cis,12-cis-Octadecadienoic acid | C01595 | 281.24752253069 | 33.784067698132 | 0.742706 | 0.006638 | 0.912449 | 0.00461916697109899 | 0.19290463114884 |
| **Lysine metabolism** | 0.12545355524 7 | 0.0014816264063 | 4-Trimethylammoniobutanal | C01149 | 169.086005572512 | 464.273075221709 | 0.705646 | 0.002568 | 0.880268 | 0.00199857948558313 | 0.16867003171186 |
| **Urea cycle/amino group metabolism** | 0.22170906292 6 | 0.00252163988827 | Agmatine; (4-Aminobutyl) guanidine | C00179 | 169.086005572512 | 464.273075221709 | 0.705646 | 0.002568 | 0.880268 | 0.00199857948558313 | 0.16867003171186 |
| **Aminosugars metabolism** | 0.74583691149 5 | 0.115588027425 | N-Trimethyl-2-aminoethylphosphonate; 2-Trimethylaminoethylphosphonate | C06459 | 169.086005572512 | 464.273075221709 | 0.705646 | 0.002568 | 0.880268 | 0.00199857948558313 | 0.16867003171186 |
| **Leukotriene metabolism** | 0.88759672329 5 | 0.210733231746 | omega-carboxy-trinor-leukotriene B4 | CE6185 | 169.086005572512 | 464.273075221709 | 0.705646 | 0.002568 | 0.880268 | 0.00199857948558313 | 0.16867003171186 |
| **Arginine and Proline Metabolism** | 0.87707836381 5 | 0.221999947583 | Agmatine; (4-Aminobutyl) guanidine | C00179 | 169.086005572512 | 464.273075221709 | 0.705646 | 0.002568 | 0.880268 | 0.00199857948558313 | 0.16867003171186 |
| **Fatty Acid Metabolism** | 0.00136264372743 | 0.0003396979711 78 | octadecenoate (n-C18:1) | ocdcea | 281.24752253069 | 33.784067698132 | 0.742706 | 0.006638 | 0.912449 | 0.00461916697 | 0.19290463114884 |

| | | | | | | | | | | | |
|---|---|---|---|---|---|---|---|---|---|---|---|
| | | | | | | | | | | 109899 | |
| **3-Chloroacrylic acid degradation** | 0.0837433021019 | 0.003909931221 25 | trans-3-Chloro-2-propene-1-ol | C06611 | 176.97187480322 | 328.360259282249 | 0.757677 | 0.003657 | 0.907638 | 0.00259702374368997 | 0.191571615555352 |
| **3-Chloroacrylic acid degradation** | 0.0837433021019 | 0.00390993122125 | cis-3-Chloro-2-propene-1-ol | C06612 | 176.97187480322 | 328.360259282249 | 0.757677 | 0.003657 | 0.907638 | 0.00259702374368997 | 0.191571615555352 |
| **Fatty Acid Metabolism** | 0.00136264372743 | 0.0003396979711 78 | L-Carnitine; L-gamma-Trimethyl-beta-hydroxybutyrobetaine; Vitamin BT; 3-Carboxy-2-hydroxy-N,N,N-trimethyl-1-propanaminium hydroxide, inner salt; Levocarnitine; (R)-Carnitine | C00318 | 163.115801406981 | 88.5776592293596 | 0.732694 | 0.007677 | 0.912029 | 0.00536372015369579 | 0.194468741890133 |
| **Fatty Acid Metabolism** | 0.00136264372743 | 0.000339697971178 | Tetradecanoic acid; Tetradecanoate; Myristic acid | C06424 | 251.197160068562 | 128.325061638681 | 0.957787 | 0.013649 | 0.988378 | 0.00717150534694222 | 0.195585714469996 |
| **Fatty Acid Metabolism** | 0.00136264372743 | 0.000339697971178 | Decanoyl-CoA | C05274 | 921.24237107895 | 150.430465242413 | 1.028509 | 0.014277 | 1.049735 | 0.00699517578790609 | 0.195585714469996 |
| **Fatty Acid Metabolism** | 0.00136264372743 | 0.000339697971178 | Hexadecanoic acid; Hexadecanoate; Hexadecylic acid; Palmitic acid; Palmitate; Cetylic acid | C00249 | 279.231911695023 | 28.1891724631974 | 0.739227 | 0.009696 | 0.919419 | 0.0066300345659458 | 0.195585714469996 |
| **Hexose phosphorylation** | 0.0877751590204 | 0.001170651118 52 | D-Mannose; Mannose; Seminose; Carubinose | C00159 | 203.05373260968 | 74.2978463704007 | 0.707545 | 0.003846 | 0.888464 | 0.00290126728766105 | 0.192156008807199 |
| **Hexose phosphorylation** | 0.0877751590204 | 0.00117065111852 | D-Glucose; Grape sugar; Dextrose | C00031 | 203.05373260968 | 74.2978463704007 | 0.707545 | 0.003846 | 0.888464 | 0.00290126728766105 | 0.192156008807199 |
| **Hexose phosphorylation** | 0.0877751590204 | 0.00117065111852 | D-Hexose | C00738 | 203.05373260968 | 74.2978463704007 | 0.707545 | 0.003846 | 0.888464 | 0.00290126728766105 | 0.192156008807199 |
| **Hexose phosphorylation** | 0.0877751590204 | 0.00117065111852 | D-Fructose; Levulose; Fruit sugar; D-arabino-Hexulose | C00095 | 203.05373260968 | 74.2978463704007 | 0.707545 | 0.003846 | 0.888464 | 0.00290126728766105 | 0.192156008807199 |
| **Glycosphingolipid metabolism** | 0.2915437 | 0.00607553 | D-Galactose | C00124 | 203.05373260968 | 74.2978463704007 | 0.707545 | 0.003846 | 0.888464 | 0.0029012 | 0.192156008807199 |

| | | | | | | | | | | | |
|---|---|---|---|---|---|---|---|---|---|---|---|
| | 46312 | 428773 | | | | | | | | 6728766105 | |
| **Glycosphingolipid metabolism** | 0.291543746312 | 0.00607553428773 | D-Glucose; Grape sugar; Dextrose | C00031 | 203.05373260968 | 74.2978463704007 | 0.707545 | 0.003846 | 0.888464 | 0.002901267287661 05 | 0.192156008807199 |
| **Glycosphingolipid metabolism** | 0.291543746312 | 0.00607553428773 | Galactose | C01582 | 203.05373260968 | 74.2978463704007 | 0.707545 | 0.003846 | 0.888464 | 0.0029012672876610 5 | 0.192156008807199 |
| **Glycosphingolipid metabolism** | 0.291543746312 | 0.00607553428773 | beta-D-Galactose | C00962 | 203.05373260968 | 74.2978463704007 | 0.707545 | 0.003846 | 0.888464 | 0.00290126728766105 | 0.192156008807199 |
| **Tyrosine metabolism** | 0.436905874443 | 0.00917404265803 | adrenochrome o-semiquinone | CE5541 | 203.05373260968 | 74.2978463704007 | 0.707545 | 0.003846 | 0.888464 | 0.00290126728766105 | 0.192156008807199 |
| **Caffeine metabolism** | 0.226313297646 | 0.0134036375154 | 1,7-Dimethylxanthine; Paraxanthine | C13747 | 203.05373260968 | 74.2978463704007 | 0.707545 | 0.003846 | 0.888464 | 0.00290126728766105 | 0.192156008807199 |
| **Caffeine metabolism** | 0.226313297646 | 0.0134036375154 | | C07480 | 203.05373260968 | 74.2978463704007 | 0.707545 | 0.003846 | 0.888464 | 0.00290126728766105 | 0.192156008807199 |
| **Glycosphingolipid biosynthesis - ganglioseries** | 0.378251327416 | 0.0325194161802 | D-Galactose | C00124 | 203.05373260968 | 74.2978463704007 | 0.707545 | 0.003846 | 0.888464 | 0.00290126728766105 | 0.192156008807199 |
| **Glycosphingolipid biosynthesis - ganglioseries** | 0.378251327416 | 0.0325194161802 | Galactose | C01582 | 203.05373260968 | 74.2978463704007 | 0.707545 | 0.003846 | 0.888464 | 0.00290126728766105 | 0.192156008807199 |
| **Phosphatidylinositol phosphate metabolism** | 0.673947532305 | 0.0821391815902 | myo-Inositol; D-myo-Inositol; 1D-myo-Inositol; L-myo-Inositol; 1L-myo-Inositol; meso-Inositol; Inositol; Dambose; Cyclohexitol; Meat sugar; Bios I | C00137 | 203.05373260968 | 74.2978463704007 | 0.707545 | 0.003846 | 0.888464 | 0.00290126728766105 | 0.192156008807199 |
| **Phosphatidylinositol phosphate metabolism** | 0.673947532305 | 0.0821391815902 | D-Galactose | C00124 | 203.05373260968 | 74.2978463704007 | 0.707545 | 0.003846 | 0.888464 | 0.00290126728766105 | 0.192156008807199 |

| | | | | | | | | | | | |
|---|---|---|---|---|---|---|---|---|---|---|---|
| **Glycolysis and Gluconeogenesis** | 0.76425000843 3 | 0.106764598315 | beta-D-Glucose | C00221 | 203.05373260968 | 74.2978463704007 | 0.707545 | 0.003846 | 0.888464 | 0.00290126728766105 | 0.192156008807199 |
| **Glycolysis and Gluconeogenesis** | 0.76425000843 3 | 0.106764598315 | D-Glucose; Grape sugar; Dextrose | C00031 | 203.05373260968 | 74.2978463704007 | 0.707545 | 0.003846 | 0.888464 | 0.00290126728766105 | 0.192156008807199 |
| **Glycolysis and Gluconeogenesis** | 0.76425000843 3 | 0.106764598315 | alpha-D-Glucose | C00267 | 203.05373260968 | 74.2978463704007 | 0.707545 | 0.003846 | 0.888464 | 0.00290126728766105 | 0.192156008807199 |
| **Aspartate and asparagine metabolism** | 0.82287336873 2 | 0.123478035657 | O-Acetylcarnitine; O-Acetyl-L-carnitine | C02571 | 204.122297531216 | 91.1524890948529 | 0.751563 | 0.004096 | 0.907027 | 0.00290826579096826 | 0.192156008807199 |
| **Aspartate and asparagine metabolism** | 0.82287336873 2 | 0.123478035657 | N1,N12-Diacetylspermine | C03413 | 309.226965042255 | 20.824989804138 | 0.880258 | 0.004752 | 0.957284 | 0.00288243445805536 | 0.192156008807199 |
| **Vitamin B3 (nicotinate and nicotinamide) metabolism** | 0.69820367706 | 0.130985025415 | Iminoaspartate | C05840 | 133.032939768675 | 481.767486679155 | 0.919843 | 0.004649 | 0.971473 | 0.00271111469010 91 | 0.192156008807199 |
| **Galactose metabolism** | 0.82143718677 2 | 0.166638916907 | D-Mannose; Mannose; Seminose; Carubinose | C00159 | 203.05373260968 | 74.2978463704007 | 0.707545 | 0.003846 | 0.888464 | 0.00290126728766105 | 0.192156008807199 |
| **Galactose metabolism** | 0.82143718677 2 | 0.166638916907 | beta-D-Fructose; beta-Fruit sugar; beta-D-arabino-Hexulose; beta-Levulose; Fructose | C02336 | 203.05373260968 | 74.2978463704007 | 0.707545 | 0.003846 | 0.888464 | 0.00290126728766105 | 0.192156008807199 |
| **Galactose metabolism** | 0.82143718677 2 | 0.166638916907 | D-Galactose | C00124 | 203.05373260968 | 74.2978463704007 | 0.707545 | 0.003846 | 0.888464 | 0.00290126728766105 | 0.192156008807199 |
| **Galactose metabolism** | 0.82143718677 2 | 0.166638916907 | beta-D-Glucose | C00221 | 203.05373260968 | 74.2978463704007 | 0.707545 | 0.003846 | 0.888464 | 0.00290126728766105 | 0.192156008807199 |
| **Galactose metabolism** | 0.82143718677 2 | 0.166638916907 | D-Glucose; Grape sugar; Dextrose | C00031 | 203.05373260968 | 74.2978463704007 | 0.707545 | 0.003846 | 0.888464 | 0.002901267 28 | 0.192156008807199 |

| | | | | | | | | | | | |
|---|---|---|---|---|---|---|---|---|---|---|---|
| | | | | | | | | | | 766105 | |
| **Galactose metabolism** | 0.8214371867772 | 0.166638916907 | alpha-D-Glucose | C00267 | 203.05373260968 | 74.2978463704007 | 0.707545 | 0.003846 | 0.888464 | 0.00290126728766105 | 0.192156008807199 |
| **Galactose metabolism** | 0.8214371867772 | 0.166638916907 | D-Tagatose; lyxo-Hexulose | C00795 | 203.05373260968 | 74.2978463704007 | 0.707545 | 0.003846 | 0.888464 | 0.00290126728766105 | 0.192156008807199 |
| **Galactose metabolism** | 0.8214371867772 | 0.166638916907 | D-Fructose; Levulose; Fruit sugar; D-arabino-Hexulose | C00095 | 203.05373260968 | 74.2978463704007 | 0.707545 | 0.003846 | 0.888464 | 0.00290126728766105 | 0.192156008807199 |
| **Fructose and mannose metabolism** | 0.854540537723 | 0.246018290225 | D-Mannose; Mannose; Seminose; Carubinose | C00159 | 203.05373260968 | 74.2978463704007 | 0.707545 | 0.003846 | 0.888464 | 0.00290126728766105 | 0.192156008807199 |
| **Fructose and mannose metabolism** | 0.854540537723 | 0.246018290225 | alpha-D-Glucose | C00267 | 203.05373260968 | 74.2978463704007 | 0.707545 | 0.003846 | 0.888464 | 0.00290126728766105 | 0.192156008807199 |
| **Fructose and mannose metabolism** | 0.854540537723 | 0.246018290225 | D-Fructose; Levulose; Fruit sugar; D-arabino-Hexulose | C00095 | 203.05373260968 | 74.2978463704007 | 0.707545 | 0.003846 | 0.888464 | 0.00290126728766105 | 0.192156008807199 |
| **Fructose and mannose metabolism** | 0.854540537723 | 0.246018290225 | D-Glucose; Grape sugar; Dextrose | C00031 | 203.05373260968 | 74.2978463704007 | 0.707545 | 0.003846 | 0.888464 | 0.00290126728766105 | 0.192156008807199 |
| **Sialic acid metabolism** | 0.883202469758 | 0.279395532525 | D-Galactose | C00124 | 203.05373260968 | 74.2978463704007 | 0.707545 | 0.003846 | 0.888464 | 0.00290126728766105 | 0.192156008807199 |
| **Sialic acid metabolism** | 0.883202469758 | 0.279395532525 | beta-D-Glucose | C00221 | 203.05373260968 | 74.2978463704007 | 0.707545 | 0.003846 | 0.888464 | 0.00290126728766105 | 0.192156008807199 |
| **Sialic acid metabolism** | 0.883202469758 | 0.279395532525 | myo-Inositol; D-myo-Inositol; 1D-myo-Inositol; L-myo-Inositol; 1L-myo-Inositol; meso-Inositol; Inositol; Dambose; Cyclohexitol; Meat sugar; Bios I | C00137 | 203.05373260968 | 74.2978463704007 | 0.707545 | 0.003846 | 0.888464 | 0.00290126728766105 | 0.192156008807199 |

| | | | | | | | | | | | |
|---|---|---|---|---|---|---|---|---|---|---|---|
| **Sialic acid metabolism** | 0.883202469758 | 0.279395532525 | alpha-D-Glucose | C00267 | 203.05373260968 | 74.2978463704007 | 0.707545 | 0.003846 | 0.888464 | 0.002901267287661 05 | 0.192156008807199 |
| **Sialic acid metabolism** | 0.883202469758 | 0.279395532525 | Galactose | C01582 | 203.05373260968 | 74.2978463704007 | 0.707545 | 0.003846 | 0.888464 | 0.002901267287661 05 | 0.192156008807199 |
| **Pyrimidine metabolism** | 0.92475584473 | 0.292941294268 | Deoxyribose; 2-Deoxy-D-erythro-pentose; Thyminose; 2-Deoxy-D-ribose | C01801 | 203.05373260968 | 74.2978463704007 | 0.707545 | 0.003846 | 0.888464 | 0.002901267287661 05 | 0.192156008807199 |
| **Pentose phosphate pathway** | 0.933609039998 | 0.359900575187 | Deoxyribose; 2-Deoxy-D-erythro-pentose; Thyminose; 2-Deoxy-D-ribose | C01801 | 203.05373260968 | 74.2978463704007 | 0.707545 | 0.003846 | 0.888464 | 0.002901267287661 05 | 0.192156008807199 |
| **Pentose phosphate pathway** | 0.933609039998 | 0.359900575187 | beta-D-Glucose | C00221 | 203.05373260968 | 74.2978463704007 | 0.707545 | 0.003846 | 0.888464 | 0.002901267287661 05 | 0.192156008807199 |
| **Tryptophan metabolism** | 0.991312719246 | 0.543477262998 | | C00936 | 203.05373260968 | 74.2978463704007 | 0.707545 | 0.003846 | 0.888464 | 0.002901267287661 05 | 0.192156008807199 |
| **Starch and Sucrose Metabolism** | 0.876115428409 | 0.999986263096 | alpha-D-Glucose | C00267 | 203.05373260968 | 74.2978463704007 | 0.707545 | 0.003846 | 0.888464 | 0.002901267287661 05 | 0.192156008807199 |
| **Starch and Sucrose Metabolism** | 0.876115428409 | 0.999986263096 | beta-D-Glucose | C00221 | 203.05373260968 | 74.2978463704007 | 0.707545 | 0.003846 | 0.888464 | 0.002901267287661 05 | 0.192156008807199 |
| **Starch and Sucrose Metabolism** | 0.876115428409 | 0.999986263096 | D-Glucose; Grape sugar; Dextrose | C00031 | 203.05373260968 | 74.2978463704007 | 0.707545 | 0.003846 | 0.888464 | 0.002901267287661 05 | 0.192156008807199 |
| **N-Glycan Degradation** | 0.500317023052 | 0.999986263096 | D-Mannose; Mannose; Seminose; Carubinose | C00159 | 203.05373260968 | 74.2978463704007 | 0.707545 | 0.003846 | 0.888464 | 0.002901267287661 05 | 0.192156008807199 |
| **N-Glycan Degradation** | 0.500317023052 | 0.999986263096 | D-Galactose | C00124 | 203.05373260968 | 74.2978463704007 | 0.707545 | 0.003846 | 0.888464 | 0.00290126728 | 0.192156008807199 |

| | | | | | | | | | | | |
|---|---|---|---|---|---|---|---|---|---|---|---|
| | | | | | | | | | | 766105 | |
| **Alanine and Aspartate Metabolism** | 0.929271964075 | 0.999986263096 | | C02362 | 133.032939768675 | 481.767486679155 | 0.919843 | 0.004649 | 0.971473 | 0.0027111146901091 | 0.192156008807199 |
| **Heparan sulfate degradation** | 0.670828513167 | 0.999986263096 | D-Galactose | C00124 | 203.05373260968 | 74.2978463704007 | 0.707545 | 0.003846 | 0.888464 | 0.00290126728766105 | 0.192156008807199 |
| **Propanoate metabolism** | 0.946589142844 | 0.999986263096 | Propinol adenylate; Propionyladenylate | C05983 | 203.05373260968 | 74.2978463704007 | 0.707545 | 0.003846 | 0.888464 | 0.00290126728766105 | 0.192156008807199 |
| **Glycosphingolipid biosynthesis - globoseries** | 0.500317023052 | 0.999986263096 | alpha-D-Galactose | C00984 | 203.05373260968 | 74.2978463704007 | 0.707545 | 0.003846 | 0.888464 | 0.00290126728766105 | 0.192156008807199 |
| **Keratan sulfate degradation** | 0.565178203425 | 0.999986263096 | D-Galactose | C00124 | 203.05373260968 | 74.2978463704007 | 0.707545 | 0.003846 | 0.888464 | 0.00290126728766105 | 0.192156008807199 |
| **Chondroitin sulfate degradation** | 0.565178203425 | 0.999986263096 | D-Galactose | C00124 | 203.05373260968 | 74.2978463704007 | 0.707545 | 0.003846 | 0.888464 | 0.00290126728766105 | 0.192156008807199 |
| **N-Glycan biosynthesis** | 0.783338757398 | 0.999986263096 | D-Mannose; Mannose; Seminose; Carubinose | C00159 | 203.05373260968 | 74.2978463704007 | 0.707545 | 0.003846 | 0.888464 | 0.00290126728766105 | 0.192156008807199 |
| **N-Glycan biosynthesis** | 0.783338757398 | 0.999986263096 | D-Glucose; Grape sugar; Dextrose | C00031 | 203.05373260968 | 74.2978463704007 | 0.707545 | 0.003846 | 0.888464 | 0.00290126728766105 | 0.192156008807199 |
| **Fatty Acid Metabolism** | 0.00136264372743 | 0.000339697971178 | Dodecanoic acid; Dodecanoate; Dodecylcarboxylate; Lauric acid | C02679 | 269.174558814739 | 17.3993398078578 | 0.962523 | 0.017933 | 0.990613 | 0.00925132273475404 | 0.196468734356961 |
| **Glycerophospholipid metabolism** | 5.97538527321e-05 | 0.000329266567154 | Sterol | C00370 | 250.224262076074 | 488.878549975896 | 0.925344 | 3.41E-05 | 0.959743 | 3.09361595268731e-05 | 0.0221922750091705 |
| **Glycerophospholipid metabolism** | 5.9753852 | 0.00032926 | N-Trimethyl-2-aminoethylphosphonate; 2-Trimethylaminoethylphosphonate | C06459 | 169.086005572512 | 464.273075221709 | 0.705646 | 0.002568 | 0.880268 | 0.0019985 | 0.16867003171186 |

| | | | | | | | | | | | |
|---|---|---|---|---|---|---|---|---|---|---|---|
| | 7321 e-05 | 6567154 | | | | | | | | 7948558313 | |
| **Glycerophospholipid metabolism** | 5.9753852 7321 e-05 | 0.00032926 6567154 | myo-Inositol; D-myo-Inositol; 1D-myo-Inositol; L-myo-Inositol; 1L-myo-Inositol; meso-Inositol; Inositol; Dambose; Cyclohexitol; Meat sugar; Bios I | C00137 | 203.05373260968 | 74.2978463704007 | 0.707545 | 0.003846 | 0.888464 | 0.0029012672876105 | 0.192156008807199 |
| **Glycerophospholipid metabolism** | 5.9753852 7321 e-05 | 0.00032926 6567154 | (R)-glycerol 1-acetate | CE0520 | 203.05373260968 | 74.2978463704007 | 0.707545 | 0.003846 | 0.888464 | 0.0029012672876105 | 0.192156008807199 |
| **Glycerophospholipid metabolism** | 5.9753852 7321 e-05 | 0.00032926 6567154 | Galactose | C01582 | 203.05373260968 | 74.2978463704007 | 0.707545 | 0.003846 | 0.888464 | 0.0029012672876105 | 0.192156008807199 |
| **Glycerophospholipid metabolism** | 5.9753852 7321 e-05 | 0.00032926 6567154 | Linoleate; Linoleic acid; (9Z,12Z)-Octadecadienoic acid; 9-cis,12-cis-Octadecadienoate; 9-cis,12-cis-Octadecadienoic acid | C01595 | 281.24752253069 | 33.784067698132 | 0.742706 | 0.006638 | 0.912449 | 0.0046191669710989 9 | 0.19290463114884 |
| **Glycerophospholipid metabolism** | 5.9753852 7321 e-05 | 0.00032926 6567154 | S-Adenosyl-L-homocysteine; S-Adenosylhomocysteine | C00021 | 407.10724469524 | 86.7848403342757 | 1.089523 | 0.007888 | 1.154639 | 0.0037914134359235 | 0.19290463114884 |
| **Glycerophospholipid metabolism** | 5.9753852 7321 e-05 | 0.00032926 6567154 | (5Z,8Z,11Z,14Z)-Icosatetraenoic acid; Arachidonate; Arachidonic acid; cis-5,8,11,14-Eicosatetraenoic acid | C00219 | 307.246565208586 | 518.104796795187 | 0.972521 | 0.007512 | 0.991165 | 0.0040281506735649 9 | 0.19290463114884 |
| **Carnitine shuttle** | 0.0831103651174 | 0.00087481745998 4 | arachidyl carnitine | arachcrn | 540.363210427572 | 523.451786821455 | 0.759818 | 0.00473 | 0.912563 | 0.0032924926422423 | 0.19290463114884 |
| **De novo fatty acid biosynthesis** | 0.104709729668 | 0.00101148958738 | Linoleate; Linoleic acid; (9Z,12Z)-Octadecadienoic acid; 9-cis,12-cis-Octadecadienoate; 9-cis,12-cis-Octadecadienoic acid | C01595 | 281.24752253069 | 33.784067698132 | 0.742706 | 0.006638 | 0.912449 | 0.0046191669710989 9 | 0.19290463114884 |
| **De novo fatty acid biosynthesis** | 0.104709729668 | 0.00101148958738 | (4Z,7Z,10Z,13Z,16Z,19Z)-Docosahexaenoic acid; 4,7,10,13,16,19-Docosahexaenoic acid; Docosahexaenoic acid | C06429 | 328.240480663843 | 539.109765179623 | 1.032436 | 0.005957 | 1.054484 | 0.0030671930822523 4 | 0.19290463114884 |
| **De novo fatty acid biosynthesis** | 0.104709729668 | 0.00101148958738 | (5Z,8Z,11Z,14Z)-Icosatetraenoic acid; Arachidonate; Arachidonic acid; cis-5,8,11,14-Eicosatetraenoic acid | C00219 | 307.246565208586 | 518.104796795187 | 0.972521 | 0.007512 | 0.991165 | 0.0040281506735649 9 | 0.19290463114884 |

| | | | | | | | | | | | |
|---|---|---|---|---|---|---|---|---|---|---|---|
| **Lysine metabolism** | 0.12545355247 | 0.00148162 64063 | Carnitine; gamma-Trimethyl-hydroxybutyrobetaine; 3-Hydroxy-4-trimethylammoniobutanoate | C00487 | 162.11254587791 | 93.3036471083725 | 0.760748 | 0.006384 | 0.91807 | 0.00435358070530589 | 0.19290463114884 |
| **Lysine metabolism** | 0.12545355247 | 0.00148162 64063 | N6,N6,N6-Trimethyl-L-lysine | C03793 | 257.146796364654 | 121.291800902871 | 0.845864 | 0.006521 | 0.948062 | 0.00402096306555932 | 0.19290463114884 |
| **Lysine metabolism** | 0.12545355247 | 0.00148162 64063 | S-Adenosyl-L-homocysteine; S-Adenosylhomocysteine | C00021 | 407.10724469524 | 86.7848403342757 | 1.089523 | 0.007888 | 1.154639 | 0.00379141343359235 | 0.19290463114884 |
| **Urea cycle/amino group metabolism** | 0.22170906292 6 | 0.00252163988827 | S-Adenosyl-L-homocysteine; S-Adenosylhomocysteine | C00021 | 407.10724469524 | 86.7848403342757 | 1.089523 | 0.007888 | 1.154639 | 0.00379141343359235 | 0.19290463114884 |
| **Urea cycle/amino group metabolism** | 0.22170906292 6 | 0.00252163988827 | Phenylethanolamine; 2-Amino-1-phenylethanol | C02735 | 138.091443322759 | 32.6063494904282 | 0.842558 | 0.006983 | 0.947711 | 0.00430319848558158 | 0.19290463114884 |
| **Drug metabolism - cytochrome P450** | 0.33976603682 5 | 0.00657261942317 | 2-Hydroxyfelbamate | C16582 | 323.082070585836 | 119.575736641393 | 1.094165 | 0.007329 | 1.16235 | 0.00352544984850578 | 0.19290463114884 |
| **Drug metabolism - cytochrome P450** | 0.33976603682 5 | 0.00657261942317 | p-Hydroxyfelbamate | C16584 | 323.082070585836 | 119.575736641393 | 1.094165 | 0.007329 | 1.16235 | 0.00352544984850578 | 0.19290463114884 |
| **Tyrosine metabolism** | 0.43690587444 3 | 0.00917404265803 | L-Phenylalanine; (S)-alpha-Amino-beta-phenylpropionic acid | C00079 | 166.086229234692 | 119.872320624957 | 0.797537 | 0.007121 | 0.932674 | 0.00461475470142025 | 0.19290463114884 |
| **Tyrosine metabolism** | 0.43690587444 3 | 0.00917404265803 | S-Adenosyl-L-homocysteine; S-Adenosylhomocysteine | C00021 | 407.10724469524 | 86.7848403342757 | 1.089523 | 0.007888 | 1.154639 | 0.00379141343359235 | 0.19290463114884 |
| **Tyrosine metabolism** | 0.43690587444 3 | 0.00917404265803 | Tyramine; 2-(p-Hydroxyphenyl)ethylamine | C00483 | 138.091443322759 | 32.6063494904282 | 0.842558 | 0.006983 | 0.947711 | 0.00430319848558158 | 0.19290463114884 |
| **Tyrosine metabolism** | 0.43690587444 3 | 0.00917404265803 | | CE2172 | 166.086229234692 | 119.872320624957 | 0.797537 | 0.007121 | 0.932674 | 0.00461475470 | 0.19290463114884 |

| | | | | | | | | | | | |
|---|---|---|---|---|---|---|---|---|---|---|---|
| | | | | | | | | | | 1420 25 | |
| **Caffeine metabolism** | 0.226 3132 9764 6 | 0.013 40363 75154 | 5-Acetylamino-6-formylamino-3-methyluracil; AFMU | C16365 | 249.0606 72142736 | 99.3598241199535 | 0.76693 8 | 0.006 875 | 0.921 569 | 0.004 6316 4014 4932 34 | 0.19290463114884 |
| **Ubiquinone Biosynthesis** | 0.277 4459 0961 2 | 0.018 59815 69187 | S-Adenosyl-L-homocysteine; S-Adenosylhomocysteine | C00021 | 407.1072 4469524 | 86.7848403342757 | 1.08952 3 | 0.007 888 | 1.154 639 | 0.003 7914 1343 3592 35 | 0.19290463114884 |
| **Glycine, serine, alanine and threonine metabolism** | 0.518 5832 9863 2 | 0.019 90447 96148 | S-Adenosyl-L-homocysteine; S-Adenosylhomocysteine | C00021 | 407.1072 4469524 | 86.7848403342757 | 1.08952 3 | 0.007 888 | 1.154 639 | 0.003 7914 1343 3592 35 | 0.19290463114884 |
| **Fatty acid activation** | 0.481 1620 6370 4 | 0.019 93238 27058 | Linoleate; Linoleic acid; (9Z,12Z)-Octadecadienoic acid; 9-cis,12-cis-Octadecadienoate; 9-cis,12-cis-Octadecadienoic acid | C01595 | 281.2475 2253069 | 33.784067698132 | 0.74270 6 | 0.006 638 | 0.912 449 | 0.004 6191 6697 1098 99 | 0.19290463114884 |
| **Fatty acid activation** | 0.481 1620 6370 4 | 0.019 93238 27058 | vaccenic acid | vacc | 281.2475 2253069 | 33.784067698132 | 0.74270 6 | 0.006 638 | 0.912 449 | 0.004 6191 6697 1098 99 | 0.19290463114884 |
| **Fatty acid activation** | 0.481 1620 6370 4 | 0.019 93238 27058 | octadecenoate (n-C18:1) | ocdcea | 281.2475 2253069 | 33.784067698132 | 0.74270 6 | 0.006 638 | 0.912 449 | 0.004 6191 6697 1098 99 | 0.19290463114884 |
| **Vitamin E metabolism** | 0.507 3545 8165 | 0.023 18729 8019 | | CE7047 | 438.3433 23184971 | 41.1997352870357 | 0.95650 3 | 0.005 666 | 0.985 153 | 0.003 1429 9625 3437 72 | 0.19290463114884 |
| **Di-unsaturated fatty acid beta-oxidation** | 0.378 2513 2741 6 | 0.032 51941 61802 | Linoleate; Linoleic acid; (9Z,12Z)-Octadecadienoic acid; 9-cis,12-cis-Octadecadienoate; 9-cis,12-cis-Octadecadienoic acid | C01595 | 281.2475 2253069 | 33.784067698132 | 0.74270 6 | 0.006 638 | 0.912 449 | 0.004 6191 6697 1098 99 | 0.19290463114884 |
| **Biopterin metabolism** | 0.516 7582 9544 2 | 0.062 27580 94373 | L-Phenylalanine; (S)-alpha-Amino-beta-phenylpropionic acid | C00079 | 166.0862 29234692 | 119.872320624957 | 0.79753 7 | 0.007 121 | 0.932 674 | 0.004 6147 5470 1420 25 | 0.19290463114884 |
| **Porphyrin metabolism** | 0.745 8369 1149 5 | 0.115 58802 7425 | Fe3+; Fe(III); Ferric ion; Iron(3+) | C14819 | 140.9032 51581497 | 77.195477277448 | 0.68453 8 | 0.005 763 | 0.888 348 | 0.004 3667 9741 9495 64 | 0.19290463114884 |

| | | | | | | | | | | | |
|---|---|---|---|---|---|---|---|---|---|---|---|
| **Porphyrin metabolism** | 0.74583691149 5 | 0.115588027425 | Fe2+; Fe(II); Ferrous ion; Iron(2+) | C14818 | 140.903251581497 | 77.195477277448 | 0.684538 | 0.005763 | 0.888348 | 0.004366797419495 64 | 0.19290463114884 |
| **Porphyrin metabolism** | 0.74583691149 5 | 0.115588027425 | Iron | C00023 | 140.903251581497 | 77.195477277448 | 0.684538 | 0.005763 | 0.888348 | 0.004366797419495 64 | 0.19290463114884 |
| **Aspartate and asparagine metabolism** | 0.82287336873 2 | 0.123478035657 | Carnitine; gamma-Trimethyl-hydroxybutyrobetaine; 3-Hydroxy-4-trimethylammoniobutanoate | C00487 | 162.11254587791 | 93.3036471083725 | 0.760748 | 0.006384 | 0.91807 | 0.004353580705305 89 | 0.19290463114884 |
| **Vitamin B3 (nicotinate and nicotinamide) metabolism** | 0.69820367706 | 0.130985025415 | S-Adenosyl-L-homocysteine; S-Adenosylhomocysteine | C00021 | 407.10724469524 | 86.7848403342757 | 1.089523 | 0.007888 | 1.154639 | 0.003791413433592 35 | 0.19290463114884 |
| **Histidine metabolism** | 0.69820367706 | 0.130985025415 | S-Adenosyl-L-homocysteine; S-Adenosylhomocysteine | C00021 | 407.10724469524 | 86.7848403342757 | 1.089523 | 0.007888 | 1.154639 | 0.003791413433592 35 | 0.19290463114884 |
| **Omega-6 fatty acid metabolism** | 0.77744172845 1 | 0.179119678411 | (5Z,8Z,11Z,14Z)-Icosatetraenoic acid; Arachidonate; Arachidonic acid; cis-5,8,11,14-Eicosatetraenoic acid | C00219 | 307.246565208586 | 518.104796795187 | 0.972521 | 0.007512 | 0.991165 | 0.004028150673564 99 | 0.19290463114884 |
| **Methionine and cysteine metabolism** | 0.87740544892 2 | 0.198255230658 | S-Adenosyl-L-homocysteine; S-Adenosylhomocysteine | C00021 | 407.10724469524 | 86.7848403342757 | 1.089523 | 0.007888 | 1.154639 | 0.003791413433592 35 | 0.19290463114884 |
| **Leukotriene metabolism** | 0.88759672329 5 | 0.210733231746 | 10,11-dihydro-12-epi-leukotriene B4 | CE4988 | 339.251168308686 | 554.421674845691 | 0.740259 | 0.006852 | 0.912231 | 0.004773668633582 31 | 0.19290463114884 |
| **Leukotriene metabolism** | 0.88759672329 5 | 0.210733231746 | 6,7-dihydro-12-epi-LTB4 | CE5350 | 339.251168308686 | 554.421674845691 | 0.740259 | 0.006852 | 0.912231 | 0.004773668633582 31 | 0.19290463114884 |
| **Leukotriene metabolism** | 0.88759672329 5 | 0.210733231746 | 6,7-dihydro-leukotriene B4 | CE5352 | 339.251168308686 | 554.421674845691 | 0.740259 | 0.006852 | 0.912231 | 0.004773668633582 31 | 0.19290463114884 |
| **Leukotriene metabolism** | 0.88759672329 5 | 0.210733231746 | (5Z,8Z,11Z,14Z)-Icosatetraenoic acid; Arachidonate; Arachidonic acid; cis-5,8,11,14-Eicosatetraenoic acid | C00219 | 307.246565208586 | 518.104796795187 | 0.972521 | 0.007512 | 0.991165 | 0.004028150673564 99 | 0.19290463114884 |

| | | | | | | | | | | | |
|---|---|---|---|---|---|---|---|---|---|---|---|
| | | | | | | | | | | 3564 99 | |
| **Leukotriene metabolism** | 0.887 5967 2329 5 | 0.210 73323 1746 | 10,11-dihydro-leukotriene B4 | CE4987 | 339.2511 68308686 | 554.421674845691 | 0.74025 9 | 0.006 852 | 0.912 231 | 0.004 7736 6863 3582 31 | 0.19290463114884 |
| **Leukotriene metabolism** | 0.887 5967 2329 5 | 0.210 73323 1746 | 20-dihydroxyleukotriene B4 | CE2056 | 185.1146 41021963 | 410.252213478771 | 0.72385 2 | 0.005 594 | 0.902 205 | 0.004 0299 5991 1236 | 0.19290463114884 |
| **Arachidonic acid metabolism** | 0.921 3442 8956 1 | 0.261 57551 043 | 11,12-DHET; (5Z,8Z,14Z)-11,12-Dihydroxyeicosa-5,8,14-trienoic acid; (5Z,8Z,14Z)-11,12-Dihydroxyicosa-5,8,14-trienoic acid | C14774 | 339.2511 68308686 | 554.421674845691 | 0.74025 9 | 0.006 852 | 0.912 231 | 0.004 7736 6863 3582 31 | 0.19290463114884 |
| **Arachidonic acid metabolism** | 0.921 3442 8956 1 | 0.261 57551 043 | 14,15-DHET; (5Z,8Z,11Z)-14,15-Dihydroxyeicosa-5,8,11-trienoic acid; (5Z,8Z,11Z)-14,15-Dihydroxyicosa-5,8,11-trienoic acid | C14775 | 339.2511 68308686 | 554.421674845691 | 0.74025 9 | 0.006 852 | 0.912 231 | 0.004 7736 6863 3582 31 | 0.19290463114884 |
| **Arachidonic acid metabolism** | 0.921 3442 8956 1 | 0.261 57551 043 | 5,6-DHET; (8Z,11Z,14Z)-5,6-Dihydroxyeicosa-8,11,14-trienoic acid; (8Z,11Z,14Z)-5,6-Dihydroxyicosa-8,11,14-trienoic acid | C14772 | 339.2511 68308686 | 554.421674845691 | 0.74025 9 | 0.006 852 | 0.912 231 | 0.004 7736 6863 3582 31 | 0.19290463114884 |
| **Arachidonic acid metabolism** | 0.921 3442 8956 1 | 0.261 57551 043 | 8,9-DHET; (5Z,11Z,14Z)-8,9-Dihydroxyeicosa-5,11,14-trienoic acid; (5Z,11Z,14Z)-8,9-Dihydroxyicosa-5,11,14-trienoic acid | C14773 | 339.2511 68308686 | 554.421674845691 | 0.74025 9 | 0.006 852 | 0.912 231 | 0.004 7736 6863 3582 31 | 0.19290463114884 |
| **Arachidonic acid metabolism** | 0.921 3442 8956 1 | 0.261 57551 043 | (5Z,8Z,11Z,14Z)-Icosatetraenoic acid; Arachidonate; Arachidonic acid; cis-5,8,11,14-Eicosatetraenoic acid | C00219 | 307.2465 65208586 | 518.104796795187 | 0.97252 1 | 0.007 512 | 0.991 165 | 0.004 0281 5067 3564 99 | 0.19290463114884 |
| **Prostaglandin formation from arachidonate** | 0.973 5635 0286 7 | 0.427 66573 2887 | | CE5930 | 185.1146 41021963 | 410.252213478771 | 0.72385 2 | 0.005 594 | 0.902 205 | 0.004 0299 5991 1236 | 0.19290463114884 |
| **Prostaglandin formation from arachidonate** | 0.973 5635 0286 7 | 0.427 66573 2887 | 11-dehydrothromboxane B2 | CE1447 | 185.1146 41021963 | 410.252213478771 | 0.72385 2 | 0.005 594 | 0.902 205 | 0.004 0299 5991 1236 | 0.19290463114884 |
| **Prostaglandin formation from arachidonate** | 0.973 5635 0286 7 | 0.427 66573 2887 | | CE5534 | 185.1146 41021963 | 410.252213478771 | 0.72385 2 | 0.005 594 | 0.902 205 | 0.004 0299 5991 1236 | 0.19290463114884 |
| **Prostaglandin formation from arachidonate** | 0.973 5635 0286 7 | 0.427 66573 2887 | | CE5926 | 185.1146 41021963 | 410.252213478771 | 0.72385 2 | 0.005 594 | 0.902 205 | 0.004 0299 5991 1236 | 0.19290463114884 |

| | | | | | | | | | | | |
|---|---|---|---|---|---|---|---|---|---|---|---|
| **Prostaglandin formation from arachidonate** | 0.97356350286 7 | 0.427665732887 | | CE5533 | 185.114641021963 | 410.252213478771 | 0.723852 | 0.005594 | 0.902205 | 0.0040299599112 36 | 0.19290463114884 |
| **Prostaglandin formation from arachidonate** | 0.97356350286 7 | 0.427665732887 | (5Z,8Z,11Z,14Z)-Icosatetraenoic acid; Arachidonate; Arachidonic acid; cis-5,8,11,14-Eicosatetraenoic acid | C00219 | 307.246565208586 | 518.104796795187 | 0.972521 | 0.007512 | 0.991165 | 0.0040281506735 6499 | 0.19290463114884 |
| **Androgen and estrogen biosynthesis and metabolism** | 0.98779638849 3 | 0.510935384898 | S-Adenosyl-L-homocysteine; S-Adenosylhomocysteine | C00021 | 407.10724469524 | 86.7848403342757 | 1.089523 | 0.007888 | 1.154639 | 0.0037914134335 9235 | 0.19290463114884 |
| **Tryptophan metabolism** | 0.99131271924 6 | 0.543477262998 | S-Adenosyl-L-homocysteine; S-Adenosylhomocysteine | C00021 | 407.10724469524 | 86.7848403342757 | 1.089523 | 0.007888 | 1.154639 | 0.0037914134335 9235 | 0.19290463114884 |
| **Prostaglandin formation from dihomo gama-linoleic acid** | 0.42583461114 9 | 0.999986263096 | 15-hydroperoxyeicosa-8Z,11Z,13E-trienoate | CE6230 | 339.251168308686 | 554.421674845691 | 0.740259 | 0.006852 | 0.912231 | 0.0047736686335 8231 | 0.19290463114884 |
| **Vitamin B12 (cyanocobalamin) metabolism** | 0.24212119680 7 | 0.999986263096 | S-Adenosyl-L-homocysteine; S-Adenosylhomocysteine | C00021 | 407.10724469524 | 86.7848403342757 | 1.089523 | 0.007888 | 1.154639 | 0.0037914134335 9235 | 0.19290463114884 |
| **Bile acid biosynthesis** | 0.99899436648 | 0.999986263096 | 7alpha-Hydroxycholest-4-en-3-one | C05455 | 485.300476040834 | 88.8392543646686 | 0.779597 | 0.007373 | 0.927139 | 0.0048705015094 7976 | 0.19290463114884 |
| **Vitamin D3 (cholecalciferol) metabolism** | 0.67082851316 7 | 0.999986263096 | Calcidiol; 25-Hydroxyvitamin D3; Calcifediol; Calcifediol anhydrous | C01561 | 485.300476040834 | 88.8392543646686 | 0.779597 | 0.007373 | 0.927139 | 0.0048705015094 7976 | 0.19290463114884 |
| **Phytanic acid peroxisomal oxidation** | 0.30549389355 5 | 0.0113522941772 | 2(R)-pristanal | CE5127 | 285.296896404244 | 40.5011635726067 | 0.78386 | 0.007976 | 0.929929 | 0.0052176847724 8663 | 0.194196962906609 |
| **Phytanic acid peroxisomal oxidation** | 0.30549389355 5 | 0.0113522941772 | 2(S)-pristanal | CE5124 | 285.296896404244 | 40.5011635726067 | 0.78386 | 0.007976 | 0.929929 | 0.0052176847724 8663 | 0.194196962906609 |
| **Glycerophospholipid metabolism** | 5.97538527321 e-05 | 0.0003292665671 54 | (9Z,12Z,15Z)-Octadecatrienoic acid; alpha-Linolenic acid; 9,12,15-Octadecatrienoic acid; Linolenate; alpha-Linolenate | C06427 | 279.231911695023 | 28.1891724631974 | 0.739227 | 0.59435 | 0.919419 | 0.0066300345659 458 | 0.195585714469996 |

| | | | | | | | | | | | |
|---|---|---|---|---|---|---|---|---|---|---|---|
| **Carnitine shuttle** | 0.0831103651174 | 0.000874817459984 | L-Carnitine; L-gamma-Trimethyl-beta-hydroxybutyrobetaine; Vitamin BT; 3-Carboxy-2-hydroxy-N,N,N-trimethyl-1-propanaminium hydroxide, inner salt; Levocarnitine; (R)-Carnitine | C00318 | 163.115801406981 | 88.5776592293596 | 0.732694 | 0.007677 | 0.912029 | 0.00536372015369579 | 0.194468741890133 |
| **Lysine metabolism** | 0.125453555247 | 0.0014816264063 | L-Carnitine; L-gamma-Trimethyl-beta-hydroxybutyrobetaine; Vitamin BT; 3-Carboxy-2-hydroxy-N,N,N-trimethyl-1-propanaminium hydroxide, inner salt; Levocarnitine; (R)-Carnitine | C00318 | 163.115801406981 | 88.5776592293596 | 0.732694 | 0.007677 | 0.912029 | 0.00536372015369579 | 0.194468741890133 |
| **Saturated fatty acids beta-oxidation** | 0.182491247164 | 0.00237820952794 | L-Carnitine; L-gamma-Trimethyl-beta-hydroxybutyrobetaine; Vitamin BT; 3-Carboxy-2-hydroxy-N,N,N-trimethyl-1-propanaminium hydroxide, inner salt; Levocarnitine; (R)-Carnitine | C00318 | 163.115801406981 | 88.5776592293596 | 0.732694 | 0.007677 | 0.912029 | 0.00536372015369579 | 0.194468741890133 |
| **Glycerophospholipid metabolism** | 5.97538527321e-05 | 0.000329266567154 | Choline; Bilineurine | C00114 | 104.107443902613 | 78.7452217926459 | 0.805551 | 0.0102 | 0.941055 | 0.00641288722823619 | 0.195585714469996 |
| **Glycerophospholipid metabolism** | 5.97538527321e-05 | 0.000329266567154 | 2-Deoxy-5-keto-D-gluconic acid 6-phosphate; DKHP | C06893 | 327.007830935988 | 550.437193098651 | 0.758058 | 0.009717 | 0.925383 | 0.00648738452413733 | 0.195585714469996 |
| **Glycerophospholipid metabolism** | 5.97538527321e-05 | 0.000329266567154 | Sphinganine; Dihydrosphingosine; 2-Amino-1,3-dihydroxyoctadecane | C00836 | 370.293118638006 | 94.0202052076728 | 0.853676 | 0.014078 | 0.959976 | 0.00823705347633774 | 0.195585714469996 |
| **Glycerophospholipid metabolism** | 5.97538527321e-05 | 0.000329266567154 | sn-Glycerol 3-phosphate; Glycerophosphoric acid; D-Glycerol 1-phosphate; Glycerol-3-phosphate | C00093 | 256.983790028814 | 124.33968809078 | 1.047063 | 0.01211 | 1.0819 | 0.00588730598760534 | 0.195585714469996 |
| **Glycerophospholipid metabolism** | 5.97538527321e-05 | 0.000329266567154 | CDP-choline; Cytidine 5'-diphosphocholine; Citicoline | C00307 | 557.097268018433 | 110.96015512232 | 0.802193 | 0.015547 | 0.947817 | 0.00960554270191638 | 0.196956027492531 |
| **Glycerophospholipid metabolism** | 5.97538527321e-05 | 0.000329266567154 | Acetylcholine; O-Acetylcholine | C01996 | 146.117576611507 | 38.6922760654623 | 0.77875 | 0.017016 | 0.943782 | 0.0107716983520912 | 0.199226826059028 |
| **Linoleate metabolism** | 0.00310216651967 | 0.000353032634069 | | C08261 | 189.112039227693 | 453.758387539726 | 0.7742 | 0.003228 | 0.912459 | 0.002267257887823 | 0.177891179432863 |
| **Linoleate metabolism** | 0.00310216651967 | 0.000353032634069 | | CE5527 | 295.226406338711 | 29.037835348842 | 0.70858 | 0.003996 | 0.88962 | 0.00300201834681391 | 0.19290463114884 |

| | | | | | | | | | | | |
|---|---|---|---|---|---|---|---|---|---|---|---|
| **Linoleate metabolism** | 0.0031021665196 7 | 0.000353032634069 | Linoleate; Linoleic acid; (9Z,12Z)-Octadecadienoic acid; 9-cis,12-cis-Octadecadienoate; 9-cis,12-cis-Octadecadienoic acid | C01595 | 281.24752253069 | 33.784067698132 | 0.742706 | 0.006638 | 0.912449 | 0.00461916697109899 | 0.19290463114884 |
| **Carnitine shuttle** | 0.083110365117 4 | 0.00087481745998 4 | stearoylcarnitine | stcrn | 512.335111814579 | 579.824989617636 | 0.783174 | 0.013283 | 0.939433 | 0.00846010235033501 | 0.195585714469996 |
| **De novo fatty acid biosynthesis** | 0.104709729668 | 0.001011489587 38 | (9Z,12Z,15Z)-Octadecatrienoic acid; alpha-Linolenic acid; 9,12,15-Octadecatrienoic acid; Linolenate; alpha-Linolenate | C06427 | 279.231911695023 | 28.1891724631974 | 0.739227 | 0.009696 | 0.919419 | 0.00663003456594 58 | 0.195585714469996 |
| **De novo fatty acid biosynthesis** | 0.104709729668 | 0.001011489587 38 | (6Z,9Z,12Z)-Octadecatrienoic acid; 6,9,12-Octadecatrienoic acid; gamma-Linolenic acid; Gamolenic acid | C06426 | 279.231911695023 | 28.1891724631974 | 0.739227 | 0.009696 | 0.919419 | 0.00663003456594 58 | 0.195585714469996 |
| **De novo fatty acid biosynthesis** | 0.104709729668 | 0.001011489587 38 | Tetradecanoic acid; Tetradecanoate; Myristic acid | C06424 | 251.197160068562 | 128.325061638681 | 0.957787 | 0.013649 | 0.988378 | 0.00717150534694222 | 0.195585714469996 |
| **De novo fatty acid biosynthesis** | 0.104709729668 | 0.001011489587 38 | Hexadecanoic acid; Hexadecanoate; Hexadecylic acid; Palmitic acid; Palmitate; Cetylic acid | C00249 | 279.231911695023 | 28.1891724631974 | 0.739227 | 0.009696 | 0.919419 | 0.00663003456594 58 | 0.195585714469996 |
| **De novo fatty acid biosynthesis** | 0.104709729668 | 0.001011489587 38 | Decanoyl-CoA | C05274 | 921.24237107895 | 150.430465242413 | 1.028509 | 0.014277 | 1.049735 | 0.00699517578790609 | 0.195585714469996 |
| **Hexose phosphorylation** | 0.087775159020 4 | 0.00117065111852 | cis-2-Hydroxycinnamate; 2-Coumarinate | C05838 | 165.054695403805 | 116.367243933837 | 0.785605 | 0.012888 | 0.939492 | 0.00819622313896295 | 0.195585714469996 |
| **Saturated fatty acids beta-oxidation** | 0.182491247164 | 0.002378209527 94 | Hexadecanoic acid; Hexadecanoate; Hexadecylic acid; Palmitic acid; Palmitate; Cetylic acid | C00249 | 279.231911695023 | 28.1891724631974 | 0.739227 | 0.009696 | 0.919419 | 0.00663003456594 58 | 0.195585714469996 |
| **Saturated fatty acids beta-oxidation** | 0.182491247164 | 0.002378209527 94 | trans-Dec-2-enoyl-CoA; (2E)-Decenoyl-CoA | C05275 | 921.24237107895 | 150.430465242413 | 1.028509 | 0.014277 | 1.049735 | 0.00699517578790609 | 0.195585714469996 |
| **Saturated fatty acids beta-oxidation** | 0.182491247164 | 0.002378209527 94 | Decanoyl-CoA | C05274 | 921.24237107895 | 150.430465242413 | 1.028509 | 0.014277 | 1.049735 | 0.00699517578 | 0.195585714469996 |

| | | | | | | | | | | | |
|---|---|---|---|---|---|---|---|---|---|---|---|
| | | | | | | | | | | 790609 | |
| **Urea cycle/amino group metabolism** | 0.2217090629226 | 0.00252163988827 | L-Methionine; Methionine; L-2-Amino-4methylthiobutyric acid | C00073 | 172.040105107381 | 89.6909060143427 | 0.76934 | 0.012499 | 0.934207 | 0.00812074625284076 | 0.195585714469996 |
| **Urea cycle/amino group metabolism** | 0.2217090629226 | 0.00252163988827 | Queuine; Base Q | C01449 | 140.068869698569 | 119.161714223817 | 0.824641 | 0.013955 | 0.951868 | 0.00844193863948328 | 0.195585714469996 |
| **Urea cycle/amino group metabolism** | 0.2217090629226 | 0.00252163988827 | 4-Aminobutanal; 4-Aminobutyraldehyde; Butyraldehyde, 4-amino- | C00555 | 89.0795646241381 | 42.0813089452326 | 0.748234 | 0.008425 | 0.919211 | 0.00574002335236268 | 0.195585714469996 |
| **Urea cycle/amino group metabolism** | 0.2217090629226 | 0.00252163988827 | Carbamoyl phosphate | C00169 | 140.982056004229 | 310.601686958314 | 0.759972 | 0.009107 | 0.924639 | 0.00608744857666477 | 0.195585714469996 |
| **Glycosphingolipid metabolism** | 0.291543746312 | 0.00607553428773 | Hexadecanal; Palmitaldehyde | C00517 | 263.236750689057 | 32.3161891554784 | 0.763471 | 0.012306 | 0.932175 | 0.0080603897999465 | 0.195585714469996 |
| **Glycosphingolipid metabolism** | 0.291543746312 | 0.00607553428773 | Hexadecanoic acid; Hexadecanoate; Hexadecylic acid; Palmitic acid; Palmitate; Cetylic acid | C00249 | 279.231911695023 | 28.1891724631974 | 0.739227 | 0.009696 | 0.919419 | 0.0066300345659458 | 0.195585714469996 |
| **Glycosphingolipid metabolism** | 0.291543746312 | 0.00607553428773 | Sphinganine; Dihydrosphingosine; 2-Amino-1,3-dihydroxyoctadecane | C00836 | 370.293118638006 | 94.0202052076728 | 0.853676 | 0.014078 | 0.959976 | 0.00823705347633774 | 0.195585714469996 |
| **Glycosphingolipid metabolism** | 0.291543746312 | 0.00607553428773 | 3'-Phosphoadenylyl sulfate; 3'-Phosphoadenosine 5'-phosphosulfate; 3'-Phospho-5'-adenylyl sulfate; PAPS | C00053 | 507.997040927732 | 146.431216399674 | 1.027868 | 0.015748 | 1.048851 | 0.00768108378459198 | 0.195585714469996 |
| **Drug metabolism - cytochrome P450** | 0.339766036825 | 0.00657261942317 | Citalopram aldehyde | C16612 | 149.059827417723 | 587.065552074052 | 0.733327 | 0.010852 | 0.920298 | 0.00743251628811692 | 0.195585714469996 |
| **Drug metabolism - cytochrome P450** | 0.339766036825 | 0.00657261942317 | Lidocaine | C07073 | 235.180426962792 | 572.59294617902 | 0.756548 | 0.013014 | 0.931499 | 0.00857439446332453 | 0.195585714469996 |
| **Beta-Alanine metabolism** | 0.2310597 | 0.00680777 | 4-Aminobutanal; 4-Aminobutyraldehyde; Butyraldehyde, 4-amino- | C00555 | 89.0795646241381 | 42.0813089452326 | 0.748234 | 0.008425 | 0.919211 | 0.0057400 | 0.195585714469996 |

| | | | | | | | | | | | |
|---|---|---|---|---|---|---|---|---|---|---|---|
| | 97676 | 802656 | | | | | | | | 2335236268 | |
| **Tyrosine metabolism** | 0.4369058744443 | 0.00917404265803 | Dopamine; 4-(2-Aminoethyl)-1,2-benzenediol; 4-(2-Aminoethyl)benzene-1,2-diol; 3,4-Dihydroxyphenethylamine; 2-(3,4-Dihydroxyphenyl)ethylamine | C03758 | 192.041597841569 | 104.377676781287 | 0.761511 | 0.012665 | 0.932268 | 0.00830391915355844 | 0.195585714469996 |
| **Tyrosine metabolism** | 0.4369058744443 | 0.00917404265803 | Phenylpyruvate; Phenylpyruvic acid; alpha-Ketohydrocinnamic acid; keto-Phenylpyruvate; 3-Phenyl-2-oxopropanoate | C00166 | 165.054695403805 | 116.367243933837 | 0.785605 | 0.012888 | 0.939492 | 0.00819622313896295 | 0.195585714469996 |
| **Tyrosine metabolism** | 0.4369058744443 | 0.00917404265803 | enol-Phenylpyruvate; enol-Phenylpyruvic acid; enol-alpha-Ketohydrocinnamic acid; 2-Hydroxy-3-phenylpropenoate | C02763 | 165.054695403805 | 116.367243933837 | 0.785605 | 0.012888 | 0.939492 | 0.00819622313896295 | 0.195585714469996 |
| **Tyrosine metabolism** | 0.4369058744443 | 0.00917404265803 | 3'-Phosphoadenylyl sulfate; 3'-Phosphoadenosine 5'-phosphosulfate; 3'-Phospho-5'-adenylyl sulfate; PAPS | C00053 | 507.997040927732 | 146.431216399674 | 1.027868 | 0.015748 | 1.048851 | 0.00768108378459198 | 0.195585714469996 |
| **Tyrosine metabolism** | 0.4369058744443 | 0.00917404265803 | Iodine; I2 | C01382 | 256.811618444137 | 92.4428949881603 | 0.75608 | 0.011255 | 0.927992 | 0.0074742425245512 | 0.195585714469996 |
| **Glyoxylate and Dicarboxylate Metabolism** | 0.2263132976466 | 0.0134036375154 | 2-Phospho-D-glycerate; D-Glycerate 2-phosphate | C00631 | 254.986763532727 | 134.097743130349 | 1.061564 | 0.017552 | 1.109685 | 0.00825743494882991 | 0.195585714469996 |
| **Glyoxylate and Dicarboxylate Metabolism** | 0.2263132976466 | 0.0134036375154 | 2-Phosphoglycolate; Phosphoglycolic acid | C00988 | 157.994045674886 | 480.18638863615 | 0.892666 | 0.011903 | 0.969073 | 0.00673392941102124 | 0.195585714469996 |
| **Ubiquinone Biosynthesis** | 0.2774459096122 | 0.0185981569187 | 4-Coumarate; p-Coumaric acid; trans-4-Hydroxycinnamate; trans-p-Hydroxycinnamate; 4-Hydroxycinnamic acid; 4-Hydroxycinnamate | C00811 | 165.054695403805 | 116.367243933837 | 0.785605 | 0.012888 | 0.939492 | 0.00819622313896295 | 0.195585714469996 |
| **Glycine, serine, alanine and threonine metabolism** | 0.5185832986632 | 0.0199044796148 | Choline; Bilineurine | C00114 | 104.107443902613 | 78.7452217926459 | 0.805551 | 0.0102 | 0.941055 | 0.00641288722823619 | 0.195585714469996 |
| **Glycine, serine, alanine and threonine metabolism** | 0.5185832986632 | 0.0199044796148 | L-Methionine; Methionine; L-2-Amino-4methylthiobutyric acid | C00073 | 172.040105107381 | 89.6909060143427 | 0.76934 | 0.012499 | 0.934207 | 0.00812074625284076 | 0.195585714469996 |

| | | | | | | | | | | | |
|---|---|---|---|---|---|---|---|---|---|---|---|
| **Glycine, serine, alanine and threonine metabolism** | 0.518583298632 | 0.0199044796148 | 3-Phospho-D-glycerate; D-Glycerate 3-phosphate; 3-Phospho-(R)-glycerate; 3-Phosphoglycerate | C00197 | 254.986763532727 | 134.097743130349 | 1.061564 | 0.017552 | 1.109685 | 0.00825743494882991 | 0.195585714469996 |
| **Glycine, serine, alanine and threonine metabolism** | 0.518583298632 | 0.0199044796148 | 2-Phosphoglycolate; Phosphoglycolic acid | C00988 | 157.994045674886 | 480.18638863615 | 0.892666 | 0.011903 | 0.969073 | 0.00673392941102124 | 0.195585714469996 |
| **Fatty acid activation** | 0.481162063704 | 0.0199323827058 | (9Z,12Z,15Z)-Octadecatrienoic acid; alpha-Linolenic acid; 9,12,15-Octadecatrienoic acid; Linolenate; alpha-Linolenate | C06427 | 279.231911695023 | 28.1891724631974 | 0.739227 | 0.009696 | 0.919419 | 0.0066300345659458 | 0.195585714469996 |
| **Fatty acid activation** | 0.481162063704 | 0.0199323827058 | (6Z,9Z,12Z)-Octadecatrienoic acid; 6,9,12-Octadecatrienoic acid; gamma-Linolenic acid; Gamolenic acid | C06426 | 279.231911695023 | 28.1891724631974 | 0.739227 | 0.009696 | 0.919419 | 0.0066300345659458 | 0.195585714469996 |
| **Fatty acid activation** | 0.481162063704 | 0.0199323827058 | Tetradecanoic acid; Tetradecanoate; Myristic acid | C06424 | 251.197160068562 | 128.325061638681 | 0.957787 | 0.013649 | 0.988378 | 0.00717150534694222 | 0.195585714469996 |
| **Fatty acid activation** | 0.481162063704 | 0.0199323827058 | Hexadecanoic acid; Hexadecanoate; Hexadecylic acid; Palmitic acid; Palmitate; Cetylic acid | C00249 | 279.231911695023 | 28.1891724631974 | 0.739227 | 0.009696 | 0.919419 | 0.0066300345659458 | 0.195585714469996 |
| **Vitamin E metabolism** | 0.50735458165 | 0.023187298019 | 13'-carboxy-alpha-tocopherol | CE5843 | 461.359536518834 | 41.1319407207359 | 0.739854 | 0.011201 | 0.923038 | 0.00759498232844524 | 0.195585714469996 |
| **Xenobiotics metabolism** | 0.600684633088 | 0.0283528809697 | Bromobenzene-3,4-dihydrodiol; 4-Bromo-3,5-cyclohexadiene-1,2-diol | C14844 | 189.961795729798 | 68.5108382328493 | 0.681336 | 0.010947 | 0.904868 | 0.00803591941502935 | 0.195585714469996 |
| **Xenobiotics metabolism** | 0.600684633088 | 0.0283528809697 | Tetradecanoic acid; Tetradecanoate; Myristic acid | C06424 | 251.197160068562 | 128.325061638681 | 0.957787 | 0.013649 | 0.988378 | 0.00717150534694222 | 0.195585714469996 |
| **Xenobiotics metabolism** | 0.600684633088 | 0.0283528809697 | Bromobenzene-2,3-dihydrodiol; 3-Bromo-3,5-cyclohexadiene-1,2-diol | C14842 | 189.961795729798 | 68.5108382328493 | 0.681336 | 0.010947 | 0.904868 | 0.00803591941502935 | 0.195585714469996 |
| **Xenobiotics metabolism** | 0.600684633088 | 0.0283528809697 | Hexadecanoic acid; Hexadecanoate; Hexadecylic acid; Palmitic acid; Palmitate; Cetylic acid | C00249 | 279.231911695023 | 28.1891724631974 | 0.739227 | 0.009696 | 0.919419 | 0.00663003456 | 0.195585714469996 |

| | | | | | | | | | | | |
|---|---|---|---|---|---|---|---|---|---|---|---|
| | | | | | | | | | | 59458 | |
| **Xenobiotics metabolism** | 0.60068463308 | 0.0283528809697 | 3'-Phosphoadenylyl sulfate; 3'-Phosphoadenosine 5'-phosphosulfate; 3'-Phospho-5'-adenylyl sulfate; PAPS | C00053 | 507.997040927732 | 146.431216399674 | 1.027868 | 0.015748 | 1.048851 | 0.00768108378459198 | 0.195585714469996 |
| **Di-unsaturated fatty acid beta-oxidation** | 0.378251327416 | 0.0325194161802 | trans-Dec-2-enoyl-CoA; (2E)-Decenoyl-CoA | C05275 | 921.24237107895 | 150.430465242413 | 1.028509 | 0.014277 | 1.049735 | 0.00699517578790609 | 0.195585714469996 |
| **Glycosphingolipid biosynthesis - ganglioseries** | 0.378251327416 | 0.0325194161802 | 3'-Phosphoadenylyl sulfate; 3'-Phosphoadenosine 5'-phosphosulfate; 3'-Phospho-5'-adenylyl sulfate; PAPS | C00053 | 507.997040927732 | 146.431216399674 | 1.027868 | 0.015748 | 1.048851 | 0.00768108378459198 | 0.195585714469996 |
| **Phosphatidylinositol phosphate metabolism** | 0.673947532305 | 0.0821391815902 | Hexadecanoic acid; Hexadecanoate; Hexadecylic acid; Palmitic acid; Palmitate; Cetylic acid | C00249 | 279.231911695023 | 28.1891724631974 | 0.739227 | 0.009696 | 0.919419 | 0.0066300345659458 | 0.195585714469996 |
| **Phosphatidylinositol phosphate metabolism** | 0.673947532305 | 0.0821391815902 | D-myo-Inositol 1,2-cyclic phosphate; 1D-myo-Inositol 1,2-cyclic phosphate | C04299 | 133.006644128167 | 64.7740819337371 | 0.74598 | 0.008872 | 0.919582 | 0.00604399918616394 | 0.195585714469996 |
| **Omega-3 fatty acid metabolism** | 0.633373940637 | 0.101234111137 | (9Z,12Z,15Z)-Octadecatrienoic acid; alpha-Linolenic acid; 9,12,15-Octadecatrienoic acid; Linolenate; alpha-Linolenate | C06427 | 279.231911695023 | 28.1891724631974 | 0.739227 | 0.009696 | 0.919419 | 0.0066300345659458 | 0.195585714469996 |
| **Glycolysis and Gluconeogenesis** | 0.764250008433 | 0.106764598315 | 2-Phospho-D-glycerate; D-Glycerate 2-phosphate | C00631 | 254.986763532727 | 134.097743130349 | 1.061564 | 0.017552 | 1.109685 | 0.00825743494882991 | 0.195585714469996 |
| **Glycolysis and Gluconeogenesis** | 0.764250008433 | 0.106764598315 | 3-Phospho-D-glycerate; D-Glycerate 3-phosphate; 3-Phospho-(R)-glycerate; 3-Phosphoglycerate | C00197 | 254.986763532727 | 134.097743130349 | 1.061564 | 0.017552 | 1.109685 | 0.00825743494882991 | 0.195585714469996 |
| **Glycolysis and Gluconeogenesis** | 0.764250008433 | 0.106764598315 | Carbamoyl phosphate | C00169 | 140.982056004229 | 310.601686958314 | 0.759972 | 0.009107 | 0.924639 | 0.00608744857666477 | 0.195585714469996 |
| **Porphyrin metabolism** | 0.745836911495 | 0.115588027425 | Uroporphyrinogen I | C05766 | 921.24237107895 | 150.430465242413 | 1.028509 | 0.014277 | 1.049735 | 0.00699517578790609 | 0.195585714469996 |

| | | | | | | | | | | | |
|---|---|---|---|---|---|---|---|---|---|---|---|
| **Porphyrin metabolism** | 0.745836911495 | 0.115588027425 | Hydroxymethylbilane | C01024 | 877.275430523807 | 145.450729868039 | 1.025287 | 0.017436 | 1.04453 | 0.00848137283612593 | 0.195585714469996 |
| **Porphyrin metabolism** | 0.745836911495 | 0.115588027425 | Uroporphyrinogen III | C01051 | 921.24237107895 | 150.430465242413 | 1.028509 | 0.014277 | 1.049735 | 0.00699517578790609 | 0.195585714469996 |
| **Aspartate and asparagine metabolism** | 0.822873368732 | 0.123478035657 | 4-Aminobutanal; 4-Aminobutyraldehyde; Butyraldehyde, 4-amino- | C00555 | 89.0795646241381 | 42.0813089452326 | 0.748234 | 0.008425 | 0.919211 | 0.00574002335236268 | 0.195585714469996 |
| **Aspartate and asparagine metabolism** | 0.822873368732 | 0.123478035657 | Carbamoyl phosphate | C00169 | 140.982056004229 | 310.601686958314 | 0.759972 | 0.009107 | 0.924639 | 0.00608744857666477 | 0.195585714469996 |
| **Galactose metabolism** | 0.821437186772 | 0.166638916907 | D-Tagatose 1-phosphate | tag1p-D | 327.007830935988 | 550.437193098651 | 0.758058 | 0.009717 | 0.925383 | 0.00648738452413733 | 0.195585714469996 |
| **Omega-6 fatty acid metabolism** | 0.777441728451 | 0.179119678411 | trans-Dec-2-enoyl-CoA; (2E)-Decenoyl-CoA | C05275 | 921.24237107895 | 150.430465242413 | 1.028509 | 0.014277 | 1.049735 | 0.00699517578790609 | 0.195585714469996 |
| **Valine, leucine and isoleucine degradation** | 0.866429659058 | 0.185929680637 | L-Isoleucine; 2-Amino-3-methylvaleric acid | C00407 | 133.105291985925 | 110.034556236308 | 0.793317 | 0.013783 | 0.94301 | 0.00865567606652606 | 0.195585714469996 |
| **Valine, leucine and isoleucine degradation** | 0.866429659058 | 0.185929680637 | L-Leucine; 2-Amino-4-methylvaleric acid; (2S)-alpha-2-Amino-4-methylvaleric acid; (2S)-alpha-Leucine | C00123 | 133.105291985925 | 110.034556236308 | 0.793317 | 0.013783 | 0.94301 | 0.00865567606652606 | 0.195585714469996 |
| **Methionine and cysteine metabolism** | 0.877405448922 | 0.198255230658 | L-Methionine; Methionine; L-2-Amino-4methylthiobutyric acid | C00073 | 172.040105107381 | 89.6909060143427 | 0.76934 | 0.012499 | 0.934207 | 0.00812074625284076 | 0.195585714469996 |
| **Methionine and cysteine metabolism** | 0.877405448922 | 0.198255230658 | 3'-Phosphoadenylyl sulfate; 3'-Phosphoadenosine 5'-phosphosulfate; 3'-Phospho-5'-adenylyl sulfate; PAPS | C00053 | 507.997040927732 | 146.431216399674 | 1.027868 | 0.015748 | 1.048851 | 0.00768108378459198 | 0.195585714469996 |
| **Arginine and Proline Metabolism** | 0.877078363815 | 0.221999947583 | 4-Aminobutanal; 4-Aminobutyraldehyde; Butyraldehyde, 4-amino- | C00555 | 89.0795646241381 | 42.0813089452326 | 0.748234 | 0.008425 | 0.919211 | 0.00574002335 | 0.195585714469996 |

| | | | | | | | | | | | |
|---|---|---|---|---|---|---|---|---|---|---|---|
| | | | | | | | | | | 236268 | |
| **Arginine and Proline Metabolism** | 0.87707836381 5 | 0.221999947583 | L-Methionine; Methionine; L-2-Amino-4methylthiobutyric acid | C00073 | 172.040105107381 | 89.6909060143427 | 0.76934 | 0.012499 | 0.934207 | 0.0081207462528407 6 | 0.195585714469996 |
| **Fructose and mannose metabolism** | 0.85454053772 3 | 0.246018290225 | fructose 3-phosphate | CE3074 | 327.007830935988 | 550.437193098651 | 0.758058 | 0.009717 | 0.925383 | 0.0064873845241373 3 | 0.195585714469996 |
| **Vitamin A (retinol) metabolism** | 0.85454053772 3 | 0.246018290225 | 11-cis-Retinyl palmitate | C03455 | 263.236750689057 | 32.3161891554784 | 0.763471 | 0.012306 | 0.932175 | 0.0080603897999465 | 0.195585714469996 |
| **Arachidonic acid metabolism** | 0.92134428956 1 | 0.26157551043 | (9Z,12Z,15Z)-Octadecatrienoic acid; alpha-Linolenic acid; 9,12,15-Octadecatrienoic acid; Linolenate; alpha-Linolenate | C06427 | 279.231911695023 | 28.1891724631974 | 0.739227 | 0.009696 | 0.919419 | 0.0066300345659458 | 0.195585714469996 |
| **Pyrimidine metabolism** | 0.92475584473 | 0.292941294268 | Carbamoyl phosphate | C00169 | 140.982056004229 | 310.601686958314 | 0.759972 | 0.009107 | 0.924639 | 0.0060874485766647 7 | 0.195585714469996 |
| **Butanoate metabolism** | 0.91656492778 5 | 0.328345356392 | (S)-2-Aceto-2-hydroxybutanoate; (S)-2-Hydroxy-2-ethyl-3-oxobutanoate | C06006 | 149.059827417723 | 587.065552074052 | 0.733327 | 0.010852 | 0.920298 | 0.0074325162881169 2 | 0.195585714469996 |
| **Pentose phosphate pathway** | 0.93360903999 8 | 0.359900575187 | D-Glucono-1,5-lactone 6-phosphate; 6-Phospho-D-glucono-1,5-lactone | C01236 | 327.007830935988 | 550.437193098651 | 0.758058 | 0.009717 | 0.925383 | 0.0064873845241373 3 | 0.195585714469996 |
| **Squalene and cholesterol biosynthesis** | 0.94733241756 4 | 0.390422353572 | Episterol | C15777 | 200.183786431786 | 30.1644756714824 | 0.76105 | 0.008147 | 0.922758 | 0.0054729674110436 4 | 0.195585714469996 |
| **Squalene and cholesterol biosynthesis** | 0.94733241756 4 | 0.390422353572 | 24-Methylenecholesterol | C15781 | 200.183786431786 | 30.1644756714824 | 0.76105 | 0.008147 | 0.922758 | 0.0054729674110436 4 | 0.195585714469996 |
| **Androgen and estrogen biosynthesis and metabolism** | 0.98779638849 3 | 0.510935384898 | 3'-Phosphoadenylyl sulfate; 3'-Phosphoadenosine 5'-phosphosulfate; 3'-Phospho-5'-adenylyl sulfate; PAPS | C00053 | 507.997040927732 | 146.431216399674 | 1.027868 | 0.015748 | 1.048851 | 0.0076810837845919 8 | 0.195585714469996 |

| | | | | | | | | | | | |
|---|---|---|---|---|---|---|---|---|---|---|---|
| **Purine metabolism** | 0.9875387510228 | 0.549295313914 | Urate; Uric acid | C00366 | 252.997269313604 | 17.6542994847686 | 0.740519 | 0.01058 | 0.921865 | 0.00718885725433693 | 0.195585714469996 |
| **C21-steroid hormone biosynthesis and metabolism** | 0.997864412275 | 0.655533608793 | 3'-Phosphoadenylyl sulfate; 3'-Phosphoadenosine 5'-phosphosulfate; 3'-Phospho-5'-adenylyl sulfate; PAPS | C00053 | 507.997040927732 | 146.431216399674 | 1.027868 | 0.015748 | 1.048851 | 0.00768108378459198 | 0.195585714469996 |
| **Selenoamino acid metabolism** | 0.918621397261 | 0.999986263096 | Selenomethionine | C05335 | 235.958052221287 | 58.4177818496412 | 1.029648 | 0.011337 | 1.051162 | 0.00562018247530482 | 0.195585714469996 |
| **Glutamate metabolism** | 0.783338757398 | 0.999986263096 | Carbamoyl phosphate | C00169 | 140.982056004229 | 310.601686958314 | 0.759972 | 0.009107 | 0.924639 | 0.00608744857666477 | 0.195585714469996 |
| **Proteoglycan biosynthesis** | 0.500317023052 | 0.999986263096 | 3'-Phosphoadenylyl sulfate; 3'-Phosphoadenosine 5'-phosphosulfate; 3'-Phospho-5'-adenylyl sulfate; PAPS | C00053 | 507.997040927732 | 146.431216399674 | 1.027868 | 0.015748 | 1.048851 | 0.00768108378459198 | 0.195585714469996 |
| **Keratan sulfate biosynthesis** | 0.425834611149 | 0.999986263096 | 3'-Phosphoadenylyl sulfate; 3'-Phosphoadenosine 5'-phosphosulfate; 3'-Phospho-5'-adenylyl sulfate; PAPS | C00053 | 507.997040927732 | 146.431216399674 | 1.027868 | 0.015748 | 1.048851 | 0.00768108378459198 | 0.195585714469996 |
| **Parathio degradation** | 0.340311783887 | 0.999986263096 | Diethylthiophosphoric acid; DETP | C06607 | 254.986763532727 | 134.097743130349 | 1.061564 | 0.017552 | 1.109685 | 0.00825743494882991 | 0.195585714469996 |
| **Linoleate metabolism** | 0.00310216651967 | 0.000353032634069 | 13-OxoODE; 13-KODE; (9Z,11E)-13-Oxooctadeca-9,11-dienoic acid | C14765 | 295.226406338711 | 29.037835348842 | 0.70858 | 0.003996 | 0.88962 | 0.00300201834681391 | 0.19290463114884 |
| **De novo fatty acid biosynthesis** | 0.104709729668 | 0.00101148958738 | Dodecanoic acid; Dodecanoate; Dodecylcarboxylate; Lauric acid | C02679 | 269.174558814739 | 17.3993398078578 | 0.962523 | 0.017933 | 0.990613 | 0.00925132273475404 | 0.196468734356961 |
| **Xenobiotics metabolism** | 0.600684633088 | 0.0283528809697 | Trichloroacetate | C11150 | 164.908517444309 | 80.6034797885681 | 0.772539 | 0.014428 | 0.938322 | 0.00927274520414212 | 0.196468734356961 |
| **Xenobiotics metabolism** | 0.600684633088 | 0.0283528809697 | Dodecanoic acid; Dodecanoate; Dodecylcarboxylate; Lauric acid | C02679 | 269.174558814739 | 17.3993398078578 | 0.962523 | 0.017933 | 0.990613 | 0.00925132273 | 0.196468734356961 |

| | | | | | | | | | | | |
|---|---|---|---|---|---|---|---|---|---|---|---|
| | | | | | | | | | | 475404 | |
| **Linoleate metabolism** | 0.00310216651967 | 0.000353032634069 | | CE5526 | 295.226406338711 | 29.037835348842 | 0.70858 | 0.003996 | 0.88962 | 0.00300201834681391 | 0.19290463114884 |
| **Linoleate metabolism** | 0.00310216651967 | 0.000353032634069 | (6Z,9Z,12Z)-Octadecatrienoic acid; 6,9,12-Octadecatrienoic acid; gamma-Linolenic acid; Gamolenic acid | C06426 | 279.231911695023 | 28.1891724631974 | 0.739227 | 0.009696 | 0.919419 | 0.0066300345659458 | 0.195585714469996 |
| **Linoleate metabolism** | 0.00310216651967 | 0.000353032634069 | 13(S)-HODE; (13S)-Hydroxyoctadecadienoic acid; (9Z, 11E)-(13S)-13-Hydroxyoctadeca-9,11-dienoic acid | C14762 | 381.20362781455 | 570.884206840032 | 1.052803 | 0.020216 | 1.094562 | 0.00952119613406099 | 0.196956027492531 |
| **Linoleate metabolism** | 0.00310216651967 | 0.000353032634069 | 12(13)-EpOME; (12R,13S)-(9Z)-12,13-Epoxyoctadecenoic acid | C14826 | 381.20362781455 | 570.884206840032 | 1.052803 | 0.020216 | 1.094562 | 0.00952119613406099 | 0.196956027492531 |
| **Drug metabolism - cytochrome P450** | 0.339766036825 | 0.00657261942317 | Felbamate | C07501 | 277.060662501404 | 76.67790354067 | 0.786771 | 0.01531 | 0.943442 | 0.00964291950781193 | 0.196956027492531 |
| **Glycine, serine, alanine and threonine metabolism** | 0.518583298632 | 0.0199044796148 | Betaine; Trimethylaminoacetate; Glycine betaine; N,N,N-Trimethylglycine; Trimethylammonioacetate | C00719 | 156.041774564348 | 114.690419565258 | 0.824315 | 0.015957 | 0.954086 | 0.00959473366412922 | 0.196956027492531 |
| **Xenobiotics metabolism** | 0.600684633088 | 0.0283528809697 | | C11149 | 150.931312242014 | 68.4111685415314 | 0.758375 | 0.014552 | 0.934712 | 0.00951450398755016 | 0.196956027492531 |
| **Valine, leucine and isoleucine degradation** | 0.866429659058 | 0.185929680637 | L-Valine; 2-Amino-3-methylbutyric acid | C00183 | 156.041774564348 | 114.690419565258 | 0.824315 | 0.015957 | 0.954086 | 0.00959473366412922 | 0.196956027492531 |
| **Butanoate metabolism** | 0.916564927785 | 0.328345356392 | 5-Aminopentanoate; 5-Aminopentanoic acid; 5-Aminovaleric acid | C00431 | 156.041774564348 | 114.690419565258 | 0.824315 | 0.015957 | 0.954086 | 0.00959473366412922 | 0.196956027492531 |
| **Tryptophan metabolism** | 0.991312719246 | 0.543477262998 | Indole-3-acetate; Indole-3-acetic acid; (Indol-3-yl)acetate; Indoleacetate; Indoleacetic acid | C00954 | 177.074479811589 | 74.1646177807957 | 0.763654 | 0.014538 | 0.936106 | 0.00944340488621853 | 0.196956027492531 |

| | | | | | | | | | | | |
|---|---|---|---|---|---|---|---|---|---|---|---|
| **Tryptophan metabolism** | 0.991312719246 | 0.543477262998 | 5-Hydroxyindoleacetaldehyde | C05634 | 177.074479811589 | 74.1646177807957 | 0.763654 | 0.014538 | 0.936106 | 0.00944340488621853 | 0.196956027492531 |
| **C21-steroid hormone biosynthesis and metabolism** | 0.997864412275 | 0.655533608793 | 18-Hydroxycortisol | CE1342 | 381.20362781455 | 570.884206840032 | 1.052803 | 0.020216 | 1.094562 | 0.00952119613406099 | 0.196956027492531 |
| **Hexose phosphorylation** | 0.0877751590204 | 0.00117065111852 | D-Glucosamine; Chitosamine; 2-Amino-2-deoxy-D-glucose | C00329 | 182.081149725003 | 115.684673000946 | 0.804881 | 0.016162 | 0.949294 | 0.00993632002554234 | 0.199069197631384 |
| **Tyrosine metabolism** | 0.436905874443 | 0.00917404265803 | | CE2174 | 182.081149725003 | 115.684673000946 | 0.804881 | 0.016162 | 0.949294 | 0.00993632002554234 | 0.199069197631384 |
| **Tyrosine metabolism** | 0.436905874443 | 0.00917404265803 | L-Tyrosine; (S)-3-(p-Hydroxyphenyl)alanine; (S)-2-Amino-3-(p-hydroxyphenyl)propionic acid; Tyrosine | C00082 | 182.081149725003 | 115.684673000946 | 0.804881 | 0.016162 | 0.949294 | 0.00993632002554234 | 0.199069197631384 |
| **Biopterin metabolism** | 0.516758295442 | 0.0622758094373 | L-Tyrosine; (S)-3-(p-Hydroxyphenyl)alanine; (S)-2-Amino-3-(p-hydroxyphenyl)propionic acid; Tyrosine | C00082 | 182.081149725003 | 115.684673000946 | 0.804881 | 0.016162 | 0.949294 | 0.00993632002554234 | 0.199069197631384 |
| **Aminosugars metabolism** | 0.745836911495 | 0.115588027425 | D-Glucosamine; Chitosamine; 2-Amino-2-deoxy-D-glucose | C00329 | 182.081149725003 | 115.684673000946 | 0.804881 | 0.016162 | 0.949294 | 0.00993632002554234 | 0.199069197631384 |
| **Linoleate metabolism** | 0.003102166651967 | 0.000353032634069 | 9(10)-EpOME; (9R,10S)-(12Z)-9,10-Epoxyoctadecenoic acid | C14825 | 381.20362781455 | 570.884206840032 | 1.052803 | 0.020216 | 1.094562 | 0.00952119613406099 | 0.196956027492531 |
| **Carnitine shuttle** | 0.0831103651174 | 0.000874817459984 | Hexadecenoyl carnitine | hdcecrn | 398.325066701968 | 94.4213470768914 | 0.908373 | 0.018383 | 0.977296 | 0.0100105522166375 | 0.199226826059028 |
| **Carnitine shuttle** | 0.0831103651174 | 0.000874817459984 | trans-Hexadec-2-enoyl carnitine | hdd2crn | 398.325066701968 | 94.4213470768914 | 0.908373 | 0.018383 | 0.977296 | 0.0100105522166375 | 0.199226826059028 |
| **Lysine metabolism** | 0.125453555247 | 0.0014816264063 | 4-Trimethylammoniobutanoate | C01181 | 146.117576611507 | 38.6922760654623 | 0.77875 | 0.017016 | 0.943782 | 0.01077169835 | 0.199226826059028 |

| | | | | | | | | | | | |
|---|---|---|---|---|---|---|---|---|---|---|---|
| | | | | | | | | | | 20912 | |
| **Drug metabolism - cytochrome P450** | 0.339766036825 | 0.00657261942317 | Monoethylglycinexylidide | C16561 | 115.06978393009 | 75.5064090720009 | 0.770699 | 0.015567 | 0.939609 | 0.00999203846008179 | 0.199226826059028 |
| **Vitamin E metabolism** | 0.50735458165 | 0.023187298019 | 11'-carboxy-gama-chromanol | CE5718 | 203.150268805149 | 122.685989501057 | 0.725071 | 0.015459 | 0.927581 | 0.0105214882985428 | 0.199226826059028 |
| **Xenobiotics metabolism** | 0.600684633088 | 0.0283528809697 | Glutathione episulfonium ion | C14874 | 373.071419395144 | 137.328446330345 | 1.031412 | 0.022283 | 1.056187 | 0.0106556696994028 | 0.199226826059028 |
| **Omega-3 fatty acid metabolism** | 0.633373940637 | 0.101234111137 | Stearidonic acid; 6,9,12,15-Octadecatetraenoic acid; (6Z,9Z,12Z,15Z)-Octadecatetraenoic acid | C16300 | 277.215774306569 | 49.8184075751079 | 0.963211 | 0.020265 | 0.991225 | 0.0103890349518803 | 0.199226826059028 |
| **Squalene and cholesterol biosynthesis** | 0.947332417564 | 0.390422353572 | (R)-5-Diphosphomevalonate | C01143 | 311.007680447161 | 81.7732800696542 | 0.755917 | 0.015938 | 0.936373 | 0.0104104703692219 | 0.199226826059028 |
| **Vitamin A (retinol) metabolism** | 0.854540537723 | 0.246018290225 | all-trans-Retinoyl-beta-glucuronide; Retinoyl glucuronide | C11061 | 250.119279796206 | 35.2890526486596 | 0.786572 | 0.017228 | 0.946022 | 0.0107963678146177 | 0.19931603301876 |
| **Tyrosine metabolism** | 0.436905874443 | 0.00917404265803 | Epinine; Deoxyepinephrine | C07453 | 170.096404303439 | 44.2442279559255 | 0.778149 | 0.017231 | 0.943931 | 0.0109098185866109 | 0.199576153124468 |
| **Tyrosine metabolism** | 0.436905874443 | 0.00917404265803 | 3-Methoxytyramine | C05587 | 170.096404303439 | 44.2442279559255 | 0.778149 | 0.017231 | 0.943931 | 0.0109098185866109 | 0.199576153124468 |
| **Vitamin E metabolism** | 0.50735458165 | 0.023187298019 | 13'-hydroxy-alpha-tocotrienol | CE7144 | 479.292540202803 | 577.055640982851 | 0.80657 | 0.018031 | 0.95195 | 0.011009532429766 | 0.199784667796754 |
| **Prostaglandin formation from arachidonate** | 0.973563502867 | 0.427665732887 | 11-hydroxyeicosatetraenoate glyceryl ester | CE5707 | 395.276751604426 | 116.938437568338 | 0.952748 | 0.021392 | 0.988994 | 0.0110554257903214 | 0.199784667796754 |
| **Drug metabolism - other enzymes** | 0.7774417 | 0.179119678411 | 6-Methylmercaptopurine | C16614 | 251.000089412883 | 17.5624142347672 | 0.845059 | 0.18047 | 1.035917 | 0.1051539 | 0.46574175598885 |

| | | | | | | | | | | |
|---|---|---|---|---|---|---|---|---|---|---|
| | 2845<br>1 | | | | | | | | | 2450<br>3033 |
